# Structure and RNA template requirements of Arabidopsis RNA-DEPENDENT RNA POLYMERASE 2

**DOI:** 10.1101/2021.08.31.458351

**Authors:** Akihito Fukudome, Jasleen Singh, Vibhor Mishra, Eswar Reddem, Francisco Martinez-Marquez, Sabine Wenzel, Rui Yan, Momoko Shiozaki, Zhiheng Yu, Joseph Che-Yen Wang, Yuichiro Takagi, Craig S. Pikaard

## Abstract

RNA-dependent RNA polymerases play essential roles in RNA-mediated gene silencing in eukaryotes. In *Arabidopsis*, RNA-DEPENDENT RNA POLYMERASE 2 (RDR2) physically interacts with DNA-dependent NUCLEAR RNA POLYMERASE IV (Pol IV) and their activities are tightly coupled, with Pol IV transcriptional arrest or termination, involving the nontemplate DNA strand, somehow enabling RDR2 to engage Pol IV transcripts and generate double-stranded RNAs. The dsRNAs are then released from the Pol IV-RDR2 complex and diced into siRNAs that guide RNA-directed DNA methylation and silencing. Here we report the structure of full-length RDR2, at an overall resolution of 3.1 Å, determined by cryo-electron microscopy. The N-terminal region contains an RNA-recognition motif (RRM) adjacent to a positively charged channel that leads to a catalytic center with striking structural homology to the catalytic centers of multisubunit DNA-dependent RNA polymerases. We show that RDR2 initiates 1-2 nucleotides (nt) internal to the 3’ ends of its templates and can transcribe the RNA of an RNA-DNA hybrid provided that 9 or more nucleotides at the RNA’s 3’ end is unpaired. Using a nucleic acid configuration that mimics the arrangement of RNA and DNA strands upon Pol IV transcriptional arrest, we show that displacement of the RNA 3’ end occurs as the DNA template and non-template strands reanneal, enabling RDR2 transcription. These results suggest a model in which Pol IV arrest and backtracking displaces the RNA 3’ end as the DNA strands reanneal, allowing RDR2 to engage the RNA and transcribe the second strand.

**Significance:** RDR2 is critical for siRNA-directed DNA methylation in Arabidopsis, functioning in physical association with DNA-dependent Pol IV to synthesize the second strands of double-stranded siRNA precursors. Basepairing between the DNA template strand transcribed by Pol IV and the nontemplate DNA strand is known to induce Pol IV arrest and Pol IV-RDR2 transcriptional coupling, but how this occurs is unknown. We report the structure of RDR2 and experimental evidence for how RDR2 engages its RNA templates and initiates transcription. RDR2 engages the ends of RNAs displaced from RNA-DNA hybrids, suggesting a model in which Pol IV arrest and backtracking, accompanied by DNA strand reannealing, extrudes the 3’ end of the Pol IV transcript, allowing RNA engagement and second-strand synthesis.

## Introduction

RNA-dependent RNA polymerases (RDRs) are encoded within the genomes of many eukaryotes and function in gene silencing by converting single-stranded RNAs into double-stranded (ds) precursors of short-interfering RNAs (siRNAs) (1, 2). The siRNAs associate with Argonaute family proteins and basepair with target locus RNAs to interfere with their translation or bring about transcriptional silencing at the chromosomal loci that encode them (3-5).

In plants, transcriptional silencing via siRNA-directed DNA methylation (RdDM) requires RNA-DEPENDENT RNA POLYMERASE 2 (RDR2), which associates with DNA-dependent NUCLEAR RNA POLYMERASE IV (Pol IV) (6, 7) via direct physical interaction (8). The coupled transcription reactions of the Pol IV-RDR2 complex yield short dsRNAs of ∼30 bp (9, 10) that are then diced into 24 or 23 nt siRNAs by DICER-LIKE3 (DCL3) (9, 11-13). These siRNAs are then loaded into an Argonaute protein, primarily AGO4 (12) and guide the resulting complexes to target loci via basepairing interactions with non-coding transcripts synthesized by NUCLEAR RNA POLYMERASE V (Pol V) (14, 15). Subsequent recruitment of DNA methyltransferase DRM2 and histone modifying enzymes alters the local chromatin environment, inhibiting promoter-dependent transcription by RNA polymerases I, II, or III (15-19). In this manner, thousands of loci, mostly encoding transposable elements, are transcriptionally silenced throughout the genome. Analogous RDR-dependent transcriptional silencing pathways repress transposable elements in fission yeast, Neurospora, and nematodes (20-24).

Important mechanistic insights into the Pol IV-RDR2 partnership have come from recapitulation of siRNA biogenesis *in vitro* (11). Although Pol IV and RDR2 stably associate, they do not produce dsRNA when provided with only a single-stranded DNA oligonucleotide template. Instead, single-stranded Pol IV transcripts are generated, which remain associated with the template as persistent RNA-DNA hybrids (6, 11). However, if a non-template DNA oligonucleotide is annealed to the template DNA strand, Pol IV arrests after transcribing ∼12-16 nt into the basepaired DNA region and RDR2 transcription now occurs, yielding dsRNAs that are released from the Pol IV-RDR2 complex (11). How basepairing between the DNA template and non-template strands brings about Pol IV arrest and RDR2 coupling remains unclear.

Structural information for eukaryotic cellular RNA-dependent RNA polymerases is currently limited to crystal structures for two fungal QDE-1 (Quelling Defective -1) enzymes. One is a partial structure for the *Neurospora crassa* enzyme, missing amino acids 1-376, and the other is a partial structure for *Thermothielavioides terrestris* QDE-1 missing amino acids 1-363 (25, 26). We have determined the full-length structure of Arabidopsis RDR2 at 3.1Å overall resolution by single-particle cryo-electron microscopy. Striking structural similarities are apparent between the active centers of RDR2, the QDE-1 enzymes and multi-subunit DNA-dependent RNA polymerases. The N-terminal region of RDR2 includes an RNA recognition motif (RRM) proposed to help define the path of RNA within the enzyme. Using oligonucleotides that recapitulate the configuration of the DNA template, DNA nontemplate and RNA strands upon Pol IV arrest, we show that template - nontemplate DNA strand reannealing displaces the RNA 3’ end from the template DNA, making it available for RDR2 engagement. Because RNA strand displacement and extrusion also coincides with template-nontemplate DNA strand reannealing when multisubunit RNA polymerases undergo backtracking, we propose that Pol IV arrest and backtracking is the likely mechanism by which Pol IV transcripts are channeled to RDR2.

## Results

### Expression and single-particle cryoEM of recombinant RDR2

Full-length RDR2 expressed in insect cells using a baculovirus expression vector system (Fig. 1A and 1B) was subjected to single-particle imaging using a Titan Krios instrument equipped with a Gatan K3 detector. Images were collected with no tilt of the specimen grid or with 30° tilt. Following 2D classification, particle images from both collection methods were combined and used to generate an initial model. Successive rounds of 3D classification, 3D refinement, CTF refinement and particle polishing were then conducted, resulting in a map with 3.1 Å overall resolution using RELION 3.1 (27-29) (Fig. S1, S2, S3 and S4). Processing of the same datasets using cryoSPARC v3.2.0 (30) resulted in a very similar map with 3.57 Å overall resolution (Fig. S5, S6 and S7). The higher resolution RELION map was then used for model building, employing Buccaneer (31) and manual efforts. Structures of other RNA-dependent RNA polymerase were not used for RDR2 model building. However, the protein structure predictor I-TASSER (32) was used to aid model building for the amino terminal end of RDR2 (aa 1-100) due to relatively low quality density for this region. Details of the model building process are provided as Supplemental Information.

**Figure 1.**
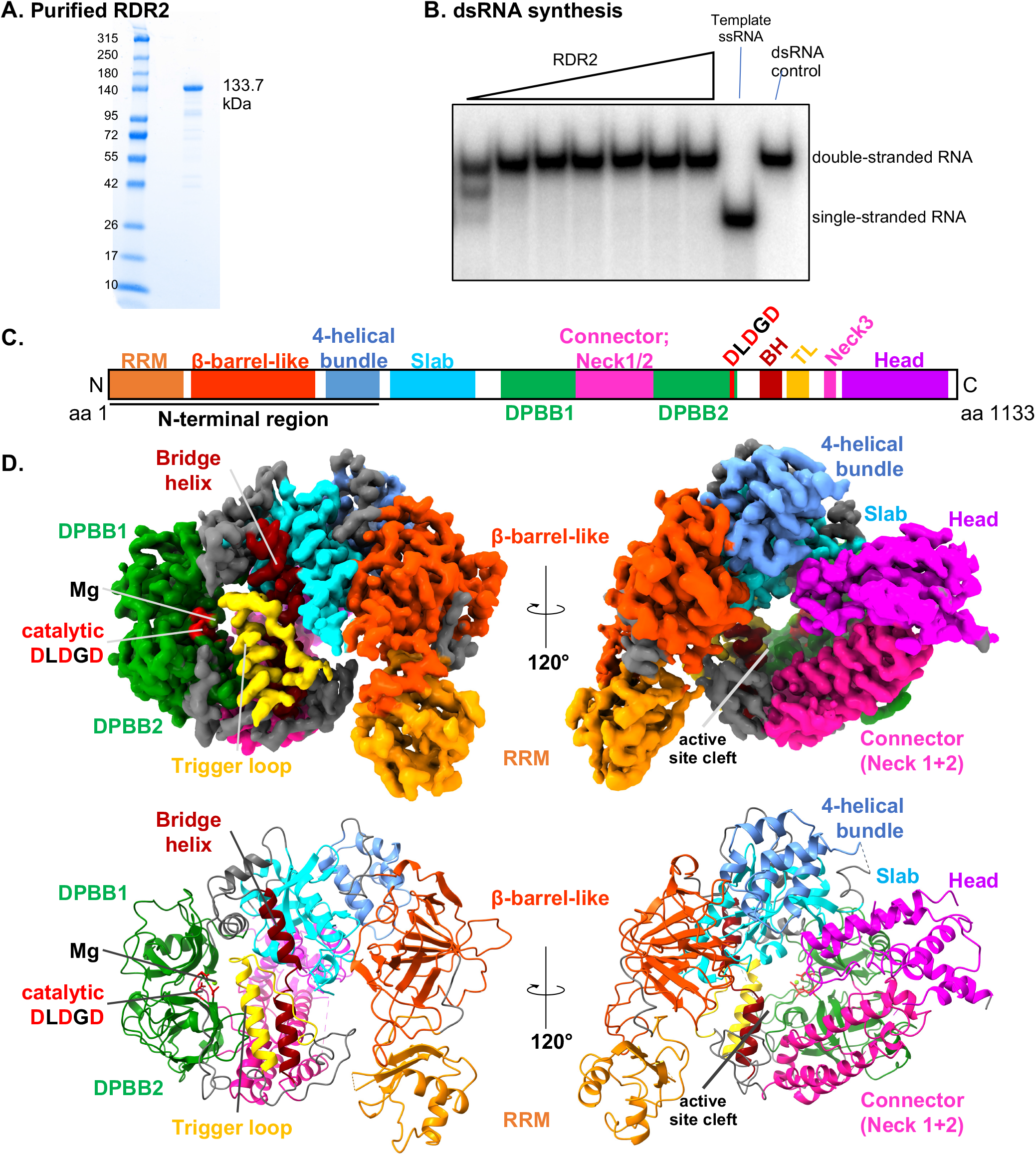
Structure of RDR2. (A) Coomassie Brilliant Blue-stained SDS-PAGE gel showing recombinant RDR2 used in this study. (B) dsRNA synthesis by recombinant RDR2. RDR2 (0.11 to 3.6 μM) was incubated with single-stranded RNA and NTPs. Products were resolved by 15% native PAGE. (C) Domain structure of RDR2: RRM (RNA-recognition motif) shown in orange; β-barrel-like (dark orange); 4-Helical bundle (cornflower blue); Slab (blue); DPBB: Double-psi β barrel (green); Connector/Neck helices (magenta); BH:Bridge-Helix (brown); TL:Trigger-Loop (yellow); and Head (purple). The catalytic site (Metal A site), DLDGD is shown in red. (D) Ribbon diagram of RDR2 structure. Domains are colored as in panel C.

### Overall structure of RDR2

The N-terminal region (aa 1-360) of RDR2 contains three domains: an N-terminal domain (NTD: residues 1-100) that includes an RNA recognition motif (RRM), a beta-barrel-like domain and a 4-helical bundle domain (Fig. 1C and 1D). Following the N-terminal region is an active site cleft composed of 7 domains termed Slab, DPBB1, connector helices Neck 1 and Neck 2, DPBB2, Bridge Helix, Trigger Loop, connector helix Neck3, and Head, respectively (Fig. 1C) (26, 33-35). The two DPBBs are linked by a connector domain, similar to Neurospora QDE-1ΔN and phi14:2 phage polymerase (35), which extends from DPBB1 and includes a short α helix and part of the Neck 1 helix, resembling the “anchor” element that leads into the Pol II clamp domain in the yeast Pol II subunit, RPB2 (Fig. S8). The three highly conserved aspartic acids (Asp triad) required for phosphodiester bond catalysis are located within a loop of DPBB2.

Following DPBB2, a loop passes across a potential secondary channel and connects to the bridge helix domain, followed by the trigger loop. Neck helix 3, part of which was not modeled due to flexibility in the loop, connects to the C-terminal Head domain, composed of two long α-helices and a loop (again, not fully modeled due to flexibility) that protrudes into the cleft. Multiple structural features typical of viral RDRs, namely Thumb, Palm, or Finger domains, are not seen in the RDR2 structure.

### Structural conservation within RDR2 and multi-subunit RNAP catalytic centers

A superimposition of the RDR2 active site loop (aa 828-836) of DPBB2 onto the corresponding structures of *Saccharomyces cerevisiae* Pol II and bacterial (*Thermus thermophilus)* RNAP is shown in Fig. 2A. The RDR2 and yeast Pol II active site loops share similarity to almost the same degree (RMSD = 0.649 Å) as the similarity between bacterial RNAP and yeast Pol II (RMSD = 0.431 Å). Other domains of the catalytic core, including the two DPBBs, bridge helix and trigger loop also align when superimposed. The Asp-triad of RDR2 and the multi-subunit RNAPs align nearly perfectly (Fig. 2B). The Asp triad of multi-subunit RNAPs corresponds to the Metal A site, one of two sites that bind and coordinate magnesium ions at the site of catalysis (33). Extra density at the position of the three Asp residues is seen in the RDR2 EM map, likely corresponding to a Mg^2+^ ion, consistent with this being the Metal A site (Fig. 2B). No density corresponding to a metal B site is observed in the RDR2 map. Metal B ions were also not observed in QDE-1ΔN crystal structures. At the position corresponding to Glu-836 in yeast Pol II, which helps coordinate metal B, a glycine (Gly-556) is present in RDR2 as well as in other RDRs This is apparent upon alignment of eleven members of the alpha clade (RDRα) and five members of the gamma clade (RDRγ) of RDRs (Fig. S9) (2).

**Figure 2.**
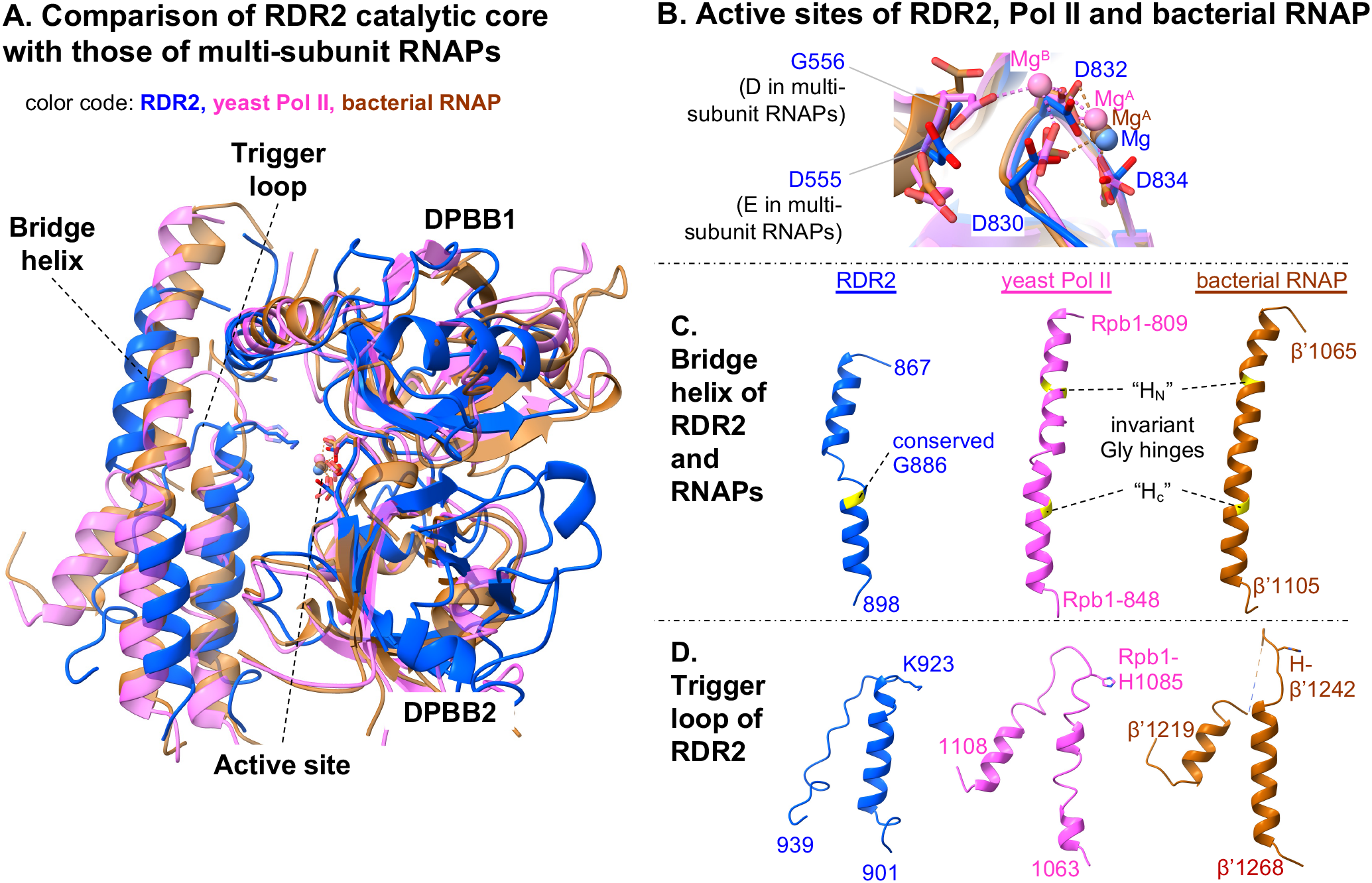
Structural comparisons of RDR2 with multi-subunit RNAPs. (A) Superimposition of the structure of the RDR2 catalytic core and those of Saccharomyces *cerevisiae* Pol II (PDB: 2E2H) and bacterial *Thermus thermophilus* RNAP (PDB: 4Q4Z). RDR2 colored in blue, yeast Pol II in magenta, and bacterial RNAP in brown. (B-D) Comparison of the active site (B), the bridge helix (C) and the trigger loop (D) of RDR2 vs. those of yeast Pol II and bacterial RNAP. For each domain, corresponding amino acid regions are shown from the same models superimposed in (A), following the same color code. Invariant glycines in the bridge helix are highlighted in yellow, and two molecular hinges (HN and HC) in multi-subunit RNAPs are indicated in broken lines. Invariant histidines in the trigger loop of multi-subunit RNAPs and corresponding lysine resides in RDR2 are indicated.

In multi-subunit RNAPs, the bridge helix is a single α-helix within which two invariant glycines serve as molecular hinges (BH-H_N_ and BH-H_C_), providing conformational flexibility during translocation of the RNA and DNA chains (36-38). In RDR2, two α-helices connected by a short loop (aa 880-884) comprise the corresponding structure (Fig. 2C). A glycine (G886) conserved in nearly all cellular RDRs examined (Fig. S10) is located at the junction between one of these helices and the intervening loop. The same structural arrangement is observed for the bridge helix of Neurospora QDE-1ΔN (26). In Arabidopsis RDR6, a paralog of RDR2, mutation G921D, which changes the conserved glycine corresponding to RDR2 G886 to aspartic acid (G921D), results in a null mutant phenotype (39), providing genetic evidence for the importance of this glycine.

Beyond the bridge helix (aa 885-896) in the C-terminal direction is an α-helix (aa 906-922) followed by a loop (aa 923-939) that likely constitutes the trigger loop. In multi-subunit RNAPs, the trigger loop is a flexible module that helps position substrate NTPs in the catalytic center during each nucleotide addition cycle (40). The putative RDR2 trigger loop is shorter than that of multi-subunit RNAPs and our structure appears to resemble the “closed” conformation observed for a substrate-bound multi-subunit RNAP elongation complex (Fig. 2D) (41, 42). The amino acid composition of the bridge helix and trigger loops of RDR2 or QDE-1 differ from those of multi-subunit RNAPs (Fig. 2D, Fig. S11). In multi-subunit RNAPs, an invariant histidine residue (His-β’1242 for *T. thermophilus* RNAP and His-1085 for yeast Rpb1) is critical for interaction with the β-phosphate of incoming NTPs. A lysine is present at the corresponding position in RDR2 (Lys-923), Neurospora QDE-1 (Lys-1119) and the crAss-like phage RNA polymerase, phi14:2 (Lys-1615) (Drobysheva et al., 2021), suggesting that Lys-923 of RDR2 may be functionally equivalent to His-1085 of Pol II.

### The N-terminal RRM domain has RNA binding activity

RDR2 amino acids 7-95 are predicted to form a β-α-β-β-α-β type RRM (RNA-recognition motif) domain based on Genome3D analysis (43). Alignment to other RRM domain-containing proteins reveals similarities to the weak consensus sequences deduced for the so-called ribonucleoprotein (RNP) motifs, RNP1 (aa 54-61) and RNP2 (aa12-17) (Fig. 3A) (44). Consistent with these analyses, the structural model for the RDR2 N-terminal domain fits a β-α-β-β-α-β RRM fold, with the RNP1 and RNP2 motifs residing in the β3 and β1 elements but with the β4 element not apparent (Fig. 3C) (see Methods).

**Figure 3.**
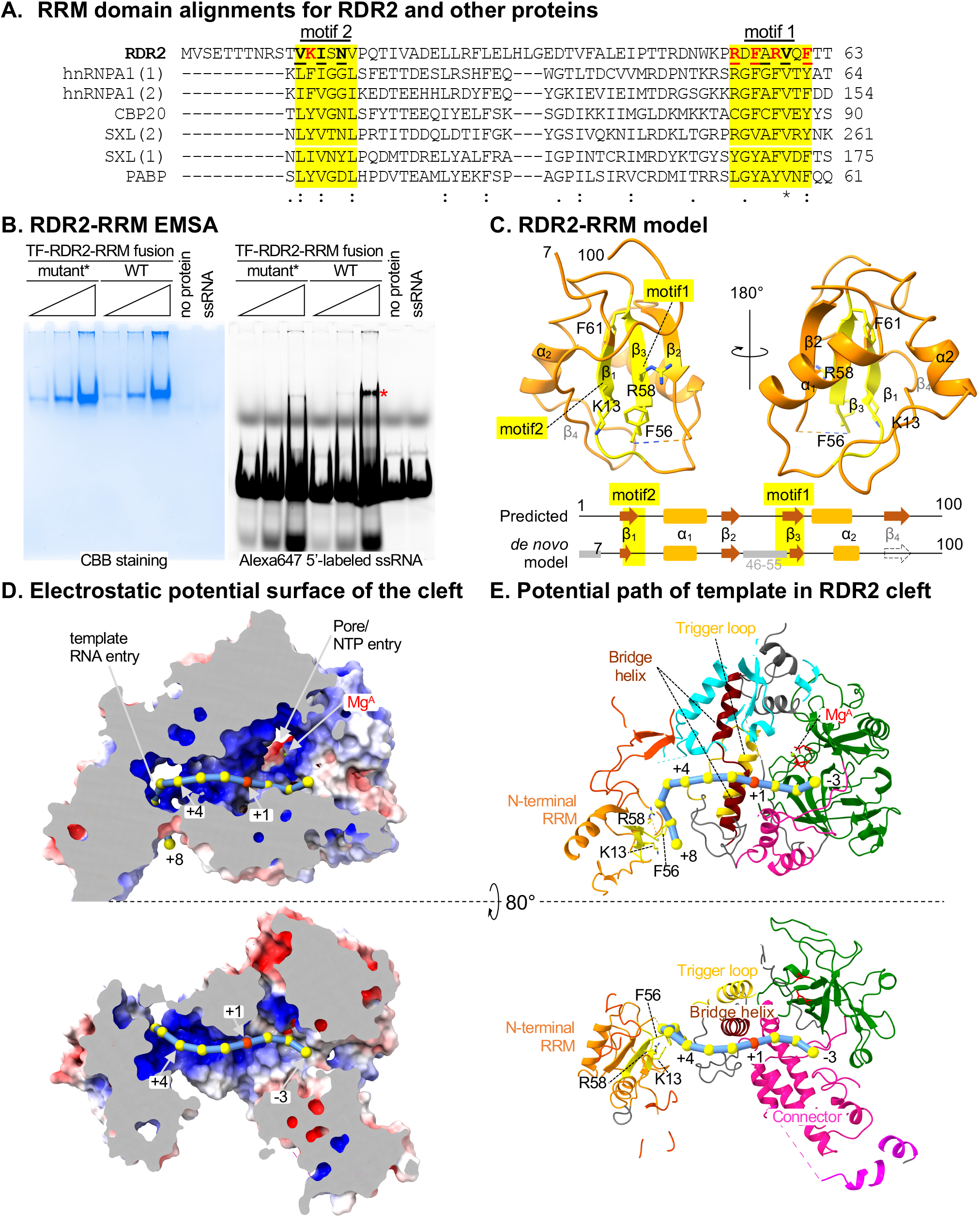
An N-terminal RNA recognition motif (RRM) domain influences the predicted path of template RNA. (A) RRM domain alignments for RDR2 (aa1-63), human proteins: heterogeneous nuclear ribonucleoprotein A1 (hnRNPA), nuclear cap-binding protein subunit 2 (CBP) and polyA-binding protein 1 (PABP) and *Drosophila melanogaster* Sex lethal (SXL). Ribonucleoprotein 1 and 2 motifs (RNP1 and RNP2), having consensus sequences [RK]-G-[FY]-[GA]-[FY]-[ILV]-X-[FY] and [ILV]-[FY]-[ILV]-X-N-L, respectively, are shaded yellow. Basic or aromatic residues potentially interacting with RNA are shown in red. (B) Electrophoretic mobility shift assay of a recombinant Trigger Factor-RDR2(aa1-100) fusion protein binding to a 37 nt RNA fluorescently labeled with Alexa 647. After acquiring the fluorescence image (right), the gel was fixed and stained with Coomassie Brilliant Blue, CBB (left). Wild-type (WT) and mutant forms of RDR2 were tested, with five amino acids of motifs 1 and 2 (shown in red in panel A) substituted by Ala in the mutant. An asterisk indicates the RNA-protein complex. (C) Structure of RDR2(aa 1-100), with RRM domain motifs 1 and 2 colored yellow. The diagram at the bottom compares a computationally predicted secondary structure for the region to our structural model. (D) Electrostatic potential surface of RDR2 calculated using Adaptive Poisson-Boltzmann Solver (APBS) in Pymol (69, 70) with negative, neutral, and positive charges shown in red, white, and blue, respectively. RNA is modeled as beads on a string with the red bead indicating the position where complementary strand synthesis would begin (+1). Mg++ and likely NTP entry pore positions are indicated. (E) Ribbon diagram views of the surface models shown in (D). Domains are colored as in Figure 1D. Additional views are shown in Figure S12.

To test whether RDR2’s N-terminal domain (aa 1-100, hereinafter referred to as RDR2-RRM) displays RNA-binding activity, we expressed in *E. coli* RDR2-RRM fused, in-frame, to Trigger-Factor (TF). A fusion with a mutant form of RDR2-RRM^mut^, having five aromatic or positively charged amino acids of the RNP1 or RNP2 motifs changed to alanine, was also generated. Electrophoretic mobility shift assays (EMSA) demonstrated that TF-RDR2-RRM exhibits ssRNA binding that is substantially impaired in TF-RDR2-RRM^mut^, consistent with RNA binding activity (Fig. 3B).

### Prediction of the RNA template path

A structure for RDR2 engaged with RNA has not yet been achieved, but structural and functional data suggest a potential path for the template RNA within RDR2. Examination of the electrostatic surface potential shows a positively charged surface starting with the RDR2-RRM and continuing along the cleft and into the active site (Fig. 3D). If an RNA template is modeled along the positively charged surface, with the transcription initiation position at the Metal A site defined as nucleotide position +1, the RNA path is relatively straight from +1 to +5, but then changes trajectory to interact with the binding surface of the RRM that helps form the putative RNA entry channel (Fig. 3D, 3E). In the opposite direction, the opening between the two DPBB domains and the connector domain is the presumed dsRNA exit channel (see “back” view, Fig. S12).

### RDR2 engages single-stranded RNAs longer than 7nt and initiates internal to their 3’ ends

The distance between the RNA-binding surface of RDR2-RRM and the catalytic site is 41 Å, which correspond to ∼7 nt of single-stranded RNA, assuming a nucleotide spacing of ∼6 Å. Our model thus predicts that an RNA of ∼7 nt or longer would be needed to serve as a template for RDR2 (Fig. 3E). To test this prediction, transcription reactions were conducted using single-stranded RNAs that varied in length from 5 nt to 15 nt. No ^32^P-labeled RNA transcripts were detected using the 5 nt template (Fig. 4A, lane 3) and only a low level of transcription occurred using the 7 nt template, but robust transcription occurred using templates of 9 nt or longer (Fig. 4A, lanes 4-8).

**Figure 4.**
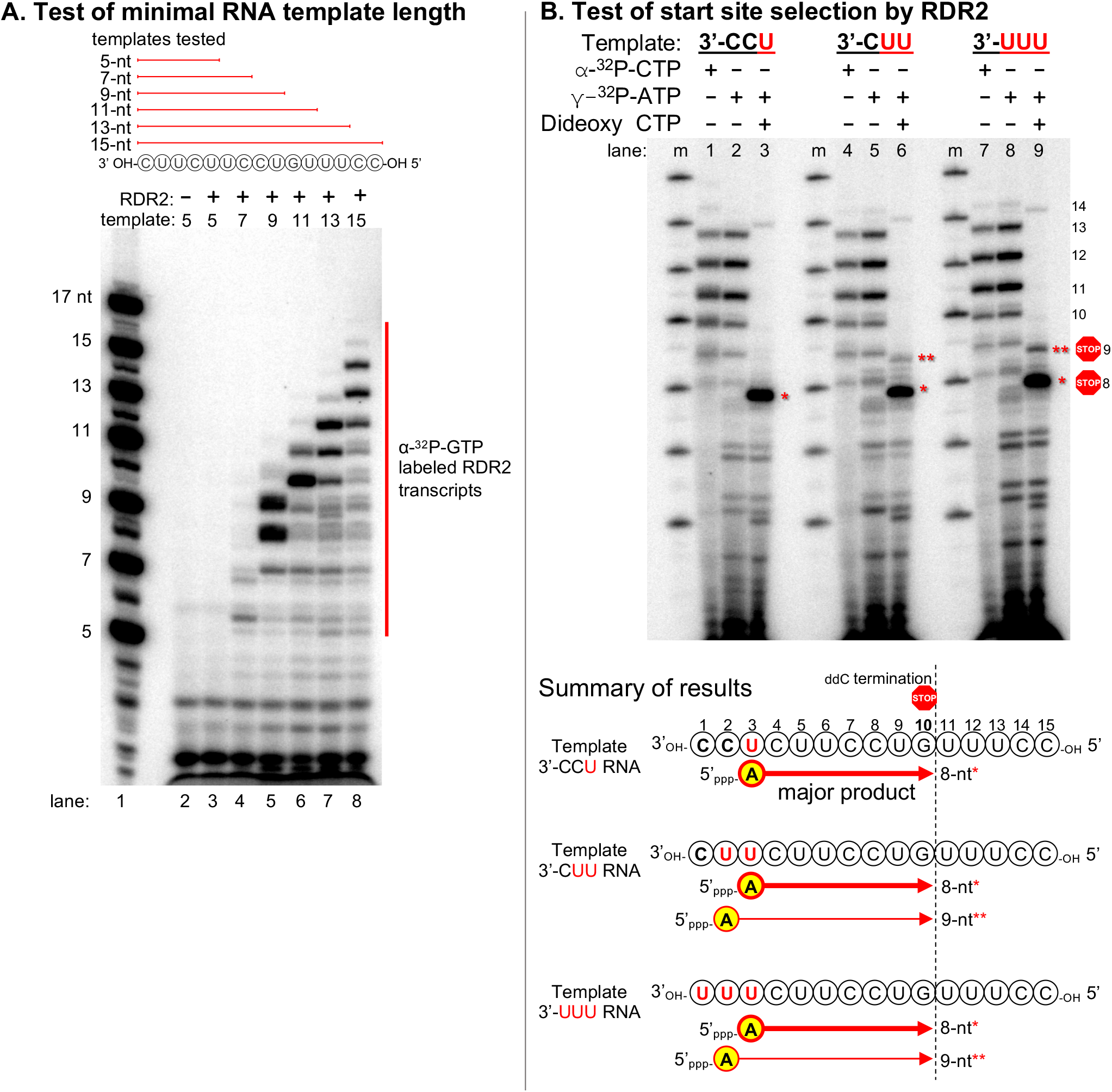
RDR2 initiates internal to its RNA templates. (A) Test of minimal RNA template length required for RDR2 transcription. Single-stranded RNAs of 5, 7, 9, 11, 13 or 15 nt were tested, with transcripts labeled by incorporation of α-^32^P-GTP. In lane 1, RNAs 5’ end-labeled using γ-^32^P-ATP serve as size markers. (B) Test of start site selection by RDR2. Using 15-nt ssRNA templates that have 3’-CCU, 3’-CUU and 3’-UUU at their 3’ ends, RDR2 transcription was carried out in the presence of α-^32^P-CTP, γ-^32^P-ATP or γ-32P-ATP plus 2’,3’-dideoxy CTP (ddC).

The most abundant transcripts in Fig, 4A were consistently 1-2 nt shorter than the templates (Fig. S13). To test the hypothesis that transcription initiates internal to the 3’ ends of template RNAs, we designed three 15 nt RNA templates that differ in the three nucleotides present at their 3’ ends; either 3’-CCU, 3’-CUU, or 3’-UUU (Fig. 4B). We then conducted RDR2 transcription reactions in which nascent transcripts were either body-labeled by incorporation of α-^32^P-CTP (Fig. 4B, lanes 1, 4, 7) or were 5’ end-labeled by γ-^32^P-ATP used as the initiating nucleotide (lanes 2,5,8). We also generated RNAs that initiated with γ-^32^P-ATP but were then terminated at a fixed position complementary to the guanosine at template position 10 upon incorporation of the chain terminator, 2’,3’-dideoxy CTP (ddC) (lanes 3,6,9).

Labeling with α-^32^P-CTP or γ-^32^P-ATP yielded similar transcription products for all three 15 nt templates, with transcripts of 8-13 nt being most abundant (compare lanes 1,2,4,5,7,8). Products of 14 nt were also detected, at much lower levels, but full-length 15 nt transcripts were undetectable or observed in only trace amounts. In the presence of ddC (lanes 3, 6, 9), a major labeled band of 8 nt was detected for all three templates, and a less abundant 9 nt band was observed for the 3’-CUU and 3’-UUU templates. Because only the 5’ terminal nucleotide retains the labeled gamma phosphate of γ-^32^P-ATP, the production of an 8 nt transcript that terminates at template position 10 indicates that transcription initiated with an adenosine complementary to the uridine at template position 3 (see summary diagrams of Fig. 4B). Likewise, 9 nt transcripts initiated at template position 2. Collectively, these results show that RDR2 initiates at positions complementary to the 2^nd^ or 3^rd^ nucleotides internal to the template.

### RDR2 can transcribe the RNA of an RNA-DNA hybrid if the 3’ end is accessible

How RDR2 engages Pol IV transcripts to generate dsRNAs is unknown. *In vitro*, Pol IV transcription of DNA templates in the absence of RDR2 yields persistent RNA-DNA hybrids (11). To determine if RDR2 can access the RNA strand of a RNA-DNA hybrid, we first tested a 37 nt RNA fully hybridized to a complementary 37 nt DNA, revealing that RDR2 fails to generate transcripts from this hybrid template (Fig. 5, lanes 2-3). However, if an unpaired, single-stranded region of 9 nt or longer is present at the 3’end of the RNA, and the rest is basepaired with DNA, RDR2 can transcribe the RNA (Fig. 5, lanes 4-8), apparently unwinding the downstream RNA-DNA hybrid region in the process. The unpaired portion of the DNA strand is dispensable for RDR2 engagement of the RNA 3’ end, as shown by the fact that it can be deleted without affecting RDR2 transcription of the remaining RNA-DNA hybrid (Fig. 5, lane 9; see bottom-most template diagram).

**Figure 5.**
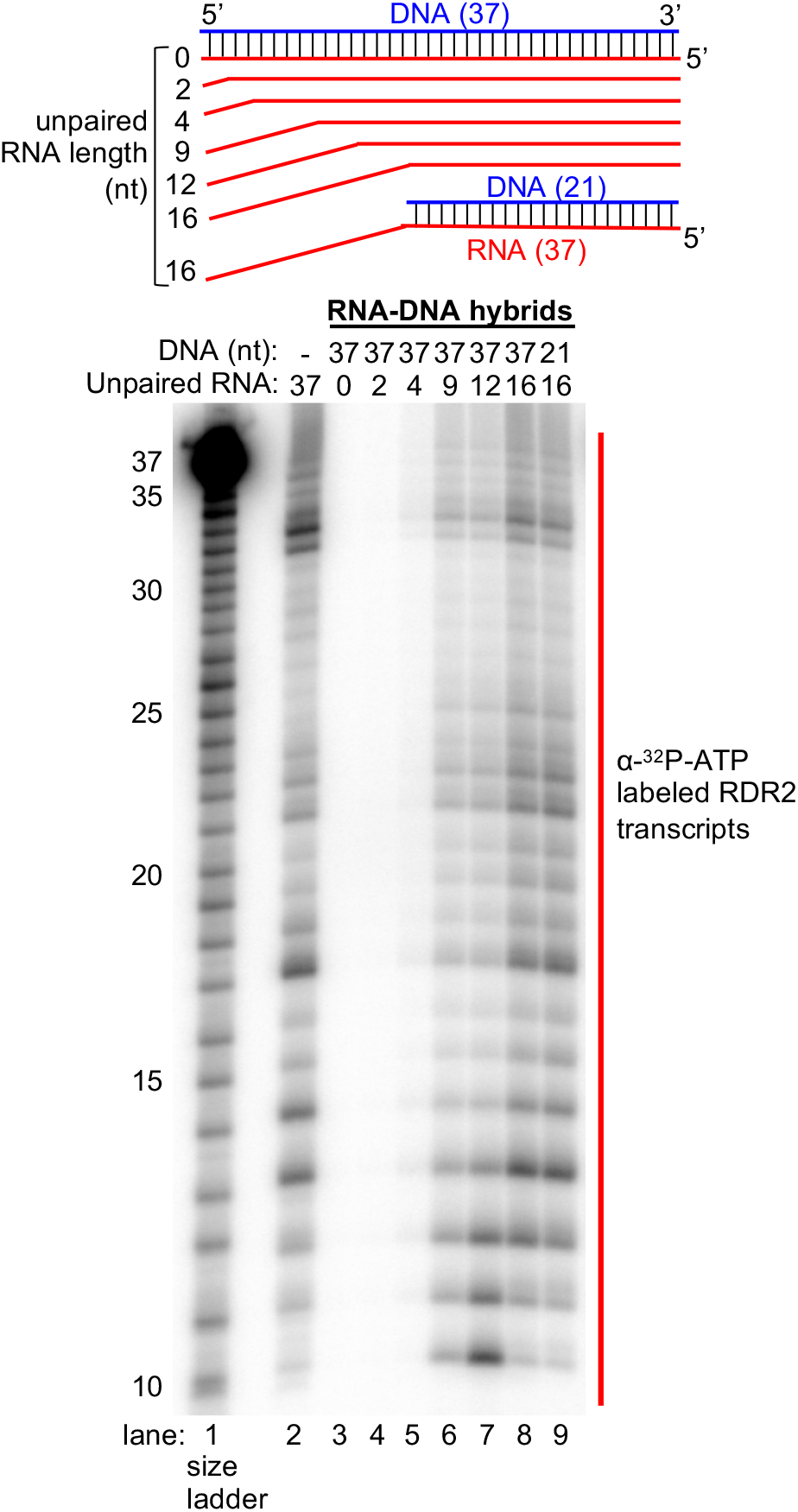
Test of RNA template engagement in the context of an RNA-DNA hybrid. As depicted in the diagram, 37-nt RNAs were hybridized to a 37-nt DNA strand to form hybrids with 0, 2, 4, 9, 12, or 16 nt of unpaired RNA at the 3’ end, then tested as templates for RDR2 transcription (lanes 3-8). A 21-nt DNA oligo was also used to generate a hybrid with 16 nt of unpaired RNA (lane 9). An RNA-only control (no DNA) was tested in lane 2 (same RNA as in lane 3). RNA transcripts were labeled by incorporation of α-^32^P-ATP. Partially hydrolyzed 5’ end-labeled 37-nt RNA was used as a size ladder in lane 1.

It is noteworthy that the need for 9 or more unpaired nucleotides at the 3’ end of a RNA-DNA hybrid template is consistent with the minimal length of single-stranded RNA needed for transcription (see Fig. 4A) and is equivalent to the distance from the RRM domain to the active site (∼7 nt) plus the two extra nucleotides needed for initiation to begin at the 3^rd^ position internal to the template (see Fig. 4B).

### Non-template-template DNA strand re-annealing displaces the RNA strand of RNA-DNA hybrids to enable RDR2 transcription

Coupling of Pol IV transcription and RDR2 transcription requires Pol IV transcriptional termination or arrest within a double-stranded DNA region (11). Previous assays of Singh et al. typically involved a 51 nt DNA oligonucleotide template annealed, at its 5’ end, to a 28 nt non-template DNA strand, forming a 27 bp double-stranded region with a 1 nt unpaired flap at the non-template strand’s 5’ end (see diagram of Fig. 6A). Pol IV transcription was then initiated using an RNA primer annealed at the template’s 3’ end, thus ensuring that resulting Pol IV transcripts have a defined 5’ end. Upon reaching the double-stranded portion of the template, Pol IV was shown to proceed only an additional 12-18 nt before arresting and/or terminating, coincident with RDR2 engagement of the Pol IV transcript and synthesis of the complementary strand (11). Using the same 51 nt DNA template and 28 nt non-template DNA strands used by Singh et al., and a 39 nt RNA corresponding to a Pol IV nascent transcript that extended 15 nt into the double-stranded DNA region, we examined the ability of RDR2 to access the RNA and convert it into dsRNA (Fig. 6A). When provided with the RNA alone, RDR2 generates abundant transcripts, forming a ladder of bands (Fig. 6A, lane 1). Some of these bands are longer than full-length due to folding of the RNA into an asymmetric stem-loop and further elongation of the 3’ end using the stem as a template. If the RNA is fully hybridized to the template DNA strand no RDR2 transcription occurs (Fig. 6A, lane 2). However, if the non-template DNA strand is included with the DNA template and RNA strands in the annealing/hybridizing reactions, RDR2 is able to access and transcribe the RNA strand (lane 3).

**Figure 6.**
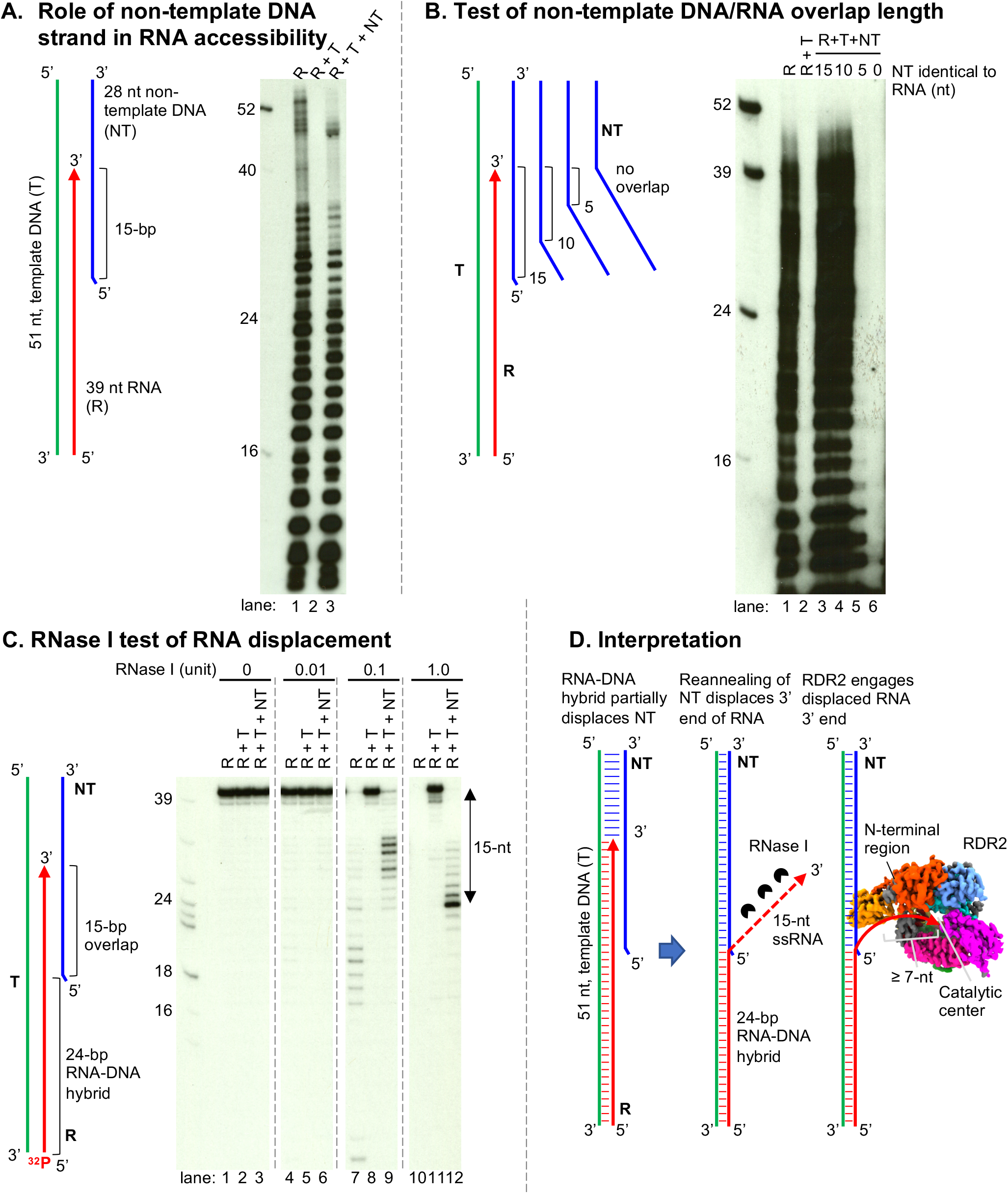
Role of non-template DNA in RDR2 engagement of RNAs in the context of RNA-DNA hybrids. A nucleic acid configuration that mimics the situation when Pol IV arrests 15-nt into a downstream dsDNA region was prepared by annealing 39 nt RNA (R, red), 51 nt template DNA (T, green) and 28 nt non-template DNA strands (NT, green). RDR2 transcription was tested using R only (lane 1), R+T (lane 2) and R+T+NT combinations (lane 3), with transcripts labeled by incorporation of α-^32^P-ATP. (B) Test of non-template DNA/RNA overlap length. Same assay as in (A), but varying the NT strand to have 10-, 5- or 0-nt of sequence identity (overlap) with the RNA strand. (C) RNase I test of RNA displacement. 5’ end-labeled R strands in R only, R+T or R+T+NT combinations were examined at four RNase I concentrations. (D) Graphical interpretation of the results in A-C.

We next tested the length of overlap between the non-template DNA strand and RNA strand needed for RDR2 to be able to transcribe the RNA (Fig. 6B). As in panel A, controls (lanes 1 and 2) show that RDR2 can transcribe free RNA but not RNA fully hybridized to template strand DNA. Inclusion of non-template DNA strands with 15 or 10 nt of identity to the 3’ end of the RNA resulted in RDR2 transcription of the RNA strand (Fig. 6B, lanes 3 and 4), whereas non-template strands with only 5 or 0 nt of overlap did not enable RDR2 transcription (lanes 5 and 6).

The results of Figures 6A and 6B suggest that the non-template DNA strand competes with the RNA strand for base pairing with the template DNA strand, resulting in displacement of the RNA’s 3’ end, thereby enabling RDR2 to engage and transcribe the RNA. To test this hypothesis, we examined the RNase I sensitivity of the RNA strand alone or in the presence of the template and/or non-template DNA strands (Fig. 6C). For this experiment, the 39-nt RNA was labeled with ^32^P at its 5’end. In the absence of complementary DNA, the RNA strand was readily digested by RNase I (lanes 7 and 10) but when hybridized to the DNA template strand, the RNA was RNAse-resistant (Fig. 6C, lanes 5,8,11). Inclusion of the non-template strand with the RNA and DNA template strand resulted in RNase I trimming of the RNA, causing the accumulation of a prominent 24 nt product (lane 9,12). This trimming of 15 nt corresponds to the 15-nt of overlap between the RNA and non-template DNA strands, consistent with the competitive base-pairing hypothesis (see Fig. 6D).

## Discussion

Cellular RDRs are widely conserved in eukaryotes and play essential roles in RNA silencing pathways. They have been classified into three major lineages, RDRα, RDRβ and RDRγ, with RDRα enzymes possessing the largest number of conserved motifs (2). Arabidopsis RDR2 belongs to the RDRα clade, thus we expect its structure to be informative for studies of other alpha subfamily members involved in gene silencing, including the Arabidopsis enzymes RDR1 and RDR6, *Neurospora crassa* SAD-1, *Schizzosaccharomyces pombe* Rdp1 and *C*.*elegans* EGO-1/RRF1/RRF3. Fungal QDE-1 proteins represent the RDRγ clade. The catalytic cores of RDR2 and QDE-1ΔN are similar. However, an intriguing difference between the enzymes is that RDR2 behaves as a monomer during purification (8) and upon imaging by cryoEM whereas QDE-1ΔN proteins were crystalized as homodimers. For QDE-1 enzymes, a “two-stroke motor” mechanism has been proposed, in which one dimer subunit becomes active (“closed”) upon RNA binding while the other subunit becomes inactive (“open”) (25, 26).

### Genetic results relevant to the RDR2 structure

There is a considerable body of genetic evidence relevant to the RDR2 structure. The Asp-triad loop, aa 830-DLDGD-834, is known to be critical for catalytic activity, as shown by the loss of activity upon substituting three alanines for DGD at positions 832-834 or upon substituting glycine for aspartate at position 834 (8, 45). Moreover, a genetic screen recovered G833E as an RDR2 loss-of-function mutant (46).

In RDR6, a paralog of RDR2 within the RDRα clade (Fig. S9A), mutations at numerous amino acid positions that are also conserved in RDR2 disrupt post-transcriptional gene silencing, including: P611L within the DPBB1 domain; G825E, D826N and S860F within the DPBB2 domain; G866E within the ASP-triad loop; G921D within the bridge helix, and E429K and E453K within the slab domain (47-49) (39) (Fig. S14A).

The RDR2 head domain is composed of five α helices. A nonsense mutation, W1083* that truncates the last 51 residues of the C-terminal head domain was recovered as a silencing defective mutant (50). In *Zea mays*, a Mu transposon insertion at amino acid position 962, within the neck3 helix leading to the head domain of the RDR2 ortholog, MOP1 resulted in the loss of MOP1 function (51). In RDR6, multiple nonsense mutations within the head domain disrupt silencing, including Q1145*, W1160*, Q1055* (48, 52) and W1039* (53) (Figure S14B). Collectively, these mutations provide genetic evidence for the importance of the head domain, yet the function of the domain remains unclear. In the RDR2 structure, the tip of the head domain faces toward the cleft (Fig. 1D). Importantly, our 3D reconstruction lacked density corresponding to the tip of the head (aa 1029-1045) but 3D variable analysis by cryoSPARC (54) captured the extension of the tip of the head pushing toward the cleft (see Movie S1). We speculate that the tip of the head may interact directly with incoming template RNA to help guide the template to the active site.

Functions for the beta-barrel-like, 4-helical bundle, slab, and neck domains remain unclear. We note that the slab domain of RDR2 contains structural features that resemble the fork loops and link domain helix in yeast Pol II. Specifically, when superimposed, the RDR2 slab (aa 470-488) corresponds well with fork loop 3 of yeast Rpb2 (aa 521-541), whereas RDR2 amino acids 511-521 form an α-helix which may correspond to the Rpb2 link domain helix (aa 757-776) (Fig. S15). The fork domain composed of loop 1-3 (aa 466 –546) of Rpb2 contacts the RNA-DNA hybrid, the template strand and downstream DNA (55-57) whereas amino acids of the link domain interact with NTPs (38). Thus, it is possible that the slab domain of RDR2 may interact with template RNA and NTPs.

### A potential structural basis for RDR2 initiation internal to template 3’ ends

For multi-subunit RNAPs, the initiating NTP (iNTP) is stabilized by interactions with conserved positively charged or polar amino acids (e.g. *T. thermophilus* β subunit amino acids Gln-567, Lys-838, Lys-846 and His-999; or corresponding *S. cerevisiae* Pol II Rpb2 subunits Gln-776, Lys-979, Lys-987 and His-1097), by base-stacking interactions with the template base at position -1 (1 nt upstream of the initiation site, +1) and by water-mediated interactions between a phosphate group of the iNTP and template bases at the -1 and -2 positions (Fig. S16) (58). We questioned whether analogous interactions might explain RDR2’s initiation 2-3 nt internal to its templates. In the RDR2 structure, positively charged or polar residues Lys-589 (equivalent to *T. thermophilus* Lys-846) and Gln-582 (equivalent to *T. thermophilus* Lys-838) are present in the equivalent iNTP α-phosphate interacting positions (Fig. S16) and are invariably conserved among the RDRs examined. However, potential iNTP γ-phosphate interacting amino acids are less conserved in RDR2 and other RDRs (Fig. S10). In the position equivalent to *T. thermophilus* His-999, RDR2 has Arg-622 whereas other RDRs have Lys, Asn or other amino acids (Fig. S16 and S10). Polar amino acids are conserved at RDR positions corresponding to RDR2 Ser-525 and may correspond to *T. thermophilus* Gln-567. These structural comparisons suggest that initiation site selection by RDR2 may have some mechanistic similarities to initiation site selection in multisubunit RNAPs, a possibility worthy of further structural investigation.

Importantly, the fact that RDR2 initiates internal to the 3’ ends of its RNA templates has biological significance, because it causes the resulting dsRNA to have a 1 or 2 nt 3’ overhang. In other studies, we have found that dsRNAs with 1 or 2 nt 3’ overhangs are the preferred substrates for DICER-LIKE 3 (DCL3). Thus, RDR2’s internal initiation is important for siRNA biogenesis.

### A model for RDR2 engagement of backtracked Pol IV transcripts

The transcription reactions of Pol IV and RDR2 are tightly coupled such that Pol IV transcripts are channeled to RDR2 rather than being released. It is unclear how channeling occurs, but an important clue has been that basepairing between the DNA template and nontemplate strands is required, inducing both Pol IV arrest and RDR2 engagement of the Pol IV transcripts for second-strand synthesis (11). Our current study provides additional insights, showing that RDR2 can engage the RNA of an RNA-DNA hybrid if ∼9 nt (or more) of the RNA’s 3’ end is unpaired from the DNA. Moreover, we’ve shown that template-nontemplate strand reannealing can displace the 3’ end of the RNA, thereby enabling RDR2 to engage and transcribe it into dsRNA (Fig. 5 and 6A-D). RNA 3’ end displacement that is coincident with DNA template-nontemplate strand reannealing is known to occur during multisubunit RNA polymerase backtracking, suggesting the model shown in Figure 7. In this model, Pol IV transcription initiates within an open DNA bubble but can elongate only 12∼18 nt into the adjacent dsDNA region before stalling, possibly as a result of amino acid substitutions and deletions in the “rudder” and “zipper” loops thought to be needed for transcription bubble perpetuation during elongation (18, 59, 60). Unable to go forward, we propose that Pol IV backtracking ensues, coinciding with the re-annealing of the nontemplate and the template DNA strands and the extrusion of the RNA’s 3’end (61). When the extruded 3’-end of the backtracked RNA becomes long enough, RDR2 engages the RNA and initiates second strand synthesis, extracting the RNA as Pol IV retreats. Upon completion of RDR2 transcription, RDR2 adds an untemplated nucleotide to the 3’ end of its transcripts (11) and the dsRNA is released from the Pol IV-RDR2 complex. This model is consistent with previous and current biochemical assays and accounts for the role of the nontemplate DNA strand and the direct transfer of RNA from Pol IV to RDR2.

**Figure 7.**
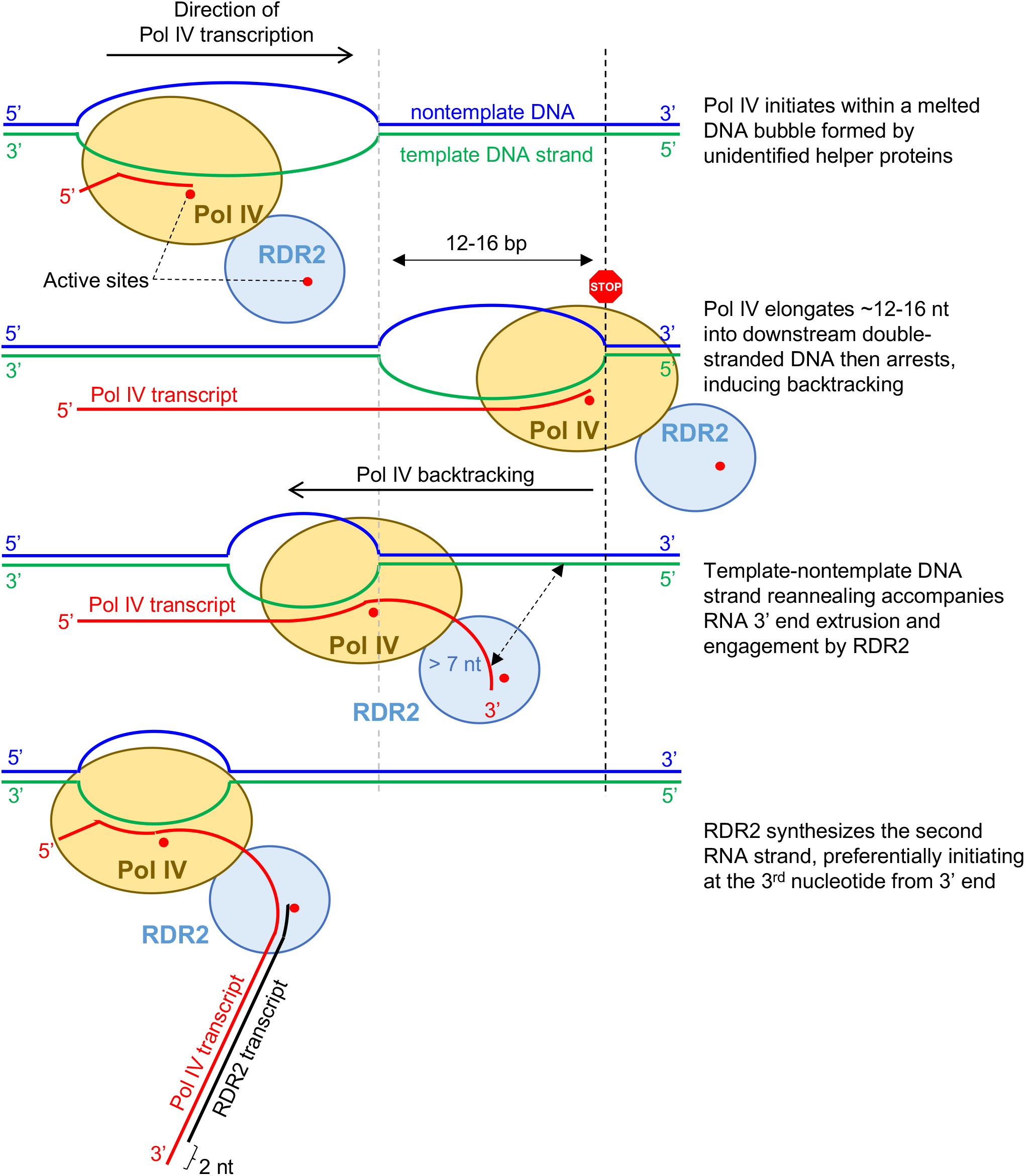
Backtracking model for Pol IV-RDR2 transcription coupling.

## Materials and Methods

### Construction of the baculovirus vector for expression of RDR2

A baculovirus vector termed pSEP10 that allows for the expression of large proteins in insect cells was engineered and used to produce RDR2. Briefly, a synthetic RDR2 open reading frame (ORF) was cloned into BamHI and Hind III sites of pSEP10, resulting in the SEP tag being fused in-frame to the RDR2 open reading frame and a Twin-Strep tag at the C-terminus. Details are provided in the Supplementary Information. Generation of baculovirus expressing RDR2 using the resulting pSEP10-RDR2 vector was performed as described previously (62)

### Expression, and purification of RDR2

Expression of recombinant RDR2 was optimized using the TEQC method (62) in which a 200 ml culture of Hi5 cells (Expression Systems, Inc) was infected at an estimated multiplicity of infection (eMOI) of 4. After a 96-hour incubation at 27°C, cells were harvested, frozen in liquid nitrogen and kept at -80°C until use. Purification of recombinant RDR2 from lysed cell pellets was carried out by Strep-Tactin affinity purification followed by Hitrap Q column chromatography. Details are provided in the Supplementary Information. Hitrap Q fractions containing the recombinant RDR2 were concentrated using a spin column (100 kD cut-off) to a final concentration of 4.8 mg/ml, as measured using a Bradford assay.

### Cryo-EM sample preparation and data collection

Grid preparation, grid screening and data collection were performed in the CryoEM facility at the HHMI Janelia Research Campus. Datasets were collected with no stage tilt or with 30° of stage tilt using a Titan Krios microscope (Thermo Fisher) operated at 300 kV and equipped with a K3 camera (Gatan). Detailed methods for grid preparation and data collection are described in the Supplementary Information section.

### Image Processing

The map used to build the *de novo* atomic model was obtained by image processing using Relion 3.1 (28, 29, 63). The workflow and detailed procedures for each step are described in the Supplementary Information methods section and Figures S1 and S2. Briefly, after beam-induced motion correction and contrast transfer function (CTF) estimation, an initial 3D model at ∼4.3Å resolution was generated from 2D-reference-picked particles, which was then used as a 3D-reference to select a sub-set of particles. 2D classifications, 3D classifications, iterative CTF refinement, Bayesian polishing and metadata filtering of these selected particles led to the final map, “Map1” of ∼3.10 Å. Independent processing of the same dataset by cryoSPARC v3.2.0, which resulted in a nearly identical 3D reconstruction at ∼3.57 Å, “Map2”, is described in the supplemental methods and Fig. S3. All figures for cryo-EM density maps, models and model surfaces were generated using Chimera v1.15 (64) or Chimera X software (65).

### Model building

Map1 was sharpened using the DeepEMhancer tool (66) of the COSMIC^2^ science gateway (67) (Map1) then used for model building. Model building for amino acid residues 61-1121 of Map1 was carried out *de novo* using the automated program Buccaneer (31) and Emap2sec (68) was used to assess secondary structure propensity in the EM map. Model building for residues 1-60 was aided by a computational model generated by I-TASSER (Yang et al., 2015). Detailed methods for the model building and refinement process are described in the supplemental methods.

### RDR2 Transcription Assays

RDR2 transcription assays were conducted by mixing templates (RNA or RNA/DNA hybrids), nucleotide triphosphates (NTPs) supplemented with [^32^P]-labeled ATP, GTP or CTP, and RDR2. After incubating at room temperature to 27 °C for 1-2 hours, the reactions were either treated by proteinase K or passed through PERFORMA spin columns (Edge Bio), precipitated, and resolved by 15% or 17% denaturing PAGE. The gels were then subjected to either phosphorimaging or autoradiography using X-ray films. In Fig. 1B, the reaction was analyzed by 15% native PAGE instead to detect double-stranded RNA. Details are provided in the Supplementary Information methods section.

### Expression and purification of recombinant TF-RDR2-RRM fusion protein

To produce recombinant the TF-RDR2-RRM fusion protein, the DNA sequence encoding RDR amino acids 1-100 (or a mutant version containing five alanine substitutions as described in Fig. 3), was codon-optimized for *E. coli*, synthesized by GenScript and cloned into the pCold-TF vector (Takara) BamHI and HindIII sites. The resulting construct was transformed into ArcticExpress competent cells (Agilent). Protein expression was induced by cold-shock. The fusion protein was affinity purified using Ni-TNA agarose (Qiagen). Details are provided in the Supplementary Information methods section.

### Electrophoretic mobility shift assay

Recombinant TF-RDR2-RRM proteins (at 9, 3 or 1 μM) were incubated with a 37 nt single stranded RNA (final 2 μM), 5’ end-labeled with Alexa647, in 25 mM HEPES-KOH pH7.6, 50 mM NaCl, 2 mM MgCl_2_ and 4 units of RNase Inhibitor Murine (NEB, M0314). After 30 min at room temperature, reactions were subjected to 6% native PAGE and fluorescence imaging followed by staining with Coomassie Brilliant Blue.

## Data Availability

Structural data have been deposited in PDB (PDB ID: 7ROZ, 7RQS) and the Electron Microscopy Data Bank (EMDB ID: EMD-24610, EMD-24635).

## Acknowledgments

This research was supported by NIH grant GM077590 to CSP, NIH grant GM111695 to YT and funds to CSP as an Investigator of the Howard Hughes Medical Institute. JS was supported, in part, by a Carlos O. Miller graduate fellowship at Indiana University. We thank the Drosophila Genome Resource Center at Indiana University, Bloomington for insect cell culture facilities used for RDR2 overexpression. We thank Drs. Kihara, Terashi, and SR Maddhuri Venkata Subramaniya at Purdue University for help assessing secondary structures of our EM map using the program Emap2sec, and Dr. Georgiadis for advice on model building. We acknowledge the Research Technologies division of University Information Technology Services in Indiana University for providing supercomputing and storage resources, supported, in part, by Lilly Endowment, Inc. funding to the Indiana University Pervasive Technology Institute. Molecular graphics and analyses were performed with UCSF ChimeraX, developed by the Resource for Biocomputing, Visualization, and Informatics at the University of California, San Francisco, with support from National Institutes of Health R01-GM129325 and the Office of Cyber Infrastructure and Computational Biology, National Institute of Allergy and Infectious Diseases.

## Author Contributions

CSP and YT conceived the project. AF and YT determined maps for RDR2. YT built the models. AF conducted experiments of Fig. 1A, 1B, 3B, 4A, 4B, and 5. JS conducted the experiments of Fig. 6A, 6B and 6C using RDR2 produced by VM. ER, SW, and FMM produced RDR2 for cryoEM and biochemical assays. RY, and MS performed cryoEM grid making, screening, and Titan Krios data collection in Janelia Farm cryoEM facility overseen by ZY. RY, MS, and ZY contributed to method writing for cryoEM data collections. JW performed initial RDR2 analyses by negative staining. ER, YT, AF, and JW conducted initial cryoEM analyses of RDR2 using a Talos Arctica instrument. AF, YT and CSP wrote the manuscript.

## References

1. M. Wassenegger, G. Krczal, Nomenclature and functions of RNA-directed RNA polymerases. Trends Plant Sci. 11, 142–151 (2006).

2. J. Zong, X. Yao, J. Y. Yin, D. B. Zhang, H. Ma, Evolution of the RNA-dependent RNA polymerase (RdRP) genes: Duplications and possible losses before and after the divergence of major eukaryotic groups. Gene 447, 29–39 (2009).

3. R. Martienssen, D. Moazed, RNAi and heterochromatin assembly. Cold Spring Harb. Perspect. Biol. 7, a019323 (2015).

4. A. Girard, G. J. Hannon, Conserved themes in small-RNA-mediated transposon control. Trends Cell Biol. 18, 136–148 (2008).

5. D. Holoch, D. Moazed, RNA-mediated epigenetic regulation of gene expression. Nat. Rev. Genet. 16, 71–84 (2015).

6. J. R. Haag et al., In Vitro Transcription Activities of Pol IV, Pol V, and RDR2 Reveal Coupling of Pol IV and RDR2 for dsRNA Synthesis in Plant RNA Silencing. Mol. Cell 48, 811–818 (2012).

7. J. A. Law, A. A. Vashisht, J. A. Wohlschlegel, S. E. Jacobsen, SHH1, a homeodomain protein required for DNA methylation, as well as RDR2, RDM4, and chromatin remodeling factors, associate with RNA polymerase IV. PLoS Genet. 7, e1002195 (2011).

8. V. Mishra et al., Assembly of a dsRNA synthesizing complex: RNA-DEPENDENT RNA POLYMERASE 2 contacts the largest subunit of NUCLEAR RNA POLYMERASE IV. Proc. Natl. Acad. Sci. U. S. A. 118, e2019276118 (2021).

9. T. Blevins et al., Identification of Pol IV and RDR2-dependent precursors of 24 nt siRNAs guiding de novo DNA methylation in Arabidopsis. eLife 4, e09591 (2015).

10. J. X. Zhai et al., A One Precursor One siRNA Model for Pol IV-Dependent siRNA Biogenesis. Cell 163, 445–455 (2015).

11. J. Singh, V. Mishra, F. Wang, H. Y. Huang, C. S. Pikaard, Reaction Mechanisms of Pol IV, RDR2, and DCL3 Drive RNA Channeling in the siRNA-Directed DNA Methylation Pathway. Mol. Cell 75, 576–589.e575 (2019).

12. Z. X. Xie et al., Genetic and functional diversification of small RNA pathways in plants. PLoS Biol. 2, 642–652 (2004).

13. I. R. Henderson et al., Dissecting Arabidopsis thaliana DICER function in small RNA processing, gene silencing and DNA methylation patterning. Nature Genet. 38, 721–725 (2006).

14. A. T. Wierzbicki, J. R. Haag, C. S. Pikaard, Noncoding Transcription by RNA Polymerase Pol IVb/Pol V Mediates Transcriptional Silencing of Overlapping and Adjacent Genes. Cell 135, 635–648 (2008).

15. A. T. Wierzbicki, T. S. Ream, J. R. Haag, C. S. Pikaard, RNA polymerase V transcription guides ARGONAUTE4 to chromatin. Nature Genet. 41, 630–634 (2009).

16. X. F. Cao, S. E. Jacobsen, Role of the Arabidopsis DRM methyltransferases in de novo DNA methylation and gene silencing. Curr. Biol. 12, 1138–1144 (2002).

17. J. A. Law, S. E. Jacobsen, Establishing, maintaining and modifying DNA methylation patterns in plants and animals. Nat. Rev. Genet. 11, 204–220 (2010).

18. J. Singh, C. S. Pikaard, Reconstitution of siRNA Biogenesis In Vitro: Novel Reaction Mechanisms and RNA Channeling in the RNA-Directed DNA Methylation Pathway. Cold Spring Harb. Symp. Quant. Biol. 84, 195–201 (2019).

19. R. M. Erdmann, C. L. Picard, RNA-directed DNA Methylation. PLoS Genet. 16, e1009034 (2020).

20. M. R. Motamedi et al., Two RNAi complexes, RITS and RDRC, physically interact and localize to noncoding centromeric RNAs. Cell 119, 789–802 (2004).

21. S. Bhattacharjee, B. Roche, R. A. Martienssen, RNA-induced initiation of transcriptional silencing (RITS) complex structure and function. RNA Biol. 16, 1133–1146 (2019).

22. E. M. Maine et al., EGO-1, a putative RNA-dependent RNA polymerase, is required for heterochromatin assembly on unpaired DNA during C-elegans meiosis. Curr. Biol. 15, 1972–1978 (2005).

23. P. K. T. Shiu, N. B. Raju, D. Zickler, R. L. Metzenberg, Meiotic silencing by unpaired DNA. Cell 107, 905–916 (2001).

24. M. Freitag et al., DNA methylation is independent of RNA interference in neurospora. Science 304, 1939–1939 (2004).

25. X. L. Qian et al., Functional Evolution in Orthologous Cell-encoded RNA-dependent RNA Polymerases. J. Biol. Chem. 291, 9295–9309 (2016).

26. P. S. Salgado et al., The structure of an RNAi polymerase links RNA silencing and transcription. PLoS Biol. 4, 2274–2281 (2006).

27. S. H. W. Scheres, RELION: Implementation of a Bayesian approach to cryo-EM structure determination. J. Struct. Biol. 180, 519–530 (2012).

28. J. Zivanov, T. Nakane, S. H. W. Scheres, Estimation of high-order aberrations and anisotropic magnification from cryo-EM data sets in RELION-3.1. IUCrJ 7, 253–267 (2020).

29. K. Ramlaul, C. M. Palmer, T. Nakane, C. H. S. Aylett, Mitigating local over-fitting during single particle reconstruction with SIDESPLITTER. J. Struct. Biol. 211, 107545 (2020).

30. A. Punjani, J. L. Rubinstein, D. J. Fleet, M. A. Brubaker, cryoSPARC: algorithms for rapid unsupervised cryo-EM structure determination. Nat. Methods 14, 290–296 (2017).

31. K. Cowtan, The Buccaneer software for automated model building. 1. Tracing protein chains. Acta Crystallogr. D Biol. Crystallogr. 62, 1002–1011 (2006).

32. J. Y. Yang et al., The I-TASSER Suite: protein structure and function prediction. Nat. Methods 12, 7–8 (2015).

33. P. Cramer, D. A. Bushnell, R. D. Kornberg, Structural basis of transcription: RNA polymerase II at 2.8 angstrom resolution. Science 292, 1863–1876 (2001).

34. L. M. Iyer, E. V. Koonin, L. Aravind, Evolutionary connection between the catalytic subunits of DNA-dependent RNA polymerases and eukaryotic RNA-dependent RNA polymerases and the origin of RNA polymerases. BMC Struct. Biol. 3, 1 (2003).

35. A. V. Drobysheva et al., Structure and function of virion RNA polymerase of a crAss-like phage. Nature 589, 306–309 (2021).

36. F. Brueckner, P. Cramer, Structural basis of transcription inhibition by alpha-amanitin and implications for RNA polymerase II translocation. Nat. Struct. Mol. Biol. 15, 811–818 (2008).

37. R. O. Weinzierl, The Bridge Helix of RNA polymerase acts as a central nanomechanical switchboard for coordinating catalysis and substrate movement. Archaea 2011, 608385 (2011).

38. R. O. Weinzierl, The nucleotide addition cycle of RNA polymerase is controlled by two molecular hinges in the Bridge Helix domain. BMC Biol. 8, 134 (2010).

39. P. Lam et al., The exosome and trans-acting small interfering RNAs regulate cuticular wax biosynthesis during Arabidopsis inflorescence stem development. Plant Physiol. 167, 323–336 (2015).

40. D. G. Vassylyev et al., Crystal structure of a bacterial RNA polymerase holoenzyme at 2.6 angstrom resolution. Nature 417, 712–719 (2002).

41. D. G. Vassylyev et al., Structural basis for substrate loading in bacterial RNA polymerase. Nature 448, 163–168 (2007).

42. D. Wang, D. A. Bushnell, K. D. Westover, C. D. Kaplan, R. D. Kornberg, Structural basis of transcription: Role of the trigger loop in substrate specificity and catalysis. Cell 127, 941–954 (2006).

43. T. E. Lewis et al., Genome3D: exploiting structure to help users understand their sequences. Nucleic Acids Res. 43, D382–386 (2015).

44. C. Maris, C. Dominguez, F. H. T. Allain, The RNA recognition motif, a plastic RNA-binding platform to regulate post-transcriptional gene expression. FEBS J. 272, 2118–2131 (2005).

45. A. Devert et al., Primer-dependent and primer-independent initiation of double stranded RNA synthesis by purified Arabidopsis RNA-dependent RNA polymerases RDR2 and RDR6. PLoS One 10, e0120100 (2015).

46. P. Dunoyer, C. Himber, V. Ruiz-Ferrer, A. Alioua, O. Voinnet, Intra-and intercellular RNA interference in Arabidopsis thaliana requires components of the microRNA and heterochromatic silencing pathways. Nature Genet. 39, 848–856 (2007).

47. T. Elmayan et al., Arabidopsis mutants impaired in cosuppression. Plant Cell 10, 1747–1757 (1998).

48. P. Mourrain et al., Arabidopsis SGS2 and SGS3 genes are required for posttranscriptional gene silencing and natural virus resistance. Cell 101, 533–542 (2000).

49. X. Adenot et al., DRB4-dependent TAS3 trans-acting siRNAs control leaf morphology through AGO7. Curr. Biol. 16, 927–932 (2006).

50. L. M. Smith et al., An SNF2 protein associated with nuclear RNA silencing and the spread of a silencing signal between cells in Arabidopsis. Plant Cell 19, 1507–1521 (2007).

51. M. R. Woodhouse, M. Freeling, D. Lisch, Initiation, establishment, and maintenance of heritable MuDR transposon silencing in maize are mediated by distinct factors. PLoS Biol. 4, 1678–1688 (2006).

52. N. Jiang et al., Synergy between the anthocyanin and RDR6/SGS3/DCL4 siRNA pathways expose hidden features of Arabidopsis carbon metabolism. Nat. Commun. 11, 2456 (2020).

53. A. Peragine, M. Yoshikawa, G. Wu, H. L. Albrecht, R. S. Poethig, SGS3 and SGS2/SDE1/RDR6 are required for juvenile development and the production of trans-acting siRNAs in Arabidopsis. Genes Dev. 18, 2368–2379 (2004).

54. A. Punjani, D. J. Fleet, 3D variability analysis: Resolving continuous flexibility and discrete heterogeneity from single particle cryo-EM. J. Struct. Biol. 213, 107702 (2021).

55. M. L. Kireeva, C. Domecq, B. Coulombe, Z. F. Burton, M. Kashlev, Interaction of RNA Polymerase II Fork Loop 2 with Downstream Non-template DNA Regulates Transcription Elongation. J. Biol. Chem. 286, 30898–30910 (2011).

56. R. D. Kornberg, The molecular basis of eucaryotic transcription. Cell Death Differ. 14, 1989–1997 (2007).

57. L. Wang et al., Molecular basis for 5-carboxycytosine recognition by RNA polymerase II elongation complex. Nature 523, 621–625 (2015).

58. R. S. Basu et al., Structural Basis of Transcription Initiation by Bacterial RNA Polymerase Holoenzyme. J. Biol. Chem. 289, 24549–24559 (2014).

59. R. Landick, Functional Divergence in the Growing Family of RNA Polymerases. Structure 17, 323–325 (2009).

60. I. Toulokhonov, R. Landick, The role of the lid element in transcription by E-coli RNA polymerase. J. Mol. Biol. 361, 644–658 (2006).

61. A. C. M. Cheung, P. Cramer, Structural basis of RNA polymerase II backtracking, arrest and reactivation. Nature 471, 249–253 (2011).

62. T. Imasaki, S. Wenzel, K. Yamada, M. L. Bryant, Y. Takagi, Titer estimation for quality control (TEQC) method: A practical approach for optimal production of protein complexes using the baculovirus expression vector system. PLoS One 13, e0195356 (2018).

63. J. Zivanov et al., New tools for automated high-resolution cryo-EM structure determination in RELION-3. eLife 7, e42166 (2018).

64. E. Pettersen et al., UCSF Chimera--a visualization system for exploratory research and analysis. J. Comput. Chem. 25, 1605–1612 (2004).

65. E. F. Pettersen et al., UCSF ChimeraX: Structure visualization for researchers, educators, and developers. Protein Sci. 30, 70–82 (2021).

66. R. Sanchez-Garcia et al., DeepEMhancer: a deep learning solution for cryo-EM volume post-processing. Commun. Biol. 4, 874 (2021).

67. M. A. Cianfrocco, M. Wong, C. Youn, R. Wagner, A. E. Leschziner, COSMIC^2^: A Science Gateway for Cryo-Electron Microscopy Structure Determination. Practice & Experience in Advanced Research Computing Article 22, 1–5 (2017).

68. S. R. Maddhuri Venkata Subramaniya, G. Terashi, D. Kihara, Protein secondary structure detection in intermediate-resolution cryo-EM maps using deep learning. Nat. Methods 16, 911–917 (2019).

69. N. A. Baker, D. Sept, S. Joseph, M. J. Holst, J. A. McCammon, Electrostatics of nanosystems: Application to microtubules and the ribosome. Proc. Natl. Acad. Sci. U. S. A. 98, 10037–10041 (2001).

70. T. J. Dolinsky, J. E. Nielsen, J. A. McCammon, N. A. Baker, PDB2PQR: an automated pipeline for the setup of Poisson-Boltzmann electrostatics calculations. Nucleic Acids Research 32, W665–W667 (2004).

