## Supplemental information for "Structure and RNA template requirements of Arabidopsis RNA-DEPENDENT RNA POLYMERASE 2"

#### Detailed Methods

##### Construction of the baculovirus vector for expression of RDR2

A unique baculovirus vector termed pSEP10 that allows for the expression of large proteins in insect cells was used to produce RDR2 for structural studies. A DNA cassette encoding maltose-binding protein (MBP) fused to a synthetic protein termed "SED" and a human rhinovirus 3C protease (HRV 3C Protease) cleavage site (LEVLFQGP) was codon optimized for insect cell expression and synthesized by GenScript. This DNA cassette was inserted immediately 5' of the *Bam*HI site in the pFastBac vector (Invitrogen) using the SLIC method (1), yielding a construct termed pSEP6 vector (pYT1361). The MBP-SED fusion is termed the SEP tag. A DNA cassette encoding Tobacco Etch Virus nuclear-inclusion-a endopeptidase (TEV protease) site (ENLYFQG) and a twin Strep tag (SAWSHPQFEKGGGS-GGGSGGSAWSHP-QFEK) was also codon optimized for insect cell expression and synthesized by GenScript (Piscataway, NJ). This DNA cassette was inserted immediately 3' of the HindIII site of the pSEP6 vector by the SLIC method (1), yielding a construct termed pSEP10 vector (pYT1537). A synthetic RDR2 open reading frame, fused in-frame at the C-terminus to the Twin-Strep tag and codon-optimized for insect cell expression, was synthesized by GenScript and sub-cloned into the *Bam*HI and *Hind*III sites of the pSEP10 vector, yielding the construct pSEP10-RDR2 (MBP-SED-3C-RDR2-TEV-Twin Strep) (pYT1654). Generation of baculovirus expressing RDR2 using pSEP10-RDR2 vector expressing RDR2 in Sf9 cells was performed as described previously (2)

##### Expression, and purification of RDR2

Expression of recombinant RDR2 was optimized using the TEQC method (2) in which a 200 ml culture of Hi5 cells (Expression Systems, Inc) was infected at an estimated multiplicity of infection (eMOI) of 4. After a 96 hour incubation at 27°C, cells were harvested, frozen in liquid nitrogen and kept at -80°C until use. The cell pellet from the 200 ml culture were lysed in 50 ml of lysis buffer: 50 mM Hepes-KOH pH7.6, 400 mM potassium chloride, 10% glycerol, 0.01% NP-40, and 5 mM β-mercaptoethanol, and 0.5 ml of 100 x protease inhibitor mix (6 mM leupeptin, 0.2 mM pepstatin A, 20 mM benzamidine, and 10 mM PMSF) as described previously

(2). The cell lysate was stirred for 30 min at 4°C followed by centrifugation at 100,000 g for 30 min at 4°C. The supernatant was loaded onto 10 ml of Strep-Tactin resin (iba-Lifesciences) pre-equilibrated in lysis buffer. After loading the lysate with gravity flow, the resin was washed with 100 ml of lysis buffer followed by 100 ml of high salt buffer, A+1M (50 mM Hepes-KOH pH7.6, 1 M sodium chloride, 10% glycerol, 0.01% NP-40, and 5 mM β-mercaptoethanol). The resin was further washed with 100 ml of buffer A+400, then equilibrated with 50 ml of buffer A+200. The SEP tag was removed by on-column digestion with GST-3C protease for 2 hours at 4°C. After protease digestion, the resin was washed extensively with 100 ml of buffer A+200. Recombinant RDR2 was then eluted from Strep-Tactin resin with elution buffer (50 mM Hepes-KOH pH7.6, 200 mM sodium chloride, 5% glycerol, 0.01% NP-40, and 5 mM β-mercaptoethanol, and 13 mM desthiobiotin). Elutions were pooled, and the buffer was exchanged with A+100 using a desalting column (Bio-Rad). The resulting sample was loaded onto a 1 ml Hitrap Q column pre-equilibrated with buffer A+100. The column was then washed with 5 ml of A+100. Proteins were eluted with a 20 column volume (20 ml) linear gradient from A+100 to A+1000. Fractions containing recombinant full-length RDR2 were identified by SDS-PAGE and pooled. The buffer was exchanged with the storage buffer: 50 mM Hepes-KOH pH7.6, 200 mM sodium chloride, 5% glycerol, and 5 mM β-mercaptoethanol, and concentrated using a spin column (100kD cut-off) to a final concentration of 4.8 mg/ml, as measured using a Bradford assay.

**Cryo-EM sample preparation and data collection**

For cryo-electron microscopy, the grid preparation, grid screening and data collection were performed in the CryoEM facility at the HHMI Janelia Research Campus. To overcome preferred orientation issue, two different datasets were collected with no stage tilt and 30° stage tilt, respectively. For the un-tilted data set, 200 mesh Quantifoil R1.2/1.3 grids were glow-discharged for 1 minute with a current of 15mA in a PELCO easiGlow system before being mounted onto a Mark IV Vitrobot (FEI/Thermo Fisher Scientific). The sample chamber on the Vitrobot was kept at 4 °C with a relative humidity of 100%. 2.5 μl RDR2 sample at a concentration of 0.6 mg/ml was applied to the grid, which was then blotted from both sides for 1 second with blot force set at 3. After blotting the grid was rapidly plunge-frozen into a liquid ethane bath cooled by liquid nitrogen.

Single particle data was collected on the Janelia Krios1 microscope operated at 300 kV. The microscope is equipped with a spherical aberration corrector, an energy filter (Gatan GIF Quantum) and a post-GIF Gatan K3 direct electron detector. Movies were taken on the K3 camera in CDS mode at a calibrated magnification of ×59,242, corresponding to 0.844 Å per physical pixel (0.422 Å per super-resolution pixel). The dose rate on the specimen was set to be 15 electrons per $Å^2$ per second, and total exposure time was 4 s, resulting in a total dose of 60 electrons per $Å^2$. With dose fractionation set at 0.08 s per frame, each movie series contained 50 frames and each frame received a dose of 1.2 electrons per $Å^2$. An energy slit with a width of 20 eV was used during data collection. Automated data collection was carried out using SerialEM with a nominal defocus range set from −0.8 to −2.0 μm. Camera gain reference map was taken at the start of the data collection session but not applied to each movie series to limit file size. To further save disk space, each movie series was saved in tiff format with LZW compression. A total of 6,417 movies were acquired for the un-tilted data set. For the tilted data set, 3 μl aliquots of 0.46 mg/ml RDR2 was applied to glow-discharged 300 mesh UltrAUFoil R1.2/1.3 grids and the grids were prepared in the same manner and imaged on the same 300 keV Titan Krios cryo-electron microscope as described above. Movies were collected with the stage tilted at 30°. The imaging parameters for the tilted data collection are the same as those for the un-tilted data collection as described above. A total of 4,545 movies were acquired at 30° tilt.

**Cryo-EM data processing by RELION v3.1: Preprocessing and template-free 2D reference generation**

Image processing and single particle analysis were performed in RELION 3.1 (3, 4) through three stages: preprocessing & 2D reference generation (Fig. S1A), 3D reference map generation (Fig. S1B) and the final reconstruction for structural determination (Fig. S2). The flowchart and graphical summary of each stage are described in Fig. S1-S2. For all 3D classification/auto-refine jobs, unless otherwise stated, 3D references were low-pass filtered to 40 Å. No symmetry was used at any point (C1 symmetry). All resolution of a reconstruction is estimated by gold-standard Fourier Shell Correlation.

Motion corrections of the movies were done by RELION's implementation and were binned 2×, resulting in dose-weighted micrographs in 0.844 Å/pixel. Contrast Transfer Function (CTF) estimation was done by CtfFind-4.1.14 (5). To remove suboptimal micrographs, metadata filtering by rlnCtfMaxResolution < 5.2 Å and for un-tilted and < 10 Å for tilted micrographs. For the +30° tilted micrographs, additional filtering by rlnCtfAstigmatism < 2250 was employed. Finally, some micrographs with trajectory warning, severe ice chunk and noises were removed manually. The filtered 5,011 and 3,907 micrographs for the un-tilted and the tilted data were used in the subsequent reconstruction. 15 micrographs were manually selected, and 5,188 and 9,069 particles were initially picked by the template-free, Laplacian of Gaussian method for the un-tilted and the tilted data. Particles were extracted in 320 pixel box and 4× down-scaled (3.376 Å/pixel, 80-pixel box). Two rounds of 2D classification with a 128 Å spherical mask yielded three and eleven 2D class average images, which were chosen as 2D references for auto-pick from all micrographs. See Figure S1A for the flowchart.

**Cryo-EM data processing by RELION v3.1: 3D reference generation**

Using the 2D class averages, which were low-pass filtered to 20 Å as the 2D reference, total 1,172,128 and 3,055,625 particles were auto-picked from the filtered micrographs of the un-tilted and the tilted dataset, respectively. The particles were extracted, 4× down-scaled (3.376 Å/pixel, 80-pixel box) and separately subjected to two rounds of 2D classification with a 128 Å spherical mask, which resulted in selection of 16 classes, 348,938 particles and 30 classes, 622,587 particles for the un-tilted and the tiled dataset, respectively. Then, the particles were combined (971,525 particles) and used to produce a de novo 3D initial model. A single round of 3D auto-refine was performed, followed by a 3D classification (tau2_fudge = 1, E-step limit = 7) where two most resolved classes were selected (344,123 particles). 2× down-scaled particles (1.688 Å/pixel, 160-pixel box) were re-extracted, and used to 3D auto-refine the most resolved map (class006) from the prior 3D classification. After a few rounds of 3D auto-refine, the wrong handedness of the map was realized, therefore it went back to the class006 map and the handedness of the map was inverted. The corrected map and the 2× down-scaled particles were then subjected to four cycles of 3D auto-refine, followed by CTF refinement (per-particle defocus), and followed by Bayesian polishing of the particles. All masks were produced from maps low-pass filtered to 15 Å, unless otherwise stated. Using a mask (6-pixel soft edge), the polished particles are 3D auto-refined to ~4.2 Å and subjected to another round of 3D classification (tau2_fudge=4 with a mask, 5-pixel soft edge). Particles from two most resolved classes were combined, re-extracted unbinned (0.844 Å/pixel, 320-pixel box), 3D-autorefined and CTF refined (per-particle defocus, beam-tilt), then 3D auto-refined to ~4.3 Å. This map was selected as the 3D reference for auto-picking particles in the next stage. See Figure S1B for the flowchart.

**Cryo-EM data processing by RELION v3.1: Structural determination**

The flowchart of the final reconstruction for structural determination is described in Figure S2. The ~4.3 Å map was low-pass filtered to 20 Å and used as the 3D reference to auto-pick 2,022,652 and 3,170,634 particles from the 5,011 un-tilted and the 3,907 tilted micrographs, respectively. Particles were extracted to 4× down-scaled (3.376 Å/pixel, 80-pixel box) and separately subjected to two rounds of 2D classification with a 128 Å spherical mask. After removing obvious junk classes, 48 classes and 889,406 particles for the un-tilted, and 73 classes and 1,592,397 particles for the tilted, were selected and combined (2,491,803 particles). Then, the combined particles were subjected to two rounds of 3D classification (tau2_fudge = 1, E-step limit = 7, then tau2 fudge = 2, E-step limit = 7). After discarding particles belonging to suboptimal classes, 669,385 particles were selected and re-extracted to 1.688 Å/pixel, 160-pixel box. The 2× down-scaled particles and the map of the most resolved class were then 3D auto-refined to ~4.50 Å.

Then, optics_groups of "un-tilted" and "tilted" were assigned to corresponding particles and subjected to two cycles of CTF refine (per-particle defocus and per micrograph astigmatism) and 3D auto-refine. The particles were further CTF refined for anisotropic magnification followed by 3D auto-refine, then CTF refined again (per-particle defocus and per micrograph astigmatism), resulting in a ~3.80 Å resolution reconstruction. Additional round of 3D classification (tau2_fudge = 4 with a mask, 6-pixel soft edge) removed a class of suboptimal particles, then the remaining 572,690 particles were re-extracted to unbinned (0.844 Å/pixel, 320-pixel box) and subjected to 3D auto-refine with the most resolved 3D class (class004) map. The particles were further CTF refined for per particle defocus and per micrograph astigmatism, then 3D auto-refined and CTF refined again for beam-tilt, trefoil, 4th order aberrations, followed by per-particle defocus and per-micrograph astigmatism refinement. After 3D auto-refine, the particles were subjected to Bayesian polishing, resulting in a ~3.70 Å resolution reconstruction after subsequent 3D auto-refine. To improve the CTF refinement, additional optics groups were assigned to particles based on their 3×3 image-shift positions (9 optics_groups each for the untilted and the tilted datasets, in total 18 optics_groups) according to the REILION FAQs. At this point, 915 particles belonged to an irregular acquisition target were manually removed. The remaining 571,775 particles were CTF refined for anisotropic magnification, beam-tilt, and per-particle defocus and per-micrograph astigmatism. Subsequent 3D auto-refine yielded a ~3.46 Å resolution reconstruction. Four more rounds of the CTF refinement and 3D auto-refine resulted in ~3.33 Å.

To remove suboptimal particles further, iterative metadata filtering (MaxValueProbDistribution and NrOfSignificantSamples) was performed. After removing particles with a cutoff value of 25% (MaxValueProbDistribution lower than 25th percentile; NrOfSignificantSamples higher than 75th percentile), the remaining 405,432 particles were 3D auto-refined to ~3.29 Å resolution. Subsequently, two rounds of CTF refinement (anisotropic magnification, per-particle defocus, beam tilt, higher-order aberrations) and 3D auto-refine, yielded a 3.22 Å reconstruction. After 5% cutoff for MaxValueProbDistribution and NrOfSignificantSamples, additional two rounds of CTF refinement followed by 3D auto-refine and Bayesian polishing with re-centering, the 374,967 particles were 3D auto-refined to ~3.18 Å. Then, particles were filtered by 10% cutoff on the metadata and the 320,243 particles were CTF refined and 3D auto-refined again. Additional particle filtering at 5% cutoff resulted in 293,561 particles, which were then subjected to 3D classification with a mask (3-pixel extension, 6-pixel soft edge) and skipping alignment. The most resolved class with 128,988 particles was selected, 3D auto-refined to 3.14 Å, metadata filtered at 5% cutoff and CTF refined for anisotropic magnification, per-particle defocus, per micrograph astigmatism, beam tilt and higher order aberrations.

Finally, the resulting 118,561 particles were 3D auto-refined using a mask (3-pixel extension, 6-pixel soft edge) with SIDESPLITTER (6) to suppress local overfitting, resulting in a 3.10 Å resolution reconstruction, which we termed “Map1”. The directional resolution anisotropy was evaluated by 3DFSC (7), showing the sphericity of 0.949 out of 1 (Fig. S3B). The local resolution estimation was done by the Local resolution job with relion_postprocess (Fig. S3C). The angular distribution was assessed by displaying the_angdist.bild file output (Fig. S3D). The map was sharpened by the DeepEMhancer (8) and used to build a model (see the Model building section). With the model, the pixel size of the final reconstruction was calibrated and determined to be 0.847 Å/pixel using the method described previously (9), and was used to make the final gold-standard FSC plot in Fig. S3A.

**Cryo-EM data processing by cryoSPARC v3.2**

The full cryoEM data processing workflow is illustrated in Fig. S4. Motion correction, CTF-estimation, particle picking, 2D classification, *Ab-initio* 3D reconstruction, and non-uniform 3D refinement were performed in cryoSPARC v3.2 (10). Both tilted and non-titled raw datasets were processed separately up to 2D classification. Particles chosen by 2D classification from each dataset were combined. Additional round of 2D classification was then performed using these combined particles followed by *Ab-initio* 3D reconstruction, which generated 5 different initial 3D maps. The best map among 5 maps was chosen. Particles in the best map were further subjected to 2$^{nd}$ of *Ab-initio* 3D reconstruction, generating 3 maps. Two of which were subjected to heterogeneous 3D refinement. The best map at 4.20Å resolution from heterogeneous refinement was further refined by non-uniform 3D refinement with CTF and defocus corrections, yielding the map at 3.63 Å resolution. Then this map was further refined by Non-uniform 3D refinement with beam title corrections, yielding the map, termed "Map2" at 3.57 Å resolution (Figure S5) estimated from Fourier shell correlation (FSC) curves calculated using the gold-standard procedure with the 0.143 cut-off criteria (11-13). 3D Fourier shell correlation analysis of the map was carried out using 3DFCS program (7) indicating the sphericity of Map2 is 0.93 out of 1.0 (Figure S6B). The map was subjected to 3D variability analysis (3DVA) via cryoSPARC v3.2 (Movie S1) (14). The map was sharpened by the program DeepEMhancer (8) via COSMIC$^2$ site (15). Visualization of EM maps was carried out by UCSF Chimera (16).

#### Model building and refinement

Map1 was sharpened using the DeepEMhancer tool (8) of the COSMIC[2] science gateway (15) then used for model building. Model building for amino acid residues 61-1121) of Map1 was carried out *de novo* using the automated model building program Buccaneer (17). After one round of automated model building by Buccaneer, the model was inspected and further built manually in Coot (18). The resulting model was then subjected to another round of automated model building by Buccaneer and the process repeated until all interpretable EM densities were filled with corresponding amino acid residues. *De novo* model building for amino acids 1-60 using Buccaneer was not possible due to relatively low quality EM density in this area as indicated by the local resolution (Fig. S2C). Instead, a structural model for RDR2 region aa 1-100 was generated by the protein structure predictor, I-TASSER (19). The predicted structure was then fitted into the *de novo* model for amino acids 61-1121 followed by several iterations of manual building in Coot (18) and refinement by Phenix (20). The fully built model was iteratively refined using Real Space Refinement in Phenix (20) followed by manual inspection and refinement in Coot. The model for the EM map from cryoSPARC (Map2) was generated by fitting the Map1 model into Map 2 using rigid body refinement by REFMAC (21) followed by manual refinement in Coot and Real Space Refinement in Phenix (20). Model refinement statistics are provided in Table S1.

#### RDR2 Transcription Assays

RDR2 transcription reactions in Fig. 1B were carried out in 10-µl reactions containing 90-4800 ng of RDR2, 170 nM 37 nt template RNA, 25 mM HEPES-KOH pH 7.9, 20 mM ammonium acetate, 2 mM MgCl2, 0.1 mM EDTA, 0.01% Triton X-100, 3% PEG-8000, 0.1 mM each of ATP, GTP and CTP, 0.4 U/µl RNase Inhibitor Murine (NEB, M0314), at 25 °C for 1 hour. The template RNA was 5'-end-labeled using T4 polynucleotide kinase (NEB) and gamma-[$^{32}$P]-ATP (Perkin Elmer) prior to the reaction. An equal volume of loading buffer containing 1xTBE, 4 mM MgCl2, 60% glycerol, 0.02% bromophenol blue and 0.02% xylene cyanol was then added and RNAs were resolved by native PAGE on a 15% polyacrylamide gel. The gel was transferred to filter paper and imaged using a Typhoon FLA 9500 phosphorimager.

Reactions of Fig. 4A were conducted as described above but included 250 ng RDR2, 0.1 mM each of ATP, GTP, CTP and 2.5 μCi alpha-[$^{32}$P]-GTP. Reactions of Fig. 4B included 125 ng RDR2, 0.1 mM each of ATP and GTP, 0.1 mM 2'3'-dideoxy-CTP in some reactions, and 2.5 μCi alpha-[$^{32}$P]-CTP or gamma-[$^{32}$P]-ATP. Reactions were incubated at room temperature for 1 hour. Reactions were stopped by adding 5 volumes of Proteinase K solution (100 mM Tris-HCl pH 7.9, 250 mM NaCl, 1 mM MgCl2, 1% SDS, 0.8 mg/ml Proteinase K, 0.6~1.0 mg/ml GlycoBlueTM (Thermo Fisher AM9515)), and incubation at 37 °C for 30 min. Following ethanol precipitation, pellets were washed with 70% ethanol then was resuspended in Novex™ TBE-Urea Sample Buffer (LC6876), heated at 72 °C for 3 min, chilled on ice, then resolved by denaturing PAGE on 17% polyacrylamide 7M Urea gels. Gels were vacuum-dried and subjected to phosphorimaging.

For transcription reactions using RNA-DNA hybrids (Fig. 5), equal molar amounts of template RNA and DNA oligos were mixed in hybridization buffer (25 mM HEPES-KOH pH7.9, 20 mM ammonium acetate), incubated at 95 °C in a heat block for 5 min, an additional 10 min with the heat turned off, and an additional 10 min at room temperature. Transcription reactions were carried out in 10-μl reactions containing 290 ng RDR2, 100 nM template RNA-DNA hybrid, 25 mM HEPES-KOH pH 7.9, 20 mM ammonium acetate, 2 mM MgCl2, 0.1 mM EDTA, 0.1 mM each of ATP, GTP and CTP, and 2.5 μCi of alpha-[$^{32}$P]-ATP. After incubating at 27 °C for 2 hours, reactions were stopped by adding EDTA to 5 mM, passed through PERFORMA spin columns (Edge Bio) and adjusted to 300 mM sodium acetate. 0.7 μl of GlycoBlueTM (Thermo Fisher AM9515) was added and products were precipitated with 3 volumes of isopropanol at -20 °C for overnight. After centrifuging at 16,100 rcf for 30 min, the pellet was washed with 70% ethanol, resuspended in Novex™ TBE-Urea Sample Buffer (LC6876), heated at 72 °C for 3 min, chilled on ice, and resolved by denaturing PAGE on 15% polyacrylamide 7M Urea gels. The gels were then vacuum-dried and subjected to phosphorimaging.

For assays involving template DNA, nontemplate DNA and RNA strands (Fig. 6), annealing of the 51 nt template DNA, 28 nt nontemplate DNA and 39 nt RNA was performed as described above. RDR2 transcription reactions were then performed using 250 nM of annealed nucleic acid mix and 200 ng of RDR2 in 40 μl reactions (22). Transcription reactions were passed through PERFORMA spin columns, precipitated, washed and subjected to denaturing PAGE on 15% polyacrylamide 7M Urea gels as described above. Gels were vacuum dried and subjected to autoradiography using Carestream® BioMax® X-ray film. For RNase I sensitivity assays, the 39 nt RNA was 5'-end labeled using T4 polynucleotide kinase (NEB) and 25 µCi of gamma-[$^{32}$P]-ATP (Perkin Elmer) prior to annealing. Annealed products were then subjected to RNase I (0.01~1.0 unit per 40 µl reaction) as previously described (23) and analyzed by denaturing PAGE as described above.

**Recombinant TF-RDR2-RRM protein expression in *E.coli* and purification**

To express and purify TF-RDR2-RRM proteins, a preculture of *E. coli* ArcticExpress cells with pCold-TF-RDR2-RRM vectors were incubated in LB media containing 50 µg/ml carbenicillin and 20 µg/ml gentamycin for overnight at 37 °C on a rotator drum. Then, the saturated cells were collected by centrifugation at 4,000 rcf for 5 min, then resuspended in the same pre-warmed media with fresh antibiotics in the same volume. The cells were 1/10$^{th}$ diluted in the same media and incubated at 37 °C, 225 rpm until the OD reaches around 0.6, then proceeded to induction by chilling the culture to 12-15 °C in an ice-water bath for 30 min. IPTG (final 0.1 mM) was added to the cold-induced culture and incubated at 12 °C, 200 rpm for about 24 hours. Cells were harvested by centrifugation at 4,000 rcf for 6 min, flash frozen in liquid nitrogen and stored at -80 °C.

The frozen cell pellet was resuspended in 1/22 th culture volume of lysis buffer containing 50 mM Tris-HCl pH 7.9, 300 mM NaCl and 50 µM $ZnSO_4$, 1 mM TCEP, 1 mM PMSF and 1x protease inhibitor cocktail (Sigma, P9599). Proteins were extracted by sonication, followed by addition of Triton X-100 (final 0.5%) and incubation at 4 °C for 30 min. Insoluble proteins were removed by centrifugation at 18,000 rcf, 4 °C for 30 min. The cleared soluble fraction was mixed with Ni-NTA agarose (Qiagen) equilibrated in the lysis buffer with 0.1% Triton X-100, and incubated batchwise at 4 °C. After at least an hour, the resin suspension was transferred into an empty gravity column and the lysate was allowed to flow through. The column was then washed with 20 column volumes (CV) of wash buffer containing 50 mM Tris-HCl pH 7.9, 300 mM NaCl and 50 µM $ZnSO_4$, 1 mM TCEP, 0.1% Triton X-100, 20 mM imidazole, 10% glycerol and 0.2 mM PMSF, followed by 20 CV of the wash buffer but with 150 mM NaCl and 0.01% Triton X100. Then, proteins were eluted from the column by a buffer containing 50 mM Tris-HCl pH 7.9, 150 mM NaCl and 50 μM $ZnSO_4$, 1 mM TCEP, 0.01% Triton X-100, 250 mM imidazole, 10% glycerol and 0.2 mM PMSF and fractionated into 1 CV each, after discarding the void volume (0.3 CV). The elution fraction 2 was used in the subsequent EMSA assay.

#### References cited in detailed methods

1. M. Z. Li, S. J. Elledge, Harnessing homologous recombination in vitro to generate recombinant DNA via SLIC. *Nat. Methods* **4**, 251-256 (2007).
2. T. Imasaki, S. Wenzel, K. Yamada, M. L. Bryant, Y. Takagi, Titer estimation for quality control (TEQC) method: A practical approach for optimal production of protein complexes using the baculovirus expression vector system. *PLoS One* **13**, e0195356 (2018).
3. J. Zivanov *et al.*, New tools for automated high-resolution cryo-EM structure determination in RELION-3. *eLife* **7**, e42166 (2018).
4. J. Zivanov, T. Nakane, S. H. W. Scheres, Estimation of high-order aberrations and anisotropic magnification from cryo-EM data sets in RELION-3.1. *IUCrJ* **7**, 253-267 (2020).
5. A. Rohou, N. Grigorieff, CTFFIND4: Fast and accurate defocus estimation from electron micrographs. *J. Struct. Biol.* **192**, 216-221 (2015).
6. K. Ramlaul, C. M. Palmer, T. Nakane, C. H. S. Aylett, Mitigating local over-fitting during single particle reconstruction with SIDESPLITTER. *J. Struct. Biol.* **211**, 107545 (2020).
7. Y. Z. Tan *et al.*, Addressing preferred specimen orientation in single-particle cryo-EM through tilting. *Nat. Methods* **14**, 793-796 (2017).
8. R. Sanchez-Garcia *et al.*, DeepEMhancer: a deep learning solution for cryo-EM volume post-processing. *Commun. Biol.* **4**, 874 (2021).
9. S. K. Natchiar, A. G. Myasnikov, H. Kratzat, I. Hazemann, B. P. Klaholz, Visualization of chemical modifications in the human 80S ribosome structure. *Nature* **551**, 472-477 (2017).
10. A. Punjani, J. L. Rubinstein, D. J. Fleet, M. A. Brubaker, cryoSPARC: algorithms for rapid unsupervised cryo-EM structure determination. *Nat. Methods* **14**, 290-296 (2017).
11. R. Henderson *et al.*, Outcome of the first electron microscopy validation task force meeting. *Structure* **20**, 205-214 (2012).
12. P. B. Rosenthal, R. Henderson, Optimal determination of particle orientation, absolute hand, and contrast loss in single-particle electron cryomicroscopy. *J. Mol. Biol.* **333**, 721-745 (2003).
13. S. H. Scheres, RELION: implementation of a Bayesian approach to cryo-EM structure determination. *J. Struct. Biol.* **180**, 519-530 (2012).
14. A. Punjani, D. J. Fleet, 3D variability analysis: Resolving continuous flexibility and discrete heterogeneity from single particle cryo-EM. *J. Struct. Biol.* **213**, 107702 (2021).
15. M. A. Cianfrocco, M. Wong, C. Youn, R. Wagner, A. E. Leschziner, COSMIC$^2$: A Science Gateway for Cryo-Electron Microscopy Structure Determination. *Practice & Experience in Advanced Research Computing* **Article 22**, 1-5 (2017).

16. E. Pettersen *et al.*, UCSF Chimera--a visualization system for exploratory research and analysis. *J. Comput. Chem.* **25**, 1605-1612 (2004).

17. K. Cowtan, The Buccaneer software for automated model building. 1. Tracing protein chains. *Acta Crystallogr. D Biol. Crystallogr.* **62**, 1002-1011 (2006).

18. P. Emsley, K. Cowtan, Coot: model-building tools for molecular graphics. *Acta Crystallogr. D Biol. Crystallogr.* **60**, 2126-2132 (2004).

19. J. Y. Yang *et al.*, The I-TASSER Suite: protein structure and function prediction. *Nat. Methods* 12, 7-8 (2015).

20. P. Adams *et al.*, PHENIX: a comprehensive Python-based system for macromolecular structure solution. *Acta Crystallogr. D Biol. Crystallogr.* **66**, 213-221 (2010).

21. G. N. Murshudov, A. A. Vagin, E. J. Dodson, Refinement of macromolecular structures by the maximum-likelihood method. *Acta Crystallogr. D Biol. Crystallogr.* **53**, 240-255 (1997).

22. V. Mishra *et al.*, Assembly of a dsRNA synthesizing complex: RNA-DEPENDENT RNA POLYMERASE 2 contacts the largest subunit of NUCLEAR RNA POLYMERASE IV. *Proc. Natl. Acad. Sci. U. S. A.* **118**, e2019276118 (2021).

23. J. Singh, V. Mishra, F. Wang, H. Y. Huang, C. S. Pikaard, Reaction Mechanisms of Pol IV, RDR2, and DCL3 Drive RNA Channeling in the siRNA-Directed DNA Methylation Pathway. *Mol. Cell* **75**, 576-589 e575 (2019).

**Table S1. Cryo-EM data collection, refinement and validation statistics**

| | | |
|---|---|---|
| **Data collection** | | |
| | **Un-titled dataset** | **Tilted dataset** |
| Magnification | ×59,242 | ×59,242 |
| Voltage (kV) | 300 | 300 |
| Electron exposure (e/Å$^2$) | 60 | 60 |
| Defocus range (μm) | -0.8 to -2.0 | -0.8 to -2.0 |
| Pixel size (Å) | 0.422 | 0.422 |
| **Data processing** | | |
| | **Map 1 (EMD-24610, PDB 7ROZ)** | **Map 2 (EMD-24635, PDB 7RQS)** |
| Symmetry imposed | C1 | C1 |
| Initial particle images (no.) | 5,193,286<br>un-tilted: 2,022,652<br>tilted: 3,170,634 | 1,340,940<br>un-tilted: 287,145<br>tilted: 1,053,795 |
| Final particle images (no.) | 118,561 | 277,019 |
| Map resolution (Å) | 3.104 / 3.115* | 3.57 |
| FSC threshold | 0.143 | 0.143 |
| Map resolution range | 3.1-4.3 | 2.8-5.2 |
| Map sharpening B factor (Å$^2$) | -73.71 / -74.24* | -173.2 |
| **Refinement** | | |
| Model resolution (Å$^2$) | 2.2 | 2.3 |
| Model composition | | |
| Non-hydrogen atoms | 7756 | 7792 |
| Protein residues | 1064 | 1074 |
| Ligand (Mg) | 1 | 1 |
| B-factors (min/max/mean) | | |
| Protein | 32.27/105.16/56.28 | 22.41/145.88/57.78 |
| Ligand | 39.21/39.21/39.21 | 35.87/35.87/35.87 |
| r.m.s.d. deviations | | |
| Bond length (Å) | 0.003 | 0.002 |
| Bond angles (° ) | 0.543 | 0.517 |
| Validation | | |
| MolProbility score | 2.01 | 1.99 |
| Clashscore | 11.4 | 9.94 |
| Rotamer outliers (%) | 0 | 0 |
| CaBLAM outliers (%) | 5.33 | 4.44 |
| Ramachandran plot (%) | | |
| Outliers | 0 | 0 |
| Allowed | 6.78 | 7.49 |
| Favored | 93.22 | 92.51 |

* Values with the model-calibrated pixel size (0.847 Å/pixel)

#### Table S2. Oligo nucleotide sequences used in the study

| Figure used | Oligo Name | Oligo sequence (5'-3') |
|---|---|---|
| 1B | 37rUGC17G3ddC | rUrCrCrGrUrCrCrGrUrCrUrGrCrUrGrUrGrGrGrUrCrUrCrCrUrUrUrCrUrCrCrU rUrCrUrU/3ddC/ |
| 3B | A647_37rUGC17G3ddC | /5Alex647N/rUrCrCrGrUrCrCrGrUrCrUrGrCrUrGrUrGrGrGrUrCrUrCrCrUrUr UrCrUrCrCrUrUrCrUrU/3ddC/ |
| 4A | Temp4s_5 | rUrCrUrUrC |
| | Temp4s_7 | rCrUrUrCrUrUrC |
| | Temp4s_9 | rUrCrCrUrUrCrUrUrC |
| | Temp4s_11 | rUrGrUrCrCrUrUrCrUrUrC |
| | Temp4s_13 | rUrUrUrGrUrCrCrUrUrCrUrUrC |
| | Temp4s_15 | rCrCrUrUrUrGrUrCrCrUrUrCrUrUrC |
| | Temp4s_17 | rGrUrCrCrUrUrUrGrUrCrCrUrUrCrUrUrC |
| 4B | Temp4s_15 | rCrCrUrUrUrGrUrCrCrUrUrCrUrUrC |
| | Temp4s_15-15U | rCrCrUrUrUrGrUrCrCrUrUrCrUrUrU |
| | Temp4s_15-14C | rCrCrUrUrUrGrUrCrCrUrUrCrUrCrC |
| 5 | RDR2RNATemp4 | rUrGrCrGrUrGrCrGrUrCrUrGrCrGrUrCrGrUrUrCrGrUrCrCrUrUrUrGrUrCrCrU rUrCrUrU/3ddC/ |
| | R2DNATemp4rev | GAAGAAGGACAAAGGACGAACGACGCAGACGCACGCA |
| | R2DNATemp4rev3'mis2 | ACAGAAGGACAAAGGACGAACGACGCAGACGCACGCA |
| | R2DNATemp4rev3'mis4 | ACCAAAGGACAAAGGACGAACGACGCAGACGCACGCA |
| | R2DNATemp4rev3'mis9 | ACCACGAACCAAAGGACGAACGACGCAGACGCACGCA |
| | R2DNATemp4rev3'mis12 | ACCACGAACACCAGGACGAACGACGCAGACGCACGCA |
| | R2DNATemp4rev3'mis16 | ACCACGAACACCGAAGCGAACGACGCAGACGCACGCA |
| | R2DNATemp4rev3'Δ16 | CGAACGACGCAGACGCACGCA |
| 6A-C | First Strand RNA complete (39nt) | rCrGrUrGrUrCrGrGrUrCrCrUrGrGrCrGrUrUrCrUrCrUrGrUrCrUrGrCrUrUrUrC rGrUrUrGrUrCrU |
| | T-Less-Template: | CAAAAACGAGACAGACAACGAAAGCAGACAGAGAACGCCAGGACCGACACG |
| | (15 bp Overlap) T-less-Non Template (28nt): | GCTGCTTTCGTTGTCTGTCTCGTTTTTG |
| | (10 bp overlap) 5nt Mismatch_T-less Non template (28nt): | GAAAAATTCGTTGTCTGTCTCGTTTTTG |
| | (5 bp overlap) 10nt Mismatch_T-less Non template (28nt): | GAAAAAAAAAATGTCTGTCTCGTTTTTG |
| | (0 bp overlap) 15nt Mismatch_T-less Non Template (28nt): | GAAAAAAAAAAAAAAAGTCTCGTTTTTG |

**Table S3. Multi-subunit RNAP structures used for structural comparisons**

| Figure | PDB used | Structure | Reference |
|---|---|---|---|
| Figure 2A-D, Figure S8 | 2E2H | *Saccharomyces cerevisiae*, RNA polymerase II elongation complex at 5 mM Mg2+ with GTP | Wang, D. et al. 2006, Cell 127: 941-954 |
| Figure S15 | 2E2I | *Saccharomyces cerevisiae*, RNA polymerase II elongation complex in 5 mM Mg+2 with 2'-dGTP (Fork loop 2 visible) | Wang, D. et al. 2006, Cell 127: 941-954 |
| Figure 2A-D, Figure S16 | 4Q4Z | *Thermus thermophilus* RNA polymerase de novo transcription initiation complex | Basu, R.S. et al. 2014, J Biol Chem 289: 24549-24559 |

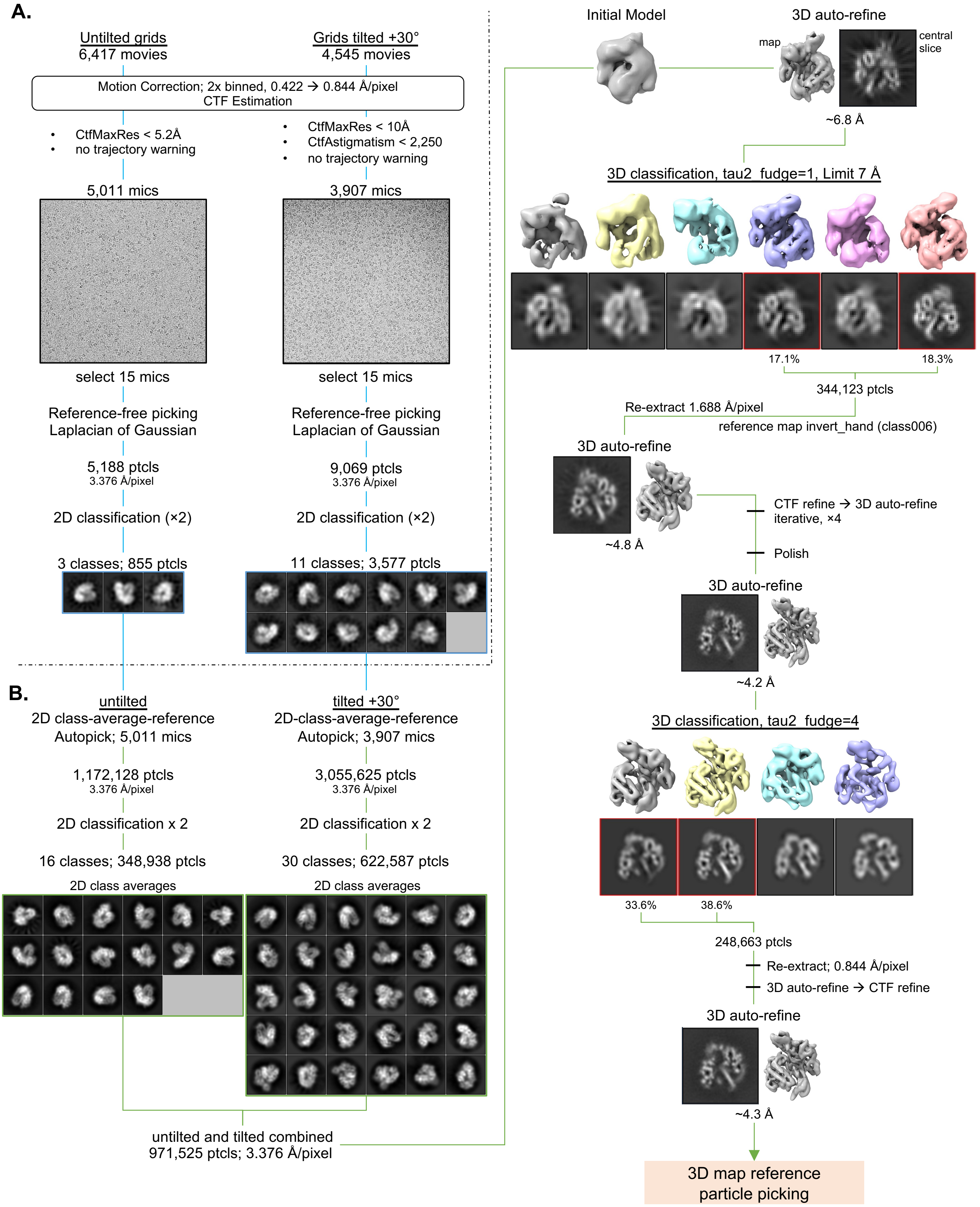

**Figure S1. Overview of cryo-EM data processing using Relion 3.1: Generation of 2D and 3D references toward structural determination**

(A) Steps involved in preprocessing and template-free 2D reference generation. (B) Steps involved in 3D reference generation. See supplemental methods for details.

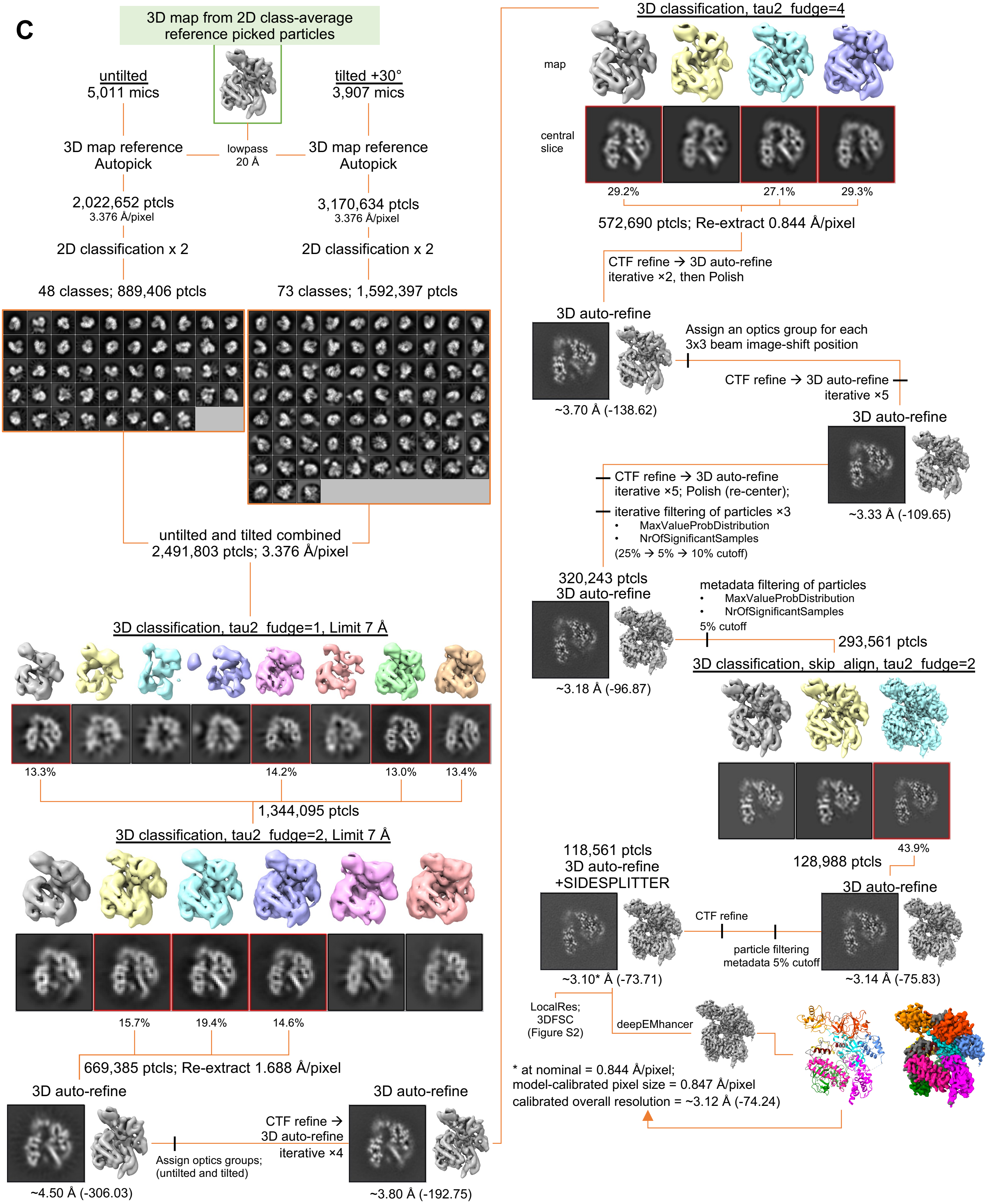

**Figure S2. Overview of cryo-EM data processing using Relion 3.1: Final reconstruction for structural determination**

Steps involved in the final reconstruction for structural determination. Numbers in parentheses indicate the b-factor estimated by relion_postprocess. See supplemental methods for details.

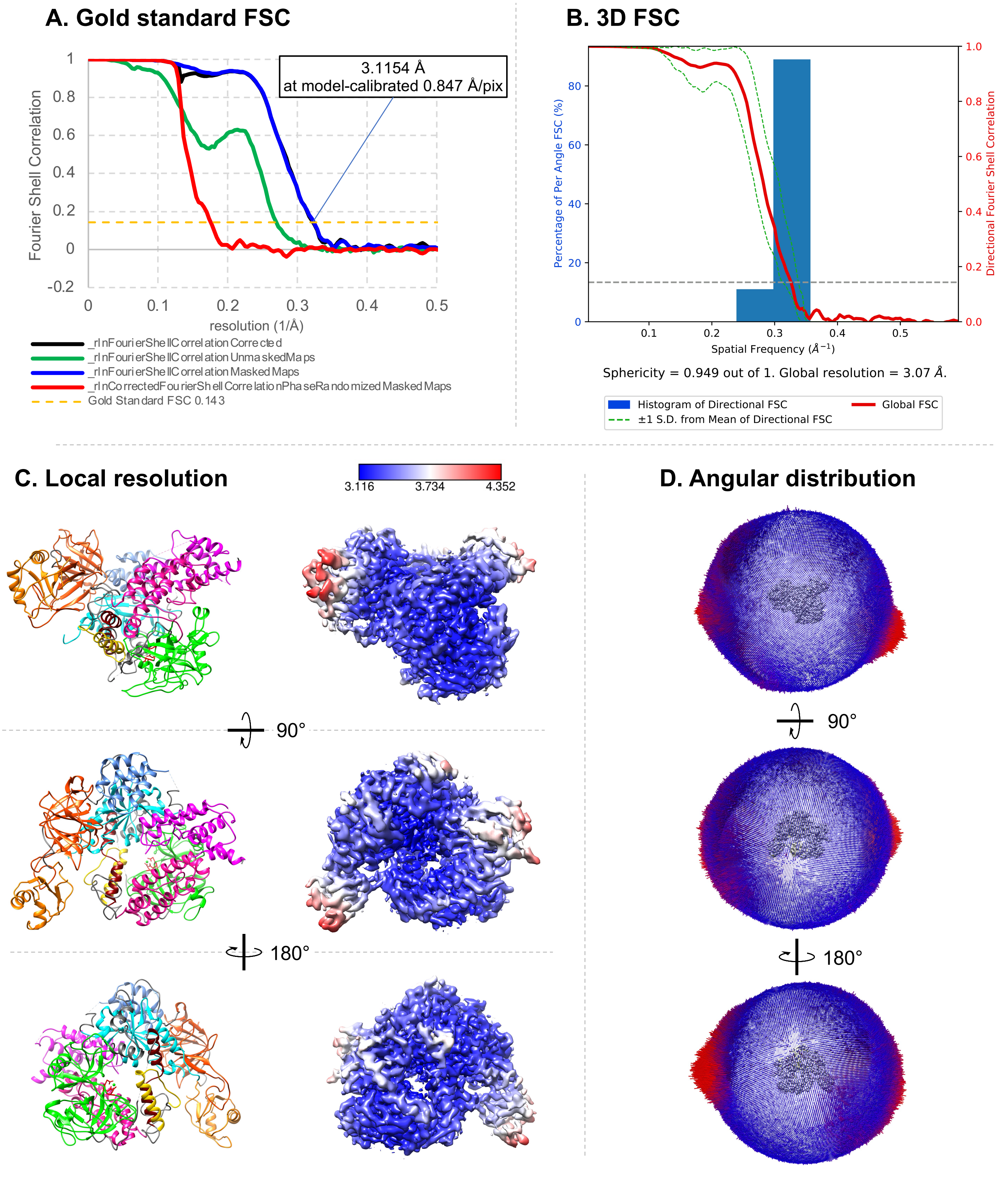

**Figure S3. RDR2 cryo-EM-Relion map and model quality estimations**
(A) Gold-standard Fourier Shell Correlation plot for the final reconstruction, using a model-calibrated pixel size of 0.847 Å/pixel. (B) 3D-FSC plot and sphericity (0.949) for the final reconstruction. (C) Local resolution estimated by relion_postprocess. (D) Angular distribution for the final reconstruction. See supplemental methods for details.

**A. Model and map comparison**

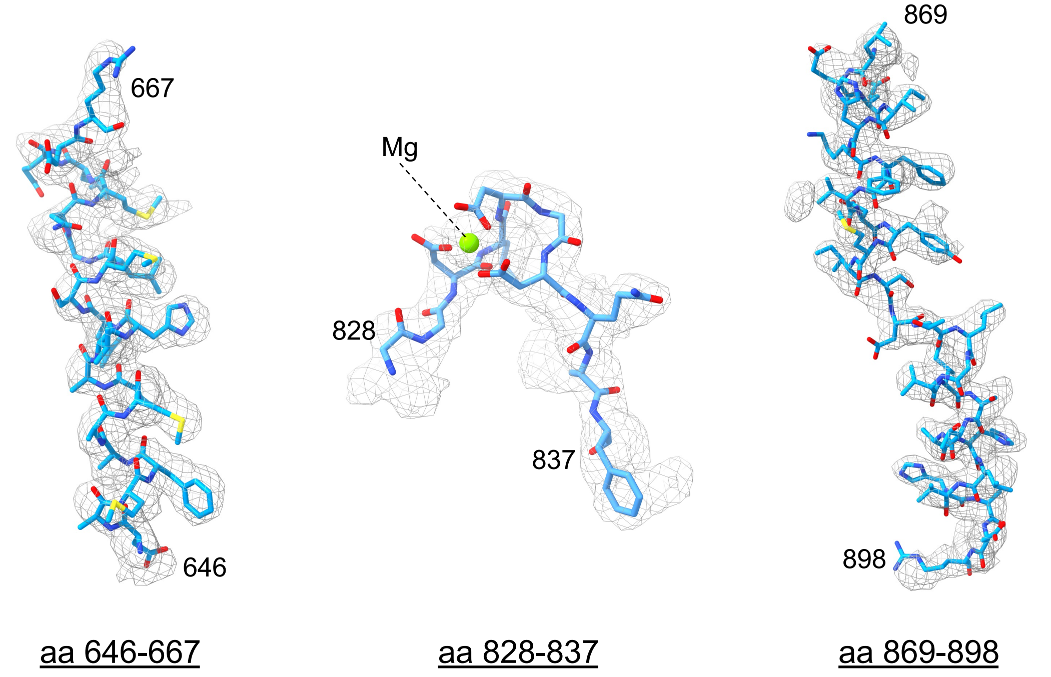

**B. Map-model FSC curve**

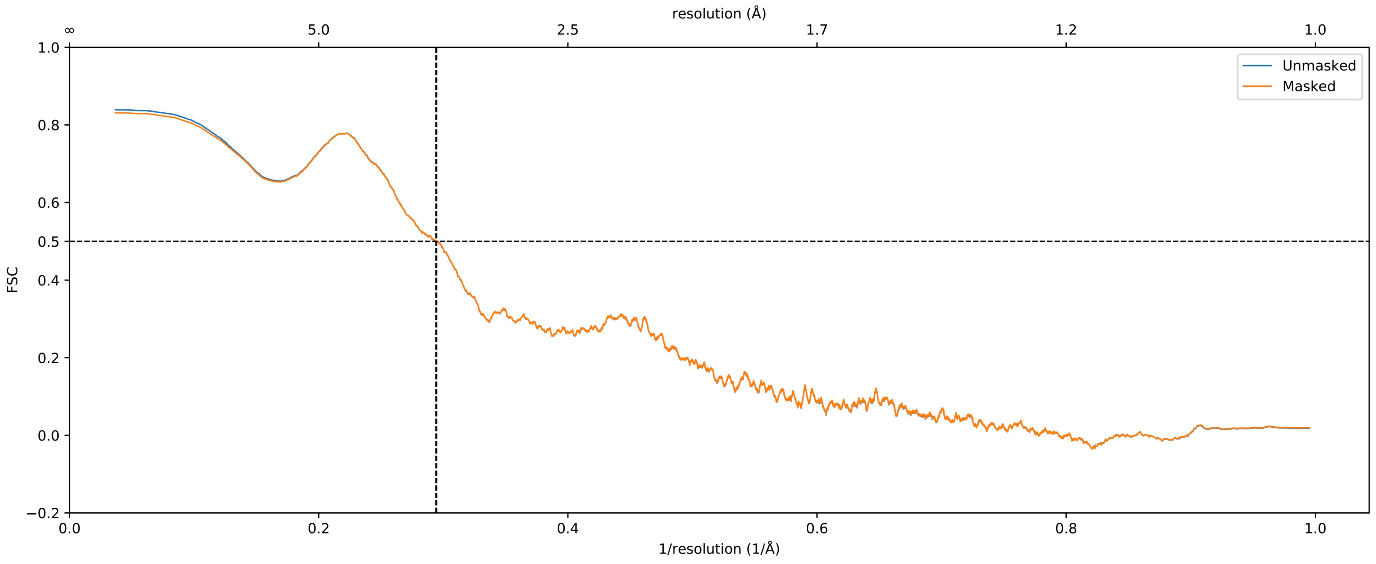

**Figure S4. Validation of cryo-EM structure Map1 and its model**
(A) Representative density of Map1 regions including the Neck1 domain (aa 646-667), active site loop (aa 828-837), and the bridge helix. (B) Model-to-map FSC curves generated in Phenix.

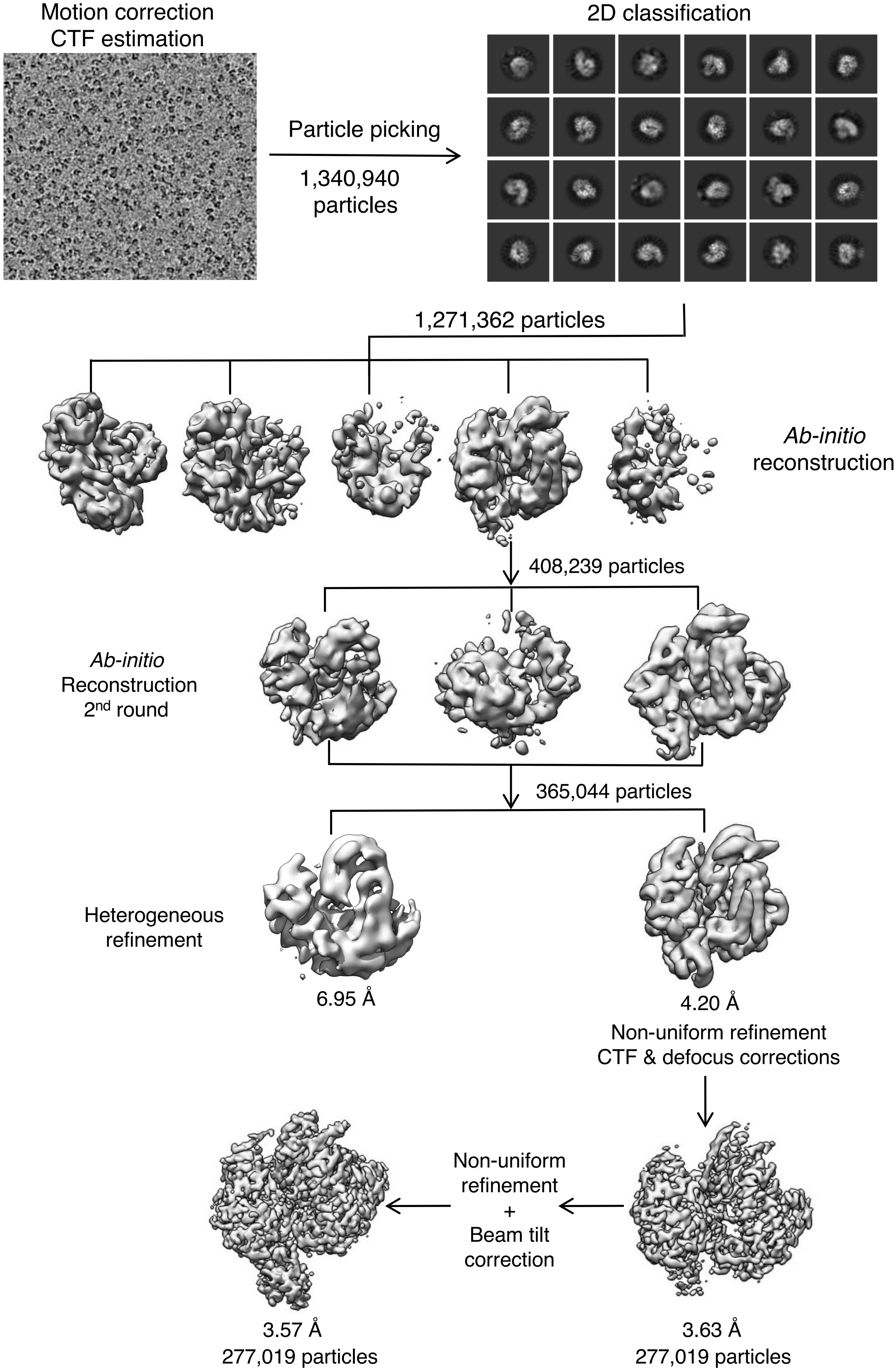

**Figure S5. Summary of cryo-EM data processing workflow using cryoSPARC v3.2 to generate Map2**

#### A. Gold standard FSC curve

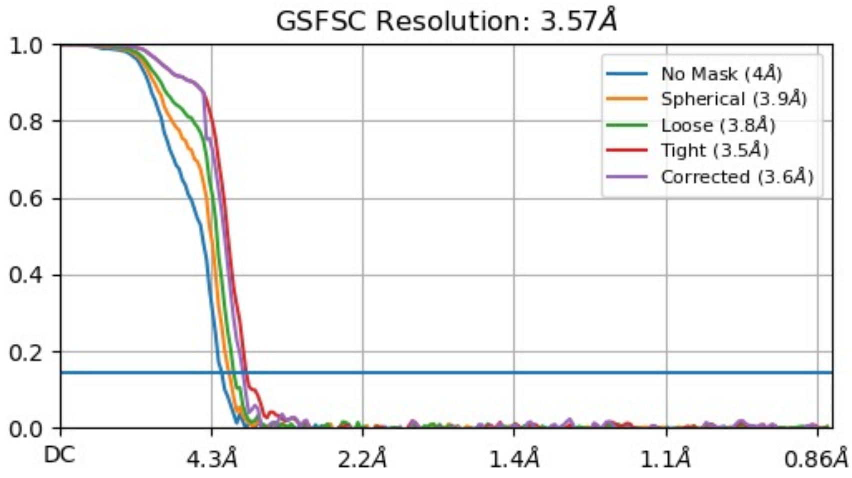

#### C. Viewing distribution plots

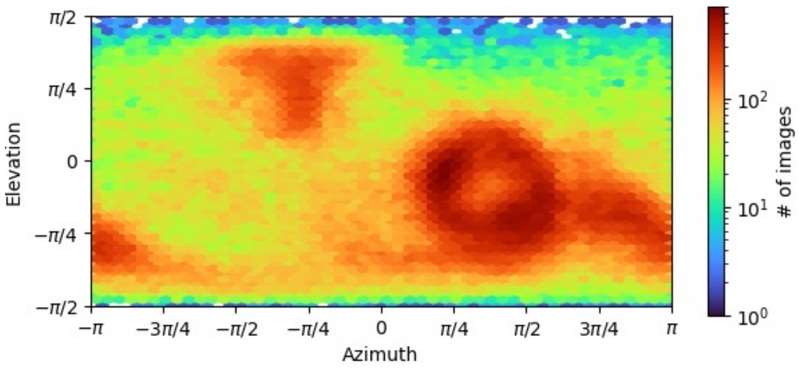

#### B. 3D FSC

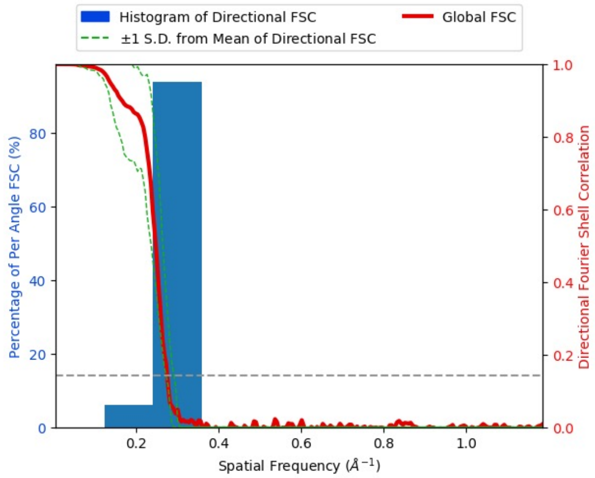

Sphericity: 0.93 out of 1.0 Global resolution 3.66 Å

#### D. Local resolution

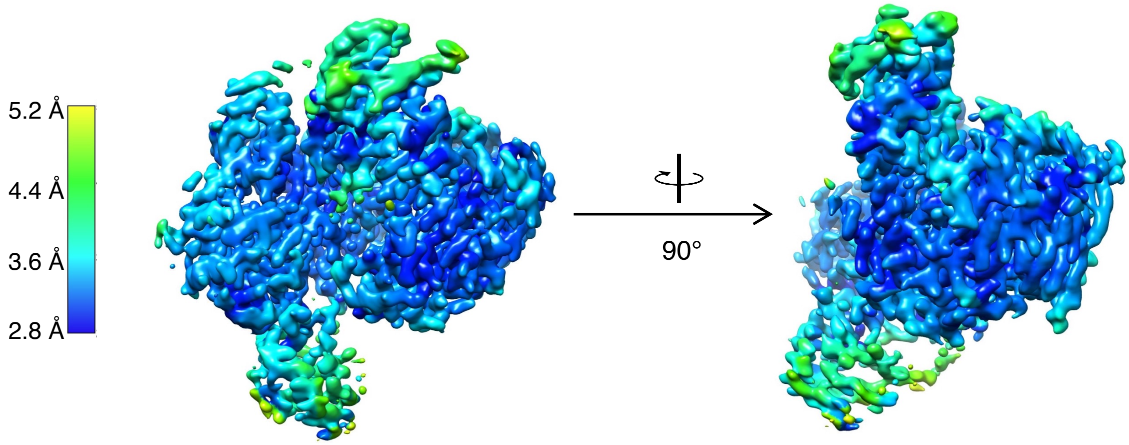

**Figure S6. Assessment of quality of RDR2 cryo-EM Map2**

(A) Gold standard FSC curve for Map2 of RDR2 generated in cryoSPARC v3.2 .
(B) 3DFSC: histogram and directional FSC plot for Map2 Sphericity and global resolution are indicated.
(C) Viewing distribution plots of Map2 generated in cryoSPARC v3.2.
(D) Local resolution of Map2 colored according to the local resolution (scale is at left).

#### A. Representative densities and corresponding models

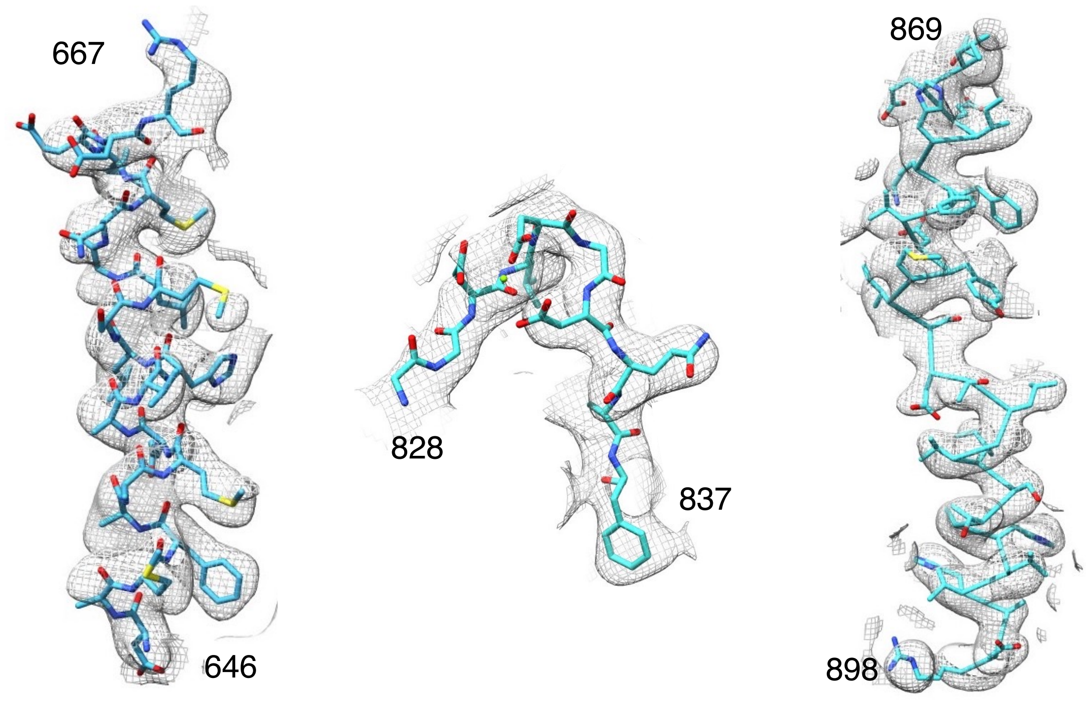

aa 646-667 aa 828-837 aa 869-898

#### B. Map-model FSC curve

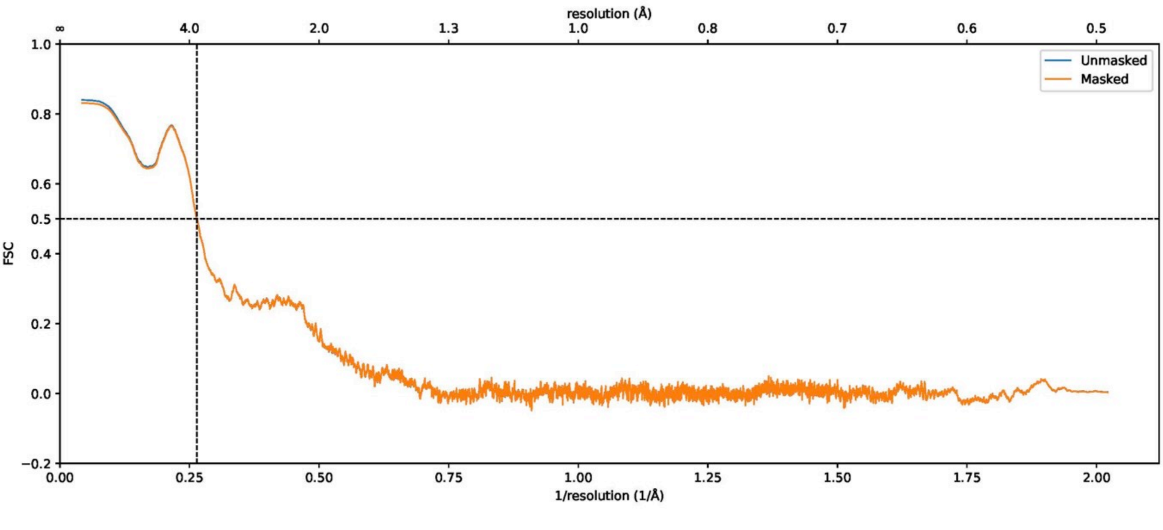

**Figure S7. Validation of cryo-EM structure: Map2 and its model**

(A) Representative densities of Map2 for the Neck1 domain (aa 646-667), active site loop (aa 828-837), and Bridge helix (BH) (aa 869-898). The amino acid residues encompassing each density are shown.

(B) Model-to-map FSC curves generated in Phenix.

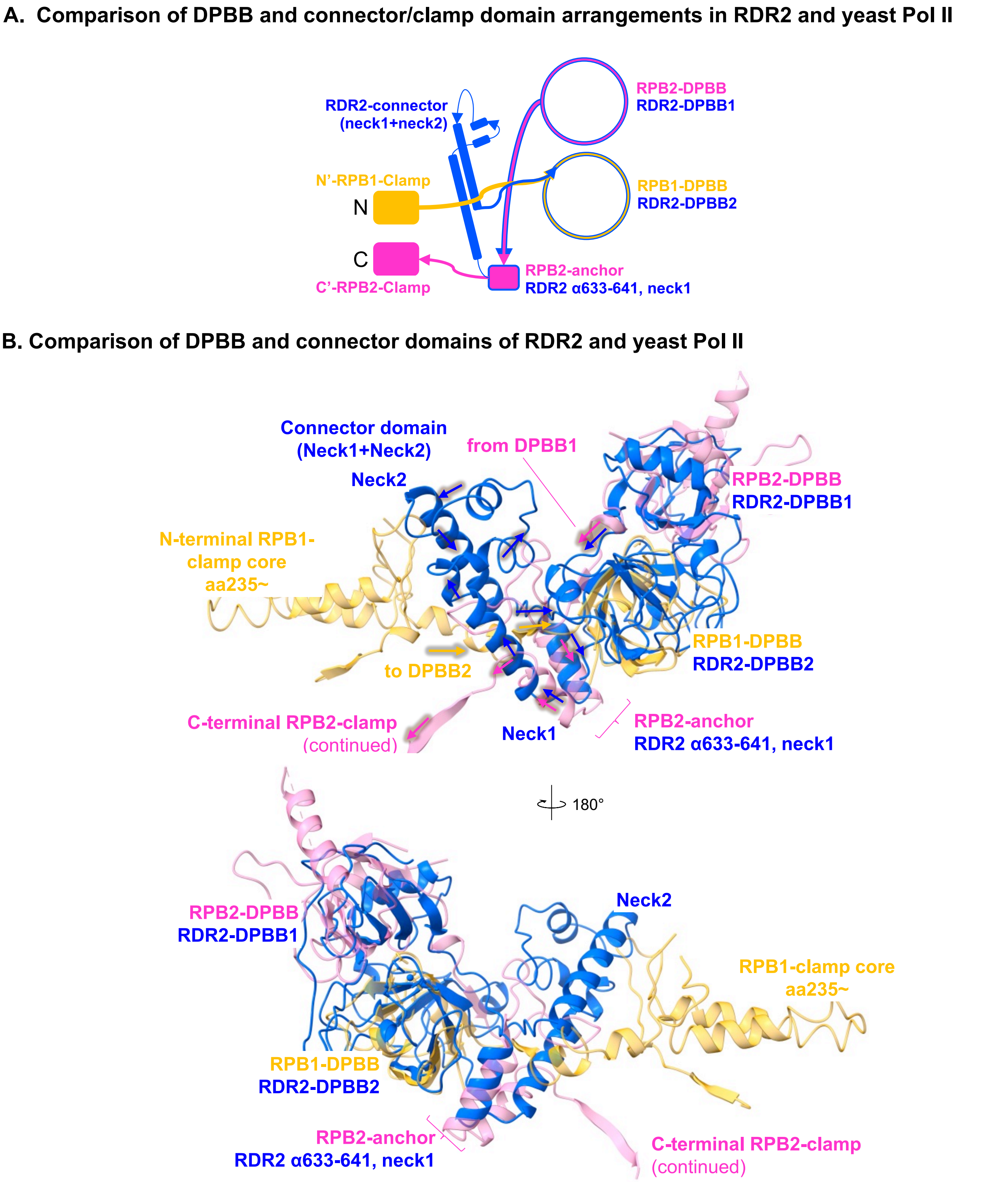

**Figure S8. The RDR2 connector domain resembles the anchor domain of yeast Pol II**
(A) Diagram of RDR2 DPBB and connector domains compared to those of yeast Pol II, including the clamp domain. (B) Structural comparison between DPBBs of RDR2 and those of yeast Pol II (2E2H). Superimposition of DPBB domains of RDR2 and those of yeast Pol II reveal similarity between a part of the connector domain of RDR2 and the anchor domain of the RPB2 subunit of yeast Pol II.

#### A. Phylogenetic tree based on amino acid alignment of 11 RDRα and 5 RDRγ proteins

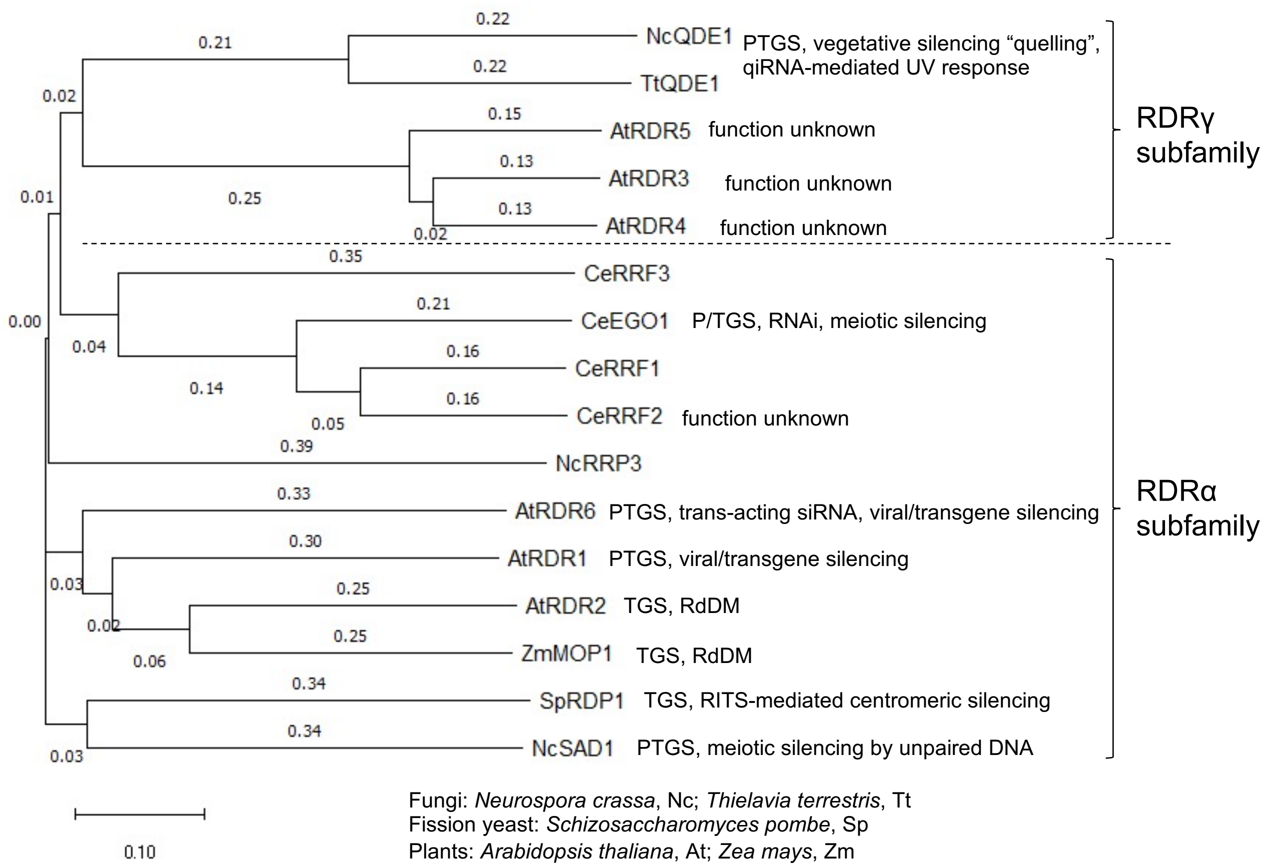

Fungi: *Neurospora crassa*, Nc; *Thielavia terrestris*, Tt
Fission yeast: *Schizosaccharomyces pombe*, Sp
Plants: *Arabidopsis thaliana*, At; *Zea mays*, Zm
Nematode: *Caenorhabditis elegans*, Ce

#### B. Percent identity among of 11 RDRα and 5 RDRγ protein amino acid sequences

| Percent Identity Matrix | | | **RDRγ** | | | | | **RDRα** | | | | | | | | | | |
|---|---|---|---|---|---|---|---|---|---|---|---|---|---|---|---|---|---|---|
| | | | fungal | | plants | | | nematodes | | | | fungal | plants | | | | fission yeast | fungal |
| | | | NcQDE1 | TtQDE1 | AtRDR5 | AtRDR3 | AtRDR4 | CeRRF3 | CeEGO1 | CeRRF1 | CeRRF2 | NcRRP3 | AtRDR6 | AtRDR1 | **AtRDR2** | ZmMOP1 | SpRDP1 | NcSAD1 |
| RDRγ | fungal | NcQDE1 | 100 | 55.86 | 16.71 | 16.95 | 17.2 | 15.57 | 15.78 | 16.02 | 16.51 | 18.23 | 18.3 | 16.12 | **16.72** | 16.58 | 16.42 | 16.41 |
| | | TtQDE1 | 55.86 | 100 | 18.53 | 17.36 | 17.89 | 17.14 | 16.56 | 15.82 | 17.4 | 17.82 | 17.95 | 17.61 | **16.2** | 17.4 | 16.52 | 16.55 |
| | plants | AtRDR5 | 16.71 | 18.53 | 100 | 71.03 | 69.97 | 18.67 | 18.99 | 17.43 | 18.51 | 18.9 | 20.52 | 20.28 | **20** | 19.6 | 20.23 | 21.22 |
| | | AtRDR3 | 16.95 | 17.36 | 71.03 | 100 | 74.32 | 18.89 | 19.88 | 17.49 | 19.05 | 19.46 | 19.18 | 19.83 | **20.45** | 19.43 | 21.13 | 20.15 |
| | | AtRDR4 | 17.2 | 17.89 | 69.97 | 74.32 | 100 | 17.98 | 18.83 | 18.12 | 19.4 | 20.34 | 21.75 | 20.86 | **21.32** | 19.31 | 21.24 | 20.29 |
| RDRα | nematodes | CeRRF3 | 15.57 | 17.14 | 18.67 | 18.89 | 17.98 | 100 | 29.82 | 29.92 | 30.22 | 19.83 | 24.19 | 25.59 | **23.29** | 24.39 | 21.61 | 21.83 |
| | | CeEGO1 | 15.78 | 16.56 | 18.99 | 19.88 | 18.83 | 29.82 | 100 | 59.66 | 56.07 | 19.28 | 24.41 | 26.71 | **22.92** | 23.4 | 24 | 21.43 |
| | | CeRRF1 | 16.02 | 15.82 | 17.43 | 17.49 | 18.12 | 29.92 | 59.66 | 100 | 68.07 | 20 | 25.45 | 27.76 | **24.29** | 23.91 | 23.11 | 22.83 |
| | | CeRRF2 | 16.51 | 17.4 | 18.51 | 19.05 | 19.4 | 30.22 | 56.07 | 68.07 | 100 | 20.41 | 25.48 | 28.2 | **23.81** | 23.73 | 22.51 | 21.88 |
| | fungal | NcRRP3 | 18.23 | 17.82 | 18.9 | 19.46 | 20.34 | 19.83 | 19.28 | 20 | 20.41 | 100 | 25.83 | 25.37 | **26.13** | 25.42 | 23.51 | 24.4 |
| | plants | AtRDR6 | 18.3 | 17.95 | 20.52 | 19.18 | 21.75 | 24.19 | 24.41 | 25.45 | 25.48 | 25.83 | 100 | 34.86 | **33.85** | 34.56 | 25.86 | 27.39 |
| | | AtRDR1 | 16.12 | 17.61 | 20.28 | 19.83 | 20.86 | 25.59 | 26.71 | 27.76 | 28.2 | 25.37 | 34.86 | 100 | **38.69** | 39.46 | 27.57 | 29.93 |
| | | **AtRDR2** | **16.72** | **16.2** | **20** | **20.45** | **21.32** | **23.29** | **22.92** | **24.29** | **23.81** | **26.13** | **33.85** | **38.69** | **100** | **49.78** | **25.63** | **26.81** |
| | | ZmMOP1 | 16.58 | 17.4 | 19.6 | 19.43 | 19.31 | 24.39 | 23.4 | 23.91 | 23.73 | 25.42 | 34.56 | 39.46 | **49.78** | 100 | 26.32 | 27.87 |
| | fission yeast | SpRDP1 | 16.42 | 16.52 | 20.23 | 21.13 | 21.24 | 21.61 | 24 | 23.11 | 22.51 | 23.51 | 25.86 | 27.57 | **25.63** | 26.32 | 100 | 32.07 |
| | fungal | NcSAD1 | 16.41 | 16.55 | 21.22 | 20.15 | 20.29 | 21.83 | 21.43 | 22.83 | 21.88 | 24.4 | 27.39 | 29.93 | **26.81** | 27.87 | 32.07 | 100 |

**Figure S9. Amino acid alignment, phylogeny and conservation among RDRα and RDRγ subfamily proteins.**
(A) Phylogenetic tree of the amino acid sequence alignment of 11 RDRα and 5 RDRγ proteins by Clustal omega. (B) Percent identity matrix of the 11 RDRα and 5 RDRγ proteins.

**Figure S10. Amino acid alignment of 11 RDRα and 5 RDRγ proteins (page 1 of 14 pages)**

Amino acid sequence alignment of 11 RDRα and 5 RDRγ, and conservation of key residues in the domains described in Figure 1 and 2. Domain annotation (colored lines above sequence) is based on the RDR2 structure in this study. *Neurospora crassa*, Nc; *Thielavia terrestris*, Tt; *Schizosaccharomyces pombe*, Sp; *Arabidopsis thaliana*, At; *Zea mays*, Zm; *Caenorhabditis elegans*, Ce.

```
NcQDE1     --------------------------------------------------MNPIT-----  5
TtQDE1     -------------------------------------------MGSAAGSAFLAT-----  12
AtRDR5     ------------------------------------------------------------  0
AtRDR3     ------------------------------------------------------------  0
AtRDR4     ------------------------------------------------------------  0
CeRRF3     -MLP-------------FDNDDSSDDATTSVRPKHPRGVPQSQSTFPRGRSNFSSGTLPN  46
CeEGO1     ------------------------------------------------------------  0
CeRRF1     ------------------------------------------------------------  0
CeRRF2     ------------------------------------------------------------  0
NcRRP3     ------------------------------------------------------------  0
AtRDR6     ------------------------------------------------------------  0
AtRDR1     ------------------------------------------------------------  0
AtRDR2     ------------------------------------------------------------  0
ZmMOP1     ------------------------------------------------------------  0
SpRDP1     MAVSLNDFI-------SVKLKRYS-RESPWERLRVPYRNKKQ------------------  34
NcSAD1     -MSGIPNWVEQAQHAILVSPDRSP-KD-QDARPRHPSRRRSSQQGH---RENTPTPRPPS  54
```

RDR2-RRM RNP2

N-terminal

```
NcQDE1     ----------------------------PRKRNSP---VEEIINRLNNDY--NLGLQCVA  32
TtQDE1     ----------------------------PSRTASA---VDKVIQQLNDDY--GLGLRI-P  38
AtRDR5     ------------------------------------------------------------  0
AtRDR3     ------------------------------------------------------------  0
AtRDR4     ------------------------------------------------------------  0
CeRRF3     RKTECTPVNTLTIGHSNKMLLTTFRMDRNSKSKSE-VDVQEQPVHSSSSAFPGNHLNNFS  105
CeEGO1     ----------------------------------------------------------MG  2
CeRRF1     -----------------------------------------------------------M  1
CeRRF2     ------------------------------------------------------------  0
NcRRP3     -------------------------------------------------------MEVYI  5
AtRDR6     -------------------------MGSEG------------------NMKKSVVTQVSI  17
AtRDR1     -----------------------------------------------------MGKTIQV  7
AtRDR2     -------------------------MVSET---------------------TTNRSTVKI  14
ZmMOP1     -----------------------MSTAAPA---------------------PGSTATVRV  16
SpRDP1     ------------------KKWASVH-NNEAQLHSANKRNDNCLIQRSSTWRLGDMITLVI  75
NcSAD1     RP-TSRPSSSSPK--PPPRAFRQFSRGKPASFASLPSL--TQPVFISREWKCARTAAIRV  109
```

RDR2-RRM RNP2 RDR2-RRM RNP1

N-terminal

```
NcQDE1     DTTLTPHRRKELAESDEDFGRHDKIYRALNF-----------------LYWRKDDS----  71
TtQDE1     DPALSPSRHNQLAEQDEQYAQWLRIFRGIKF-----------------LYYQRGDS----  77
AtRDR5     ------------------------------------------------------------  0
AtRDR3     ------------------------------------------------------------  0
AtRDR4     ------------------------------------------------------------  0
CeRRF3     YPVNRGYLRDYLLQ--SQRPST---SKPVD--CSVLKRHSLPSTHILYEKTKHRGGVNIE  158
CeEGO1     DEGYRGWIKLEIPCSLPERQ-M---GPIVK--CHVAKLEPALNEYNI--KVLTKGQVQVV  54
CeRRF1     SERSHGFIKFEFPEFDNSLQVI---EGSID--TMLLTFTSVLRKYEI--EIKSRQETQIV  54
CeRRF2     MSGQNGSIRLEFPEFDDSIDRI---EYGIE--MMLNSVPAILQHHKI--DIKSRHEMEIV  53
NcRRP3     RNLPQGLSERSLTAH---LKPFMDNLGIRHYDCNK-------------RS----------  39
AtRDR6     GGFGESTTAKQLTDY---LEDEVG---IVWR-CRLK------------TSWTPPGSYP--  56
AtRDR1     FGFPNGVSAEEVKKF---LERLTG-SGTVYA-IKVRQPK---------KG-GPR------  46
AtRDR2     SNVPQTIVADELLRF---LELHLG-EDTVFA-LEIPTTR---------DNWKPR------  54
ZmMOP1     SNIPASAIAAELLAF---FDSAVTIAGATFA-CEIVAAH---------RGWLSR------  57
SpRDP1     KDIPVTWLSNEGGKLYNLWEPLHD-YGTIEF-MKINEPL---------NG--Q-------  115
NcSAD1     KDVPP------RAKLQDLHDLFSE-YGHISY-IELDEDR---------QDVSD-------  145
```

#### Figure S10. Amino acid alignment of 11 RDRα and 5 RDRγ proteins (page 2 of 14 pages)

Amino acid sequence alignment of 11 RDRα and 5 RDRγ, and conservation of key residues in the domains described in Figure 1 and 2. Domain annotation (colored lines above sequence) is based on the RDR2 structure in this study. *Neurospora crassa*, Nc; *Thielavia terrestris*, Tt; *Schizosaccharomyces pombe*, Sp; *Arabidopsis thaliana*, At; *Zea mays*, Zm; *Caenorhabditis elegans*, Ce.

RDR2-RRM RNP1

N-terminal

```
NcQDE1    LNQAEANFFIEAKAA-------------------------SSNWVP--------------   92
TtQDE1    LEQALDSFFLEARAA-------------------------SRRWVP--------------   98
AtRDR5    ------------------------------------------------------------   0
AtRDR3    ------------------------------------------------------------   0
AtRDR4    ------------------------------------------------------------   0
CeRRF3    EQEKLVRMLWAAAEESE-----TVAKTRQFSK--KQAI--ELNFDAKLIGSMNNDCFGYC   209
CeEGO1    EEQDCEP-FYETN---------------------YEVA--TSRFSHDLIAAIQTYLKD--   88
CeRRF1    EEQDCDC-FFEIN---------------------FEVE--SEQFDHTIIDAMHDYLSD--   88
CeRRF2    EEQDCDS-FFEVN---------------------YDVA--SAKFDQTIIDAMRAYLFS--   87
NcRRP3    --------------------------NQPFGKVIFITAREALNFIAKHPRLYIMNQQAEC   73
AtRDR6    --------NFEIADTSNIPSIDEYKKVEPHAFVHFAVFESA-------GRAM--DAAGQC   99
AtRDR1    ----------------------------VYAIVQFTSERHT-------RL----IITAAA   67
AtRDR2    ----------------------------DFARVQFTTLEVK-------SRAQ--LLSSQ-   76
ZmMOP1    ----------------------------GHGFVQFDSSAAA-------THAI--DLASSG   80
SpRDP1    --------------------------TSTTAIVQFAPPPKV-------PF-W--EPNGKI   139
NcSAD1    --------------------------RGRRALIRFEPPPLS-------TD-F--LCRGIC   169
```

N-terminal

```
NcQDE1    -------KAH-------ADPDTLPW--SKEPPRAATAGQQWALQTVLLEVLNR-------   129
TtQDE1    -------KPR-------ADPGTLPS--PRSAPKAQTPDQQWNLQTILVSVLDR-------   135
AtRDR5    ------------------------------------------------------------   0
AtRDR3    ------------------------------------------------------------   0
AtRDR4    ------------------------------------------------------------   0
CeRRF3    RAHMENIKDVLKTHLKLSKVDEVNWIKVGMVPRAAYEDKSYVIDAHLVLTPNGEVEDENE   269
CeEGO1    ----------------LSTDHLMPFQRGNL------------------------------   102
CeRRF1    ----------------LN--VYVPYQRPNI------------------------------   100
CeRRF2    ----------------LN--A--NWPRPYI------------------------------   97
NcRRP3    RVSKPQPKDLKN---LVNGILYEAE--EKERRKKAWE-----------------------   105
AtRDR6    NL-ILDGQPLKV------SLGPKNP-YSLNQRR---------------------------   124
AtRDR1    ERLYYGRSYLKA------FEVE-----QDIVPK---------------------------   89
AtRDR2    SKLLFKTHNLRL------SEAY-----DDIIPR---------------------------   98
ZmMOP1    RLPPFLGSCLSV------SPAR-----ADLLPR---------------------------   102
SpRDP1    NVK---GVDLAV------QIDITAH-RSH-ISRQVFS-----------------------   165
NcSAD1    RLR-IGLVDHWL------PLETVDPKREDRKERTIKT-----------------------   199
```

N-terminal

```
NcQDE1    ----------------------------------FMPPPNNTPGR--TFGRTLSGPS---   150
TtQDE1    ----------------------------------FKAQSRAPPLA--LS--AVSGRA---   154
AtRDR5    ------------------------------------------------------------   0
AtRDR3    ------------------------------------------------------------   0
AtRDR4    ------------------------------------------------------------   0
CeRRF3    LFSEFASSFTSRITGMLHDQVFLEVPKMHTLFTKITPQHMDINISAIAIGNCPNSGLFLV   329
CeEGO1    ---------------VLHSSDF--------WSSELTCHLVDIPLAAVFFGNIQGGTF---   136
CeRRF1    ---------------VLHGSDF--------WLRTLDCHT-EIPLAAIYFGNIQGGTY---   133
CeRRF2    ---------------VLHGSDF--------WAKTLDCYI-EVPLAAMYFGNIQGYTY---   130
NcRRP3    ------------------------------ENTSNNARPARTPTLAL------DADV---   126
AtRDR6    ----------------------------------RTTVPYKLAGITLEIGTLVSRD----   146
AtRDR1    ----------------------------------PRASLHTISGLKMFFGCQVSTK----   111
AtRDR2    ----------------------------------PVDPRKRLDDIVLTVGFPESDEK---   121
ZmMOP1    ----------------------------------APDLSLRAASASLILGNRV-AER---   124
SpRDP1    ------------------------------KNSFRSDQLVKIPLSSFKLGQVYDER----   191
NcSAD1    ------------------------------QLGNTCPQTLYVFPSTLSFGLLVQPA----   225
```

**Figure S10. Amino acid alignment of 11 RDRα and 5 RDRγ proteins (page 3 of 14 pages)**

Amino acid sequence alignment of 11 RDRα and 5 RDRγ, and conservation of key residues in the domains described in Figure 1 and 2. Domain annotation (colored lines above sequence) is based on the RDR2 structure in this study. *Neurospora crassa*, Nc; *Thielavia terrestris*, Tt; *Schizosaccharomyces pombe*, Sp; *Arabidopsis thaliana*, At; *Zea mays*, Zm; *Caenorhabditis elegans*, Ce.

N-terminal

```
NcQDE1    -------------GLSR--PTSTNT-KRKDEPAN------------VTFADP--------   174
TtQDE1    -------------GARG--PPQLPAPAEESDPGS------------L-------------   174
AtRDR5    ------------------------------------------------------------   0
AtRDR3    ------------------------------------------------------------   0
AtRDR4    ------------------------------------------------------------   0
CeRRF3    RGDFISQENT---VCSVKLQSHHNADA-SRENSSFKVAGSNKYLSYARFEHDKR--LAVV   383
CeEGO1    ----INHWEV---SFWDDVRRRKSARTRNTEPTQADKIGMNQI--KVEFEFDKIDFMTVH   187
CeRRF1    ----FNHWQV---SFSRENISS--------------RDMLHKI--HAEFEFDKTDMITVQ   170
CeRRF2    ----ISHWQI---SFSGKKISA------------DANELLNKI--VAEFEFDRADMITVT   169
NcRRP3    --LDCGHWSYIDGNLTFITEW--------------------SSSLRVSAKFAKHDLVIT-   163
AtRDR6    --DFFVSWRAEGVDFLVD-----------------------PFDNTCKFCFRKSTAF---   178
AtRDR1    --KFLTLWSAQDVCVSFG-----------------------IGMRKLHFSFSW-------   139
AtRDR2    --RFCALEKWDGVRCWIL-----------------------TEKRRVEFWVWE-------   149
ZmMOP1    --ELEVAYSCDGVRAEVI-----------------------PRMRRVDLYLKH-------   152
SpRDP1    --IV----PLFGVDCGIT-VT--------------------ESNLLVYFNFKKLCVL--F   222
NcSAD1    --VF-----MKKQSVQTL-SS--------------------DSFLRLEFDFKRMRLVIKF   257
```

N-terminal

```
NcQDE1    ------------------------------PKRSLTRSATGPP------IHGAAIPLKFP   198
TtQDE1    ----------------------------------------------------------PE   176
AtRDR5    ------------------------------------------------------------   0
AtRDR3    ------------------------------------------------------------   0
AtRDR4    ------------------------------------------------------------   0
CeRRF3    YFGVR-LAEFADDGLDHAGFRLNLYYNLFV------RIVVDMSHETT-----NSIYIQMK   431
CeEGO1    FKHFENDFEVADKDAKRTKQTVTMYYQITVRRTSIRRIIVDPVVQDCNGSDRIRVHFELN   247
CeRRF1    FQCFEEKKQKF---EDSRKQKVRVNYQLTIRRDSIRRIIVDPRVEGCN----TCVHFEVN   223
CeRRF2    FQCLKRK---------------RINYQTIIRKDTIRRIIVDPQVDMNK----TRIHFELN   210
NcRRP3    ---------------------SLAESQVSIPYRYIHELVWSN---GGHVAVTLAHAPTFL   199
AtRDR6    --SFKD-AVM--------HAVINCDYKLELLVRDIQTVRQYK-T-----LHGFVLILQLA   221
AtRDR1    ---------------------YQKDYRLELSYENIWQIDLHS-PQGRS-S--KFLVIQVI   174
AtRDR2    ---------------------SGDCYKIEVRFEDIIETLSCC-VNGDA-SEIDAFLLKLK   186
ZmMOP1    ---------------------DSQSYKLEVLFEDINECFGCH-LDGTG-----AILLQLT   185
SpRDP1    DASFD---------------KQIETFRLDFDFHSIIGDVGTD---YY--DDHISLVFRFR   262
NcSAD1    SLHYQRELSG--------FRRQNEHYKLHIKFGVIKELCRTM---VGE-EHRQALVITLR   305
```

N-terminal

```
NcQDE1    DPVNTGSKRPSLESENLNQCTKRAKGKLSDNVAAAAAPPVPIASALDKVPTRRH------   252
TtQDE1    SPASTGSKRSSDGALGL--DAKRWKGQPP--ARSPSPTPTVLSNALDNVPSRRR------   226
AtRDR5    ---------------------------------------------MNQASSRIRIALSGR   15
AtRDR3    ----------------------------------------MMNNGCDEVSPRSEIALLGS   20
AtRDR4    -----------------------------------MMTTTMDYNSSDQGFSWSEIALLGS   25
CeRRF3    N------------------PPHLWEGIPKNTIFHPSKSKVL---NMETCTEWTR-VLSWP   469
CeEGO1    C------------------PVLIRRAYRTAKQESENRHS---------VPHYRRYLVI-N   279
CeRRF1    C------------------PPLIRKGYIDNDKSSFHKPF---------YERQKRFDCDWR   256
CeRRF2    C------------------PPLIRQGSVDDDKPSTQKPF---------YKRTNRYSCI--   241
NcRRP3    S------------------PPIPLLRPG-Q------------------------------   210
AtRDR6    S------------------SPRVWYRTADDDIYDTVP-----GDLLDDDDPWIRTTDFTQ   258
AtRDR1    G------------------APKIFEKED-QPINL--LFGIMDFYSDGSDEQWIRTTDFTS   213
AtRDR2    Y------------------GPKVFKRVT-VHIATKFKSDRYRFCKEDFDFMWIRTTDFSG   227
ZmMOP1    Y------------------APRIHIAISGSTVKSRFTDDRFHACKEDAKFAWVRALDFTP   227
SpRDP1    F------------------SPLIFRKSKNA-TES---RVQT---FWTASHLWRRHYDILP   297
NcSAD1    D------------------PPVAYRKRDVSKTFG---EDRL---TWSENDLWERVVDISP   341
```

**Figure S10. Amino acid alignment of 11 RDRα and 5 RDRγ proteins (page 4 of 14 pages)**

Amino acid sequence alignment of 11 RDRα and 5 RDRγ, and conservation of key residues in the domains described in Figure 1 and 2. Domain annotation (colored lines above sequence) is based on the RDR2 structure in this study. *Neurospora crassa*, Nc; *Thielavia terrestris*, Tt; *Schizosaccharomyces pombe*, Sp; *Arabidopsis thaliana*, At; *Zea mays*, Zm; *Caenorhabditis elegans*, Ce.

N-terminal

```
NcQDE1    ---------------------ANTRDPTATGH-RRADQVDSF-DTSQGTSYGSSVFSACR   289
TtQDE1    ---------------------LGFPPMDWSPQRQRLDAERSISSDTSGSSKVSSLFSRLD   265
AtRDR5    ---IETAL---ENI----YRKHNL-TPINDETRQRLSSIPENLGFEL-VRKVFSLQAGLI   63
AtRDR3    ---VETML---EKI----YEKHNHRLPISVETRRKLSSISEELALET-LRKVFNKP--YL   67
AtRDR4    ---VETML---EKV----YGKHNHHPPIKVETRRRLSSISEELALET-LRKVLNMP--NV   72
CeRRF3    GDAE-------GRGVGCTSEAFSQSSWIRLTMRK--DDDNDSVSSTQLMDIVTRL-----   515
CeEGO1    RGRS-------ANQY-PTAKAITDSPVFTIEFDQSVG-------LNEIYRLLSRL-----   319
CeRRF1    NGNV-------NHGN-PQDAAIADSPFFTIEFHKEIS-------TKEMYRVLSRL-----   296
CeRRF2    -GTK-------EYGS-PHESAISDSPFFTIELQKQESDGNSGDSNDTLYRVLSRL-----   287
NcRRP3    -------PKRMRLEACDSN--HAKVSKFCLVYHFKVSDKH--LKQNQGSYRGSDF-----   254
AtRDR6    ------------------VGAIGRCHSYRVLISPRYE-NK--------------L-----   280
AtRDR1    ------------------SSCIGQSTAFCLELPVHLNVPD--------------F-----   236
AtRDR2    ------------------SKSIGTSTCFCLEVHNGSTMLD--------------I-----   250
ZmMOP1    ------------------NSSFGECSTLVLKLSKGASVSY--------------I-----   250
SpRDP1    FNVSPTTASPIELLNCH-NAPIGRCNVLVLSFSIRDESDKDDIAFLLHNLEKFNL-----   351
NcSAD1    GLDVS--KNPVSLAENHQYIDLGRWLTYWIELDQQSTRVWDQV---QQYLLDWNL-----   391
```

N-terminal

```
NcQDE1    HNQSTTQSSFEAPPSQPREKRPVDATVF-----------EAG-HL---------------   322
TtQDE1    SQRSTQTTSD----GEEREQKH-GPTVIVS------SLSQQG-LS-----------AATN   302
AtRDR5    YNLDSFIVSKV---NQAVSFTG--YPQPAN------SLSPSGRHVSRVLQ----EEMSVD   108
AtRDR3    KTLDGLIMYFV---KGTVTVDG--SPRLSPGESPVQSPRTPAKKSCRASQ-----DVSLD   117
AtRDR4    KTLDGIIIYFL---NDAVTVDG--SPRLWSGESPVQFPRTPGKKSCRASQ----AEVSLD   123
CeRRF3    SARSKAKVMFGSIFSIRRKLAPSPA-----------------------------------   540
CeEGO1    RIRTGVSIEFADIPSIDCLIWRENPYNRWT------FLNNQH------LSPTH-FSAPI-   365
CeRRF1    RSRTKVLIEFANLPSIDVPMGSHYPYNRWN------LK----------KSPTD-SNAPI-   338
CeRRF2    RSRTGVQIEFANFPKVDVPIGKVYPYLRYP---------------------TS-SKSYA-   324
NcRRP3    R--TAINGL-----NE--------QEMFWV------T---HYTFAIQTTSPQDAYAVAVD   290
AtRDR6    R--TALDYFRMRRVQE--------ERVRWP------PRIRNEP-----C-----FGEPV-   313
AtRDR1    R--ENFANYAEHRASS--------------------FLIESGS-----S-----YSSNA-   263
AtRDR2    F--SGLPYYREDT-LS--------------------LTYVDGK-----T-----FASAA-   276
ZmMOP1    L--ESLPFSGELG--E--------------------LAIASMD-----V-----FGSSS-   275
SpRDP1    K--SQLD-------------------KVVF------HLVPDYKHRCSLI-----N-----   374
NcSAD1    R--TKLTVFPEPLPNQ--------KPKVWD------FLDDRYGHDIQQV-----SSRSW-   429
```

N-terminal

```
NcQDE1    IESPSKGRTTK---SHIDNQPLSSSSQGETSFSTYYESFPSSGGEGAIPE------PSRS   373
TtQDE1    NTVPPPTRTARPVTGGASRAPPRSSAA-----GTLYSDLSHVAWDDSVAE---------I   348
AtRDR5    SDAPS-PKSLKSEDKGGSLHIPQLVALGELEFKKVFLLLSYIPGQHV-GQVITADEI---   163
AtRDR3    LETPS-PKFMKREENGGSKYIPPLLALGELEFKKAFLLLSYIGGESLVEEVISGDQI---   173
AtRDR4    REDPS-PKFLRGDENGESKHISLLLALGELEFKKAFLLLTYLGGKSL-GEVISGDEI---   178
CeRRF3    -------------------------FHSLGSFRANYALQALITRGSVFTDQLFDATDENI   575
CeEGO1    YRDFITTAFPKKHEVCGSREV---DTNRERKFAITYLLECLISRGAVVKDQILLDEG---   419
CeRRF1    FREFLKEIFPPKYEIVDDKLI---DVNEERKFSITYLIECLLSRGAIVKDQLLLNEQ---   392
CeRRF2    FECFIYNCFPSKMKIIDAQSI---NENDGRQFAITYLIECLLSRGAIVKDQVLTDEI---   378
NcRRP3    RLRAALLEY---------------EGRDTLPYSLLFNLQALVYMSYLHPTTVLK--LA--   331
AtRDR6    SDHFFC--I---------------HHKEGISFEIMFLVNSVLHRGVFNQF-QLTER----   351
AtRDR1    NTLVPVVDP---------------PPGFSLPFEILFKLNTLVQNACLSGP-ALDLD----   303
AtRDR2    QIVPLLNAA---------------ILGLEFPYEILFQLNALVHAQKISLFAASDME----   317
ZmMOP1    NVVPLVD-C---------------PNGFSVPYEVLFRLNSLVHMGKLVAR-HVNAD----   314
SpRDP1    --------------------------DKEIEEEIAYLLQACLSKNLLSEI-DLP--II--   403
NcSAD1    SNDFSLLAA---------------PPRISLPFDVRYQLEVCISQGIINEH-NIDRPFL--   471
                                                    :
```

**Figure S10. Amino acid alignment of 11 RDRα and 5 RDRγ proteins (page 5 of 14 pages)**

Amino acid sequence alignment of 11 RDRα and 5 RDRγ, and conservation of key residues in the domains described in Figure 1 and 2. Domain annotation (colored lines above sequence) is based on the RDR2 structure in this study. *Neurospora crassa*, Nc; *Thielavia terrestris*, Tt; *Schizosaccharomyces pombe*, Sp; *Arabidopsis thaliana*, At; *Zea mays*, Zm; *Caenorhabditis elegans*, Ce.

```
NcQDE1    NGLARSEESARSQVQVHAPVVAARLRNIWPKFPKWLHEAPLAVAWEVTRLFMHCKVDLED   433
TtQDE1    PGSGR----NEADALPIHPEVYARLQNIWPKFPRWLHAAPLALAWELTRICLHCKVDLED   404
AtRDR5    --------------------------RLWKDL----------------------------   169
AtRDR3    --------------------------RKWKDL----------------------------   179
AtRDR4    --------------------------RQWKDL----------------------------   184
CeRRF3    PSSDNDNDED--DDDDVDDTKKPMELVHEPLFLKLVRRGM----KECSQATEETLEQLLN   629
CeEGO1    ---------------------------IWHRFLEVILHYY----TKDDKLCEAGLEDLVH   448
CeRRF1    ---------------------------HWKNFLEIIIWYY----RNDNQLCEAALEDLVH   421
CeRRF2    ---------------------------CWGQFLGLITHYY----LENDKLCEAALEDLIY   407
NcRRP3    --------------------------KRLADMFETARRSG----QRQDPISVDAFKDLFK   361
AtRDR6    -------------------------------FFDLLRNQP----KDVNI---ASLKHLCT   373
AtRDR1    -------------------------------FYRLLNQKK----YDRALI-DHCLEKLFH   327
AtRDR2    -------------------------------LIKILRGMS----LETA---LVILKKLHQ   339
ZmMOP1    -------------------------------LFKVLEDLS----IDTL---RRIFEKMSK   336
SpRDP1    --------------------------L------ANLKKLS----KERAK---KFLRLILT   424
NcSAD1    --------------------------EKLMEFCDNDNSFG----KDRAR---LILEYVAD   498
```

QDE1ΔN homodimerization interface residues

slab

```
NcQDE1    ESLG--LKYD-PSWSTARDVTDIWKTLYRLDAFRGKPFPEKPPNDVFVTAMTGNFESKGS   490
TtQDE1    PT----LRYD-PSWATS-DMAALWRSLTQLDVFRGKSFPERPSAEAFAAALTGNFESRGN   458
AtRDR5    ----PMV----EYEAA------VWDRLGRHYCPQKDR---------------RMLQWDSG   200
AtRDR3    ----PMV----SYEAA------VWNRLGQRYCSPKER--------------RRPLEGDSG   211
AtRDR4    ----PMV----AYERA------VWFKLGQN-----EE--------------RMQLESDSG   211
CeRRF3    AFDERRQIDV------------V-TAF-TTMYQSRKIQYERLLKGESLQDVGLAKPLPKN   675
CeEGO1    MIDGRKRIGS------------LIKCF-DRICQTRQR--NSLVNGLT------TEEMREG   487
CeRRF1    LIDGRKRIGS------------ILKCL-DKICQKREV--MKLVNGLT------EKESIEG   460
CeRRF2    LIDGRKRIGS------------IWKCF-HKICQKRLV--MQLTNGMS------EQEIEEG   446
NcRRP3    TIDWPSPSLSSAEELAQFEVEGIIEHL-KKTEKRMR----------EGYTLRLNEEIPPG   410
AtRDR6    ---YKRPVFD-AYKR-------L-KLV-QEWIQKNP----------KL---LGSHEQSED   407
AtRDR1    ---LGECCYE-PAHW-------LRDEY-KKWISKGK----------LPL-SPT-ISLDDG   363
AtRDR2    ---QSSICYD-PVFF-------VKTQM-QSVVKKMK----------HSPASAYKRLTEQN   377
ZmMOP1    ---LKSTCYE-PLQF-------IRHEA-HSMNMRKK--------------ALSNKRESGK   370
SpRDP1    S---KTALIN-PSEL-------DFTK--S------------------FVFYD---LSSAS   450
NcSAD1    E-YAGKRIFD-PMEL-------FKDHA-A------------------LAYFPTSFMIPNH   530
```

*rdr6(sgs2-19)*, E429K

slab

```
NcQDE1    AVVLSAVLDYNPDNSPTAPLYLVKLKPLMFEQGCRLTRRFGPDR--------FFEILIPS   542
TtQDE1    TVVLSASLEFNPS--KTGPLFLLDMKPLRFDEGCRLTRRFGPDR--------FLEVLVPS   508
AtRDR5    KTHY-YQCNVA----PNG---SYTFKV-LSALQGPLLEHTGTHLHKVLGDDNVLTVKFAD   251
AtRDR3    MTHY-YQCHVA----TDG---SYKFK-------GHLLENTGTHLHKVLGDDNVLTVKFDK   256
AtRDR4    KTHY-YQCHVA----PDG---SYRLK-------GYFLENTGTHLHKVLGDDNVLTVRFDQ   256
CeRRF3    CVSV-AKVIVT----PSR---ILLMAP-EVMMVNRVVRRFGPDY--------ALRCVFRD   718
CeEGO1    YQRV-RKIIFT----PTR---VIYVAP-ETLMGNRVLRRYDHDG------TRVLRITFRD   532
CeRRF1    YQRV-RKVIFT----PTR---VIYIAP-ETIMGNRVLRKFDKDG------TRVLRVTFRD   505
CeRRF2    YQRV-RKVIFT----PTR---VIYTPP-EMIMGNRVLRNFDRDG------THVLRVTFRD   491
NcRRP3    LTKI-YRALVT----PTR---IELHGP-ELEAKNRILRKFPEHQ------DHFLRVQFAE   455
AtRDR6    ISEI-RRLVIT----PTR---AYCLPP-EVELSNRVLRRYKAVA------ERFLRVTFMD   452
AtRDR1    LVYM-YRVQVT----PAR---VYFSGP-EVNVSNRVLRHYSKYI------NNFLRVSFVD   408
AtRDR2    IMSC-QRAYVT----PSK---IYLLGP-ELETANYVVKNFAEHV------SDFMRVTFVE   422
ZmMOP1    LMRC-YRIHIT----PSK---IYCLGP-EEEVSNYVVKYHSEYA------SDFARVTFVD   415
SpRDP1    SIHI-KKLYVT----PTT---LRIVED-SLEAGNRVIRNFKDFA------NRFMRVQITD   495
NcSAD1    CAWV-RRVTIT----PTR---IYFSTP-CVEPTNRVIRQWKHAQ------DYFIRIQFTD   575
                                     . .                        . .
```

#### Figure S10. Amino acid alignment of 11 RDRα and 5 RDRγ proteins (page 6 of 14 pages)

Amino acid sequence alignment of 11 RDRα and 5 RDRγ, and conservation of key residues in the domains described in Figure 1 and 2. Domain annotation (colored lines above sequence) is based on the RDR2 structure in this study. *Neurospora crassa*, Nc; *Thielavia terrestris*, Tt; *Schizosaccharomyces pombe*, Sp; *Arabidopsis thaliana*, At; *Zea mays*, Zm; *Caenorhabditis elegans*, Ce.

*rdr6(sgs2-3)*, E453K

slab

```
NcQDE1     PTSTSP--------SVPPV-------VSKQPGAVEEVIQWLTMGQHSLVGRQWRAFFAKD    587
TtQDE1     PTALNA--------P---S--------ILKDGGAAQVIRWLTEKPHSLVGRQWQAFYTKD    549
AtRDR5     VQK----------------------SSSTYSIDHYFTYKGIAKNGIMIGLRRYQFFVFKD    289
AtRDR3     VLG-----------------------VETYCNDLYSTYKGIAKNGIMVGLRRYRFFVFKD    293
AtRDR4     LPK-----------------------ESTYCDNPYSKYKEIAKNGIMVGLRRYQFFVFKD    293
CeRRF3     DNLGRL-AIRDFSINNI----------DHMSNIVTEGIYLTLKNGIQVADRVYSFLGWSN    767
CeEGO1     DDNQKM-RTN------------------KTSTMLEKTVNQYLKNGITVAGRNFGYLGSSN    573
CeRRF1     DNNKKM-RSN------------------VTGKLLDRTANKYLEHGVRIANREYGFLGCSN    546
CeRRF2     DNNRKM-RAN------------------ATGELLDICVKKYLEHGIVVANRDFGFLGCSS    532
NcRRP3     EDGQDLFF-----------------NSAVSMDAIYQRFKDVLTNGISVGGRVYRFLGFSH    498
AtRDR6     ESMQTI-NSNVLSYFVAPIVKDLTSSSFSQKTYVFKRVKSILTDGFKLCGRKYSFLAFSA    511
AtRDR1     EDLEKV-RSMDLSPRSS----------TQRRTKLYDRIYSVLRDGIVIGDKKFEFLAFSS    457
AtRDR2     EDWSKL-PANALSVNSKE-----GYFVKPSRTNIYNRVLSILGEGITVGPKRFEFLAFSA    476
ZmMOP1     EDWSKL-SPNALSARTEQ-----GFFSKPLKTGLYHRILSILKEGFCIGPKKYEFLAFSA    469
SpRDP1     EYYKQKIRGGSD---------------GFRNEKLYSRIQQLLTYGIKVGNQIYEFLAFGN    540
NcSAD1     EVLEGRIKSGEA---------------E---LPLFLRVYRVLEKGVAMGPWHWKFLAFGN    617
                                                     :    :  :
```

“Fork loop 3”, RDR2 aa476-480

slab

```
NcQDE1     AGYRKPLREFQLRAEDPKPIIKERVHFFAETGITFRPDVFKTRSVVPAEEPVEQRTEFKV    647
TtQDE1     AGYRKPARELRLGP-DAKATFKERVHFFAERGHDFRPAPLRTRAQLPPESPSRRRTEIRV    608
AtRDR5     GGKEEKKKDLS--------TKKVKCYFIRTDSTAF----YD--MQNP-----YILTGKSI    330
AtRDR3     GGKEEKKKDVS--------TKGVKCYFIRTDSTAS----ID--MQNP-----YIFAGKSM    334
AtRDR4     GGKAEKKKRNS--------TKQVKCYFIRTGSTAS----SD--MENP-----YILSGMSI    334
CeRRF3     SQMRD-----------------QGCYLYAPRVNAL-----------------TGEVTGTV    793
CeEGO1     SQMRD-----------------NGAYFMEKYSSSQCREYERIYQIKP-----PITFNPKI    611
CeRRF1     SQMRD-----------------NGAYFMMRFTDKQLDRFYKCNPTAS-----NINFKPKI    584
CeRRF2     SQMRD-----------------NGAYFMVKNTDNRHKN-----ACKM-----NSKFKPNI    565
NcRRP3     SSLRA-----------------HSLWLAAPFIYDG--------------------KLQLA    521
AtRDR6     NQLRD-----------------RSAWFFAE----D--------------------GKTRV    530
AtRDR1     SQLRE-----------------NSAWMFAP----I--------------------DRITA    476
AtRDR2     SQLRG-----------------NSVWMFAS----N--------------------EKVKA    495
ZmMOP1     SQLRG-----------------NSVWMFAS----N--------------------SSLTA    488
SpRDP1     SQLRE-----------------HGAYFFAS----G--------------------SDLNA    559
NcSAD1     SQIRE-----------------AGAFMFCE----Q--------------------SNLTG    636
                                 :
```

“Link domain”, RDR2 aa511-522

**RDR2 S525, 1st iNTP γ-P**

DPBB1

```
NcQDE1     SQMLDWLLQLDNNTWQPHLKLFSRIQLGLSKTYA-IMTLEPHQIR--------HHKT--D    696
TtQDE1     SEMLDWLLQLEQNEYQPHLKLFSRIQLGLSKTFP-TVTFEPNQIR--------HRTD--D    657
AtRDR5     YEARMHFMHVHRAP--TLANYMARFSLILSKTKTLEVDMTGITFD-QIDDIHCHDQDGKD    387
AtRDR3     HEARMHFMHVNTLS--SLPNYMSRFSLILSKTKTLEVDMTGITFE-QIDDIHCHDQDDKD    391
AtRDR4     HEARMHFMHVHTLP--SPANYMARFSLILSKTKKLEVDMTEITVM-QIDDIHCHDQSNND    391
CeRRF3     EDIRVWMGDFRDAI--SVPKMMSRMGQCFTQAQP-TVRLERHHWI-V-----EPDIE--G    842
CeEGO1     QAARKNLGRFETID--NIPKMMARLGQCFTQSRLSGVNLERCTYM-T-----TYDLT--G    661
CeRRF1     DEVRFQLGRFSEIE--NVPKLMARLGQCFTQSRLTGVGLGRDDYC-S-----TYDLT--G    634
CeRRF2     DSVRNQLGNFLQIE--NIPKLMARLGQCFTQSRLTGVSLGPDNYC-L-----THDLS--G    615
NcRRP3     SNIIEDLGDFRNIM--SPARRAARIGQAFSETPY-SVSLYDHGID-VI---RQRDVK---    571
AtRDR6     SDIKTWMGKFKD-K--NVAKCAARMGLCFSSTYA-TVDVMPHEVDTEV-----PDIE---    578
AtRDR1     AHIRAWMGDFDHIR--NVAKYAARLGQSFSSSRE-TLNVRSDEIE-VI-----PDVE---    524
AtRDR2     EDIREWMGCFRKIR--SISKCAARMGQLFSASRQ-TLIVRAQDVE-QI-----PDIE---    543
ZmMOP1     ENIRRWMGHFEDIR--SVSKCAARMGQLFSSSRQ-TFEVSSYDVE-VI-----PDIE---    536
SpRDP1     KQIREWMGDFSEIN--SVSKYAARLGQCFSTTKE-INRFCV-DIS-LQ-----DDIV---    606
NcSAD1     DMMRAWMGRFSHIK--VIAKYAARLGQCFSTTRL-VPGIPAPRIV-TI-----PDVE---    684
                  :   .          .   :*:   :: :        .
```

Amino acid sequence alignment of 11 RDRα and 5 RDRγ, and conservation of key residues in the domains described in Figure 1 and 2. Domain annotation (colored lines above sequence) is based on the RDR2 structure in this study. *Neurospora crassa*, Nc; *Thielavia terrestris*, Tt; *Schizosaccharomyces pombe*, Sp; *Arabidopsis thaliana*, At; *Zea mays*, Zm; *Caenorhabditis elegans*, Ce.

QDE1ΔN homodimerization interface residues

Metal B — DPBB1

```
NcQDE1    LLSPSGTGEVMNDGVGRMSRSVAKRI-RDVLGL-----------------------GDVPS   733
TtQDE1    ILSPA--GKIMNDGIGRMSRSVARKI-RDALGL-----------------------SDIPS   692
AtRDR5    VLDKNKKPCIHSDGTGYISEDLARMCPLNIFKGKCLRSESIQ---------EACYQDPPL   438
AtRDR3    VLDKNGKPCIHSDGTGYISEDLARMCPVNIFKGKSMRSNNIQSKNLNFEGQGPCGQEPPL   451
AtRDR4    VLDKNGKPRIHSDGTGYISEDLARMCPLNIFKGKSMRSNNIQ---------GTCVQEPPL   442
CeRRF3    GVE---NKYCFSDGCGRISIKLATHI-SKILQL-----------------------KEVPA   876
CeEGO1    GKNLKGDEYTFSDGVGMMSYRFAQMV-SEVMDF---------------------G-KGVPS   699
CeRRF1    GRATNGSEYTFSDGVGMMSYQFAQEV-SQAMQF---------------------G-KAVPS   672
CeRRF2    GRSSNGSEYTFSDGVGMMSYEFAQEV-SRLMSF---------------------G-RSVPS   653
NcRRP3    -----RNERVFSDGVGIISQGALEVIHREIP---------------------ESKGYPN   604
AtRDR6    -----RNGYVFSDGIGTITPDLADEV-MEKLKL---------------------DVHYSPC   612
AtRDR1    -IISLGTRYVFSDGIGKISAEFARKV-ARKCGL--------------------T-EFSPS   561
AtRDR2    -VTTDGADYCFSDGIGKISLAFAKQV-AQKCGL-----------------------SHVPS   579
ZmMOP1    -VTTDGTKYIFSDGIGKISTRFARQV-AKLIGL---------------------DPAHPPS   574
SpRDP1    -----RNNHCFTDGVGMASLSVIRRLSLEVKNH-----------------------DMFPS   639
NcSAD1    -----KDGFCFTDGVGKISPLLAKI-VAHDWSI-----------------------DPPPS   716
                     .** *   :                                   *
```

RDR2 Q582, 1st iNTP α-P — RDR2 K589, 1st iNTP α-P — DPBB1

```
NcQDE1    AVQGRFG--S--AKGMWVIDVDDTGDEDWI------------------ETYPSQRKWECD   771
TtQDE1    AIQGRMG--S--AKGMWLMDVADAGDDDWI------------------ETYPSQRKWKCD   730
AtRDR5    LIQFRMFYDGYAVKGTFLLNKKL---------------------CPRTVQVRPSMIKVSKD   478
AtRDR3    LIQFRIFYNGYAVKGTFLTNKKL---------------------PPRTVQVRPSMIKVYED   491
AtRDR4    LIQIRMFNDGSAVKGIFLLNKNL---------------------PPQTVQVRPSMIKVYKD   482
CeRRF3    CFQVRFK--G--FKGILVIDPTIDDIIN-----------------MPKVIFRKSQQKFGEG   916
CeEGO1    CFQFRFR--G--MKGVISIEPLLDNLRQWSISYNISKPSDDSSWSLNCMFRPSQIKFIS-   754
CeRRF1    CFQIRFR--G--NKGVIAIEPFLDEIRKWALVNGVTS-----MKMAKCLFRPSQIKFQA-   722
CeRRF2    CFQFRYR--G--MKGVLAIDPLLDKERNWQEKHGISL-----SKNIKCVFRPSQIKFEG-   703
NcRRP3    CLQVRWA--G--AKGMLALDARL---------------------TGRQICIRDSMEKFRSR   640
AtRDR6    AYQIRYA--G--FKGVVARWPSKS--------------------DGIRLALRDSMKKFFSK   649
AtRDR1    AFQIRYG--G--YKGVVAVDPNSS---------------------KKLSLRKSMSKFESE   596
AtRDR2    AFQIRYG--G--YKGVIAVDRSSF---------------------RKLSLRDSMLKFDSN   614
ZmMOP1    AFQIRYG--G--YKGVITIDPTSF---------------------FNLSLRPSMKKFESK   609
SpRDP1    AFQFRMG--G--YKGVLSLAPPTKLEY----------------HQGNLVFPRRSQDKFKSF   680
NcSAD1    AYQFRMG--G--CKGVLVTWPDV---------------------KGMEVHIRKSQEKFVAE   752
            *  *     .   **                                 *   *
```

DPBB1 — RDR2 R622, 1st iNTP γ-P — RDR2 aa633-641, "anchor" — Connector

```
NcQDE1    FVDK---HQRTLEVRSVASELKSAGLNLQLLPVLEDRA-RDKVKMRQAIGDRLINDLQRQ   827
TtQDE1    DADA---LHRTLEIRSVSTELKPAALNLQFLPVLEDRA-KDKARMRRAIAARLMNDLKKQ   786
AtRDR5    PSLSNFSTFNALEVVTTSNPPKRTKLSKNLVALLSYGG----------IPNEFFLDILLN   528
AtRDR3    RTLSNLSTFNSLEVVTTSNPPRKARLSRNLVALLSYGG----------VPNDFFLNILRN   541
AtRDR4    KNLSNFSTFNSLEVVTTSNPPKRAKLSKNLVALLSYGG----------VPNDFFLDILLN   532
CeRRF3    GGEL---QDEYLEVVKY-AMPSPVCLNRPFITILDQVSEKQSASSHRRITNRVHYYLERE   972
CeEGO1    -KRH---PRDQVEIVKY-SSPVPVALNKPFINILDQVSEMQSLECHRRVTNRIEELLDRQ   809
CeRRF1    -KAI---SGDQIEMVKF-SSAVLVALNKPFINILDQVSEMQSLDCHKRITSRIEELMDRQ   777
CeRRF2    -KQI---LGDQVEMVKY-SSPVLVALNKPLINILDQVSEMQSLECHRRITSRIEELMDLQ   758
NcRRP3    -------DEEHLEICDMASKPIPLMLNRQMIKILEDMR----------APAQWFLELQEK   683
AtRDR6    -------HT-ILEICSW-TRFQPGFLNRQIITLLSVLG----------VPDEIFWDMQES   690
AtRDR1    -------NT-KLDVLAW-SKYQPCYMNRQLITLLSTLG----------VTDSVFEKKQRE   637
AtRDR2    -------NR-MLNVTRW-TESMPCFLNREIICLLSTLG----------IEDAMFEAMQAV   655
ZmMOP1    -------ST-MLNITNW-SKSQPCYVNREIISLLSTLG----------IKDEVFESMQQD   650
SpRDP1    -------HS-TLEVIKI-SRFSNAHLNMQLITLLEGLG----------VEKTVFLELTRS   721
NcSAD1    -------FN-GLEVVRC-SQFSTATLNRQIIAVLSSLG----------VPDQVFVDMMEQ   793
                 :::                :.   ::  :*.
```

### Figure S10. Amino acid alignment of 11 RDRα and 5 RDRγ proteins (page 8 of 14 pages)

Amino acid sequence alignment of 11 RDRα and 5 RDRγ, and conservation of key residues in the domains described in Figure 1 and 2. Domain annotation (colored lines above sequence) is based on the RDR2 structure in this study. *Neurospora crassa*, Nc; *Thielavia terrestris*, Tt; *Schizosaccharomyces pombe*, Sp; *Arabidopsis thaliana*, At; *Zea mays*, Zm; *Caenorhabditis elegans*, Ce.

QDE1ΔN homodimerization interface residues

Connector

```
NcQDE1    FSEQKHALNRPVEFRQWVYESYSSRATRVSHGRVPFLAGLPDSQEETLNFLMNSGFDPKK 887
TtQDE1    FDSQKAAVERPLQFRQWVNECTNSRSERVRHGQVPFLGGLPENKGEVLSFLLNSGFDR-R 845
AtRDR5    TLEESKSIFYNKRA---------ALNA-----ALNYGE----MDDQNAAQMILVGIP-LD 569
AtRDR3    TLEESKTIFYSERA---------AFKA-----AINYGD------DQYTADMILVGIP-LD 580
AtRDR4    TLEKKKTIFFKVRA---------AGKA-----ALHYGN----MDDKNALQMIMAGIP-LD 573
CeRRF3    LCSLSNMLINENQA----------AEE-----LVNRTN----LAIDWNAASKRAGFELSV 1013
CeEGO1    MLSFAQQMVDETFC----------RNR-----LK-ELP----RRVDIDYLRTTWGFTLSS 849
CeRRF1    ILSFAKQMNEETFC----------RNK-----LK-EFP----RRIDIDNLRTMWGFTLSS 817
CeRRF2    TLSFAKQMTDEACF----------RNK-----LK-EFP----RRIDIDYLRTIWGFTLSN 798
NcRRP3    ELQRLRAITDNVQN---------VATF-----LKLQCVGDSVHLSQFLKDLDKMNIDYRR 729
AtRDR6    MLYKLNRILDDTDV---------AFEV-----LTASCA----EQGNTAAIMLSAGFKPKT 732
AtRDR1    VVDRLDAILTHPLE---------AHEA-----LGLMAPG---ENTNILKALILCGYKPDA 680
AtRDR2    HLSMLGNMLEDRDA---------ALNV-----LQKLSGE---NSKNLLVKMLLQGYAPSS 698
ZmMOP1    DMHESDGMLTNKEA---------ALSV-----LGKIGGG---DTK-TAADMLLQGYEPSS 692
SpRDP1    QLSKMNESINSKQK---------SILM-----LRDNVDE--YHSTLIIADFIQAGFLERD 765
NcSAD1    QLSDFNAAMEDKQK---------ATAI-----LKTFIDE--NHMTPIIAEMLAYGFMGSQ 837
                                                                .
```

Connector

DPBB2

```
NcQDE1    QKYLQDIAWDLQKRKCDTLKSKLN--IRVGRSAYIYMIADFWGVLEE------------- 932
TtQDE1    QKYIQDLAFDLQKQRCEVLRTKLN--IHVGRSAYMFMVVDFWGVLEE------------- 890
AtRDR5    EPHLKNYLSILLKTEKNDLK-AGK--LPVTESYYLMGTVDPTGALKE------------- 613
AtRDR3    EPYLKDRLSYLLKTERNALK-AGR--FPIDESYYIMGTVDPTGELKE------------- 624
AtRDR4    EPYLKHYLSKLLKLEKDDLK-AGK--LPIDESYYLMGTVDPTGELKE------------- 617
CeRRF3    DPLIRDMLFSIYRYNIIHHISKAKIFLPPSLGRSMYGVVDETGLLQY------------- 1060
CeEGO1    EPFFRSLIKASIKFSITRQLRKEQIPIPCDLGRSMLGVVDETGRLQY------------- 896
CeRRF1    EPFFRSLIKASIKFSITKQLCKEQIQIPSELGRSMLGVVDETGRLQY------------- 864
CeRRF2    EPFFRSLVKASIKFAITKQLFKEQIQIPSELGRSMLGVVDETGILQY------------- 845
NcRRP3    DQFLRGIVEAVVLRELRLLKHKAR--IPVPYGVTLFGVMDETGLLRE------------- 774
AtRDR6    EPHLRGMLSSVRIAQLWGLREKSR--IFVTSGRWLMGCLDEAGILEH------------- 777
AtRDR1    EPFLSMMLQNFRASKLLELRTKTR--IFISGGRSMMGCLDETRTLEY------------- 725
AtRDR2    EPYLSMMLRVHHESQLSELKSRCR--ILVPKGRILIGCMDEMGILEY------------- 743
ZmMOP1    EPYLLMILKAHRANRLTDIRTRCK--IHVQKGRVLIGCLDETCKLEY------------- 737
SpRDP1    DAFTENLLNLYYEWVLRLIKEKQK--VSVPKGAYLLGVADETGTLKGHYDD-------AV 816
NcSAD1    EPFVRTLLQLWRSWSIKTLKEKAR--LNVEKSAFVLGCVDETGTLKGHMKVIEDWKDVSS 895
          :                      .  .    .  :    *    *.
```

NcQDE1ΔN homodimerization interface residues

RDR2 K783, 1st iNTP 2'-/3'-OH

RDR2 P765, “Pro-gate”

*rdr6(sgs2-1)*, G825E

*rdr6(sgs2-5)*, D826N

DPBB2

```
NcQDE1    ---NEVHVGFSSKFRDEE--E----------SFTLLSDCDVLVARSPAHFPSDIQRVRAV 977
TtQDE1    ---NEVHVGFSSKFRDDVDDT----------TYMLLTDCDVLVARSPAHFPSDIQKVRAV 937
AtRDR5    ---DEVCVI----------------------LESGQISGEVLVYRNPGLHFGDIHILKAT 648
AtRDR3    ---NEICVI----------------------LHSGQISGDVLVYRNPGLHFGDIHVLKAT 659
AtRDR4    ---DEVSGL----------------------A----KSQDVLVYRNPGLHFGDIHILKAT 648
CeRRF3    ---GQVFIQYSPSIRQT---------SN----RPILKTGKVLITKNPCHVPGDVRVFDAV 1104
CeEGO1    ---GQIFVQYTKNLALKLPPK----NAARQ-----VLTGTVLLTKNPCIVAGDVRIFEAV 944
CeRRF1    ---GQIFVQYTKNYKKKLPPR----DSNNKVHGSEIVTGTVLLTKNPCIVPGDVRIFEAV 917
CeRRF2    ---GQVFVQYTKNHRNILPPR----DSNRKVLGSEIVTGTVLLTKNPGIVPGDVRIFEAV 898
NcRRP3    ---GEVYVTFETVDG---------RF------KDPPTAGPVVVTRSPALHPGDIQIAHNA 816
AtRDR6    ---GQCFIQVSKPSIENCFSKHGSRFKE-TKTDLEVVKGYVAIAKNPCLHPGDVRILEAV 833
AtRDR1    ---GQVVVQYSDPM---------------RPGRRFIITGPVVVAKNPCLHPGDVRVLQAV 767
AtRDR2    ---GQVYVRVTLTKAELKSRD-QSYFRK-IDEETSVVIGKVVVTKNPCLHPGDIRVLDAI 798
ZmMOP1    ---GQVYIRITKNHKEQKYSE-QPFFCN-DDGKTAVIVGKVAITKNPCLHPGDVRVLEAV 792
SpRDP1    LSVPEIFIQITDTSTSF------GSYST-GKLKTRVIVGLCIVARNPSLHPGDVRVCKAV 869
NcSAD1    EKLPQIFLQIPDDVN----------------GGYRVITGTCVVGRNPSLHPGDIRVVEAV 939
              :                                     : :.*    .*::
```

**Figure S10. Amino acid alignment of 11 RDRα and 5 RDRγ proteins (page 9 of 14 pages)**

Amino acid sequence alignment of 11 RDRα and 5 RDRγ, and conservation of key residues in the domains described in Figure 1 and 2. Domain annotation (colored lines above sequence) is based on the RDR2 structure in this study. *Neurospora crassa*, Nc; *Thielavia terrestris*, Tt; *Schizosaccharomyces pombe*, Sp; *Arabidopsis thaliana*, At; *Zea mays*, Zm; *Caenorhabditis elegans*, Ce.

QDE1ΔN homodimerization interface residues — Metal A — DPBB2

```
NcQDE1      FKPE-LH----SLKDVIIFSTKGDVPLAKKLSGGDYDGDMAWVCWDPEIVDGFV--NAEM   1030
TtQDE1      FKPQ-LH----ALKDVIVFPAKGDIPLADKLSGGDYDGDMAWVCWDPDIVENFT--NADM   990
AtRDR5      YVKA-LEEYVGNSKFAVFFPQKGPRSLGDEIAGGDFDGDMYFISRNPELLENFKPSEPWV   707
AtRDR3      YVKA-LEDYVGNAKFAVFFPQKGPRSLGDEIAGGDFDGDMYFISRNPKLLEHFKPSEPWV   718
AtRDR4      YVKS-LEQYVGNSKYGVFFPQKGPRSLGDEIAGGDFDGDMYFISRNPKLLEHYKPSEPWV   707
CeRRF3      WQPA-LA----HLVDVVVFPQHGPRPHPDEMAGSDLDGDEYSIIWDQEMLLDYN--EEAM   1157
CeEGO1      DIPE-LH----HMCDVVVFPQHGPRPHPDEMAGSDLDGDEYSIIWDQQLLLDKN--EDPY   997
CeRRF1      DIPE-LH----HMCDVVVFPQHGPRPHPDEMAGSDLDGDEYSVIWDQELLLERN--EEPF   970
CeRRF2      DIPE-LH----HLCDVVVFPQHGPRPHPDEMAGSDLDGDEYSVIWDQKLLLERN--EEAF   951
NcRRP3      IPPAGHPL--RELKNCIVFSQNGERDLPSQLSGGDLDGDTFNVIWDQSIVATLR-TFAAA   873
AtRDR6      DVPQ-LH----HMYDCLIFPQKGDRPHTNEASGSDLDGDLYFVAWDQKLIPPNRKSYPAM   888
AtRDR1      NVPA-LN----HMVDCVVFPQKGLRPHPNECSGSDLDGDIYFVCWDQELVPPRT--SEPM   820
AtRDR2      YEVH-FEE--KGYLDCIIFPQKGERPHPNECSGGDLDGDQFFVSWDEKIIPSEM--DPPM   853
ZmMOP1      YDPG-LDA--RGLIDCVVFPQRGERPHPNECSGGDLDGDLFFITWDDKLIPEKV--DAPM   847
SpRDP1      RCDE-LM----HLKNVIVFPTTGDRSIPAMCSGGDLDGDEYTVIWDQRLLPKIV-NYPPL   923
NcSAD1      DVPA-LR----HLRDVVVFPLTGDRDVPSMCSGGDLDGDDFFVIWDPLLIPKER-SHPPM   993
                            :.*   *        :*.* ***   :  :  ::
```

*rdr6-17*, G921D — RDR2 G886 — bridge helix

```
NcQDE1      PLEPDLSRYLKKDKTTFKQLMASHG-TGSAAKEQTTYDMIQKSFH--FALQPNFLGMCTN   1087
TtQDE1      PKEPDLSAYLGKDKTTFGELVRDTGRTGAAARHEAVYDMINKSFQ--FAMQPNYLGICTN   1048
AtRDR5      SLTPPSKSNSGRAPS--------------QLSPEELEEELFEMFLTAGFHASNVIGIAAD   753
AtRDR3      SSSKPSKIYCGRKPS--------------ELSEEELEEELFKMFLKARFCKRDVIGMAAD   764
AtRDR4      SSSPRSKIYTGRQPS--------------ELSPEQLEEELFKIFLKTGFSPSSVIGQAAD   753
CeRRF3      VFPSSS------AAE--------------E-DKEPTTDDMVEFFL--RYLQQDSIGRMSH   1194
CeEGO1      DFTSEK------QKA--------------SFKEDEIDDLMREFYV--KYLKLDSVGQISN   1035
CeRRF1      DFAVEK------IKV--------------PYDREKLDVLMREFYV--TYLKLDSVGQISN   1008
CeRRF2      DFAVEK------NLQ--------------TYEWEDIDDLMRDVYV--EYLKKDLVGLIAN   989
NcRRP3      DYPRVE-------PL--------------KLNREVESKDMADFFV--EFMKADHLGVIAV   910
AtRDR6      HYDAAE-------EK--------------SLGRAVNHQDIIDFFA--RNLANEQLGTICN   925
AtRDR1      DYTPEP-------TQ--------------ILDHDVTIEEVEEYFA--NYIVNDSLGIIAN   857
AtRDR2      DYAGSR-------PR--------------LMDHDVTLEEIHKFFV--DYMISDTLGVIST   890
ZmMOP1      DYTATR-------PR--------------IMDHAVTLEEIQKHFV--SYMINDTLGAIST   884
SpRDP1      LESSPK-----KSID--------------FLEGKPLIDSVKEFFV--NYIKYDSLGLISN   962
NcSAD1      ISEPIA-------GK--------------ELATEPTVNNLITFFV--LYMKYNNLPLIAH   1030
                                                   :   :        . :
```

bridge helix — RDR2 K923 — trigger loop

```
NcQDE1      YKERL----------CYINNSVSNKPAIILSSLVGNLVDQSKQGIVFNEASWA--QL---   1132
TtQDE1      YKERV----------CYHNNSVSDGVALLLSTLVGKLVDQSKQGILFDAASWD--RL---   1093
AtRDR5      SWLTIMDRFLILGDDRAEEKAEMKKKMLELIDIYYDALDAPKKGDKVYLPNKL-------   806
AtRDR3      CWLGIMDPFLTLGDESAKEKYERKKNILKLIDIYYDALDAPKKGAKVDLPPDL-------   817
AtRDR4      SWLAIMDRFLTLGDENVKEKAEMKKKMLKLTDIYYDAIDAPKTGTEVNLPLDV-------   806
CeRRF3      AHLA-----------YADLHGLFHENCHAIALKCAVAVDFPKSGVPAEPLSSF-------   1236
CeEGO1      SHLH-----------NSDQYGLNARVCMDLAKKNCQAVDFTKSGQPPDELERKWRKDEET   1084
CeRRF1      SHLH-----------NSDQYGLNSRVCMDLAKKNCQAVDFTKSGQPPDPLETKWRADPVT   1057
CeRRF2      SHLH-----------NSDQYGLTSRVCMNLAKKSCQAVDFSKSGKPPDELQTTWKTDDAT   1038
NcRRP3      RHMI---------LADERNEGTLDADCLKLAALHSKAVDFSKSGIHVDITELP-------   954
AtRDR6      AHVV---------HADRSEYGAMDEECLLLAELAATAVDFPKTGKIVSMPFHL-------   969
AtRDR1      AHTA---------FADKEPLKAFSDPCIELAKKFSTAVDFPKTGVAAVIPQHL-------   901
AtRDR2      AHLV---------HADRDPEKARSQKCLELANLHSRAVDFAKTGAPAEMPYAL-------   934
ZmMOP1      AHLI---------HADRDPLKARSPECVQLAALHSMAVDFAKTGAPAEMPLAL-------   928
SpRDP1      AWKAWAH------DHDNNPEGIFGNVCLELAEMHSKAVDFAKSGVACKMQAKY-------   1009
NcSAD1      AHLA---------TADAEVEGVKSPKCLELASLHSMAVDYVKTGVAAEFPRRL-------   1074
                                         :       :*  *  *
```

#### Figure S10. Amino acid alignment of 11 RDRα and 5 RDRγ proteins (page 10 of 14 pages)

Amino acid sequence alignment of 11 RDRα and 5 RDRγ, and conservation of key residues in the domains described in Figure 1 and 2. Domain annotation (colored lines above sequence) is based on the RDR2 structure in this study. *Neurospora crassa*, Nc; *Thielavia terrestris*, Tt; *Schizosaccharomyces pombe*, Sp; *Arabidopsis thaliana*, At; *Zea mays*, Zm; *Caenorhabditis elegans*, Ce.

trigger loop — Neck 3

```
NcQDE1    -------RRELL--------------------------------GGALSLPDPMYKSDSWL   1154
TtQDE1    -------RRERL--------------------------------GGRMSVEDPAYKGDVWA   1115
AtRDR5    -------KPDIF-PHYM----------------------------ERD--KKFQSTSIL   827
AtRDR3    -------EIKNF-PHYM----------------------------ERDPKRDFRSTSIL   840
AtRDR4    -------KVDLF-PHYM----------------------------ERN--KTFKSTSIL   827
CeRRF3    ------EQCEMT-PDYMM---------------------------S-GGKPMYYSTRLN   1260
CeEGO1    GEMIPPERAERV-PDYHM---------------------------GNDHTPMYVSPRLC   1115
CeRRF1    FEVIPPENPERI-PDFHM---------------------------GNERSPMYVSPRLC   1088
CeRRF2    GEMIPPERAERV-PDYHV---------------------------GSDHMPKYVSPRLC   1069
NcRRP3    -------RPPMYRPDFLVNGPDIKIHDKSTIDMEEQYLRQDDDDGDDTPRYKYYKSDKIL   1007
AtRDR6    -------KPKLY-PDFM----------------------------GKEDYQTYKSNKIL   992
AtRDR1    -------YVKEY-PDFM----------------------------EKPDKPTYESKNVI   924
AtRDR2    -------KPREF-PDFL----------------------------ERFEKPTYISESVF   957
ZmMOP1    -------RPREF-PDFM----------------------------ERWERPMYVSNGVL   951
SpRDP1    -------HPKRY-PDFM----------------------------QKTKTRSFRSETAV   1032
NcSAD1    -------DPKTW-PHFM----------------------------EKNRH-TYHSVTAL   1096
                                                          :  .
```

Neck 3 — *mop1-1*, Mu insertion between R956-A957 — QDE1ΔN homodimerization interface residues — Head

```
NcQDE1    GRGEPTHIIDYLKFSIARPAIDKELEAFHNAMKAA----KDTEDGAHFWDPDLAS-----   1205
TtQDE1    GAGEPRHIVDYLKFAVAKPTIDRELEELHKVMQASRRAGPDDDAAAHSWDPDLAV-----   1170
AtRDR5    GLIFD-----FVK-------------SQTTEEP--------SPSSEISKLPCFED-----   856
AtRDR3    GLIFD-----TVD-------------SHNAEEP--------PP-SEISKLWYFED-----   868
AtRDR4    GLIFD-----TVD-------------FHNAEDT--------TP-SGISKLQCFED-----   855
CeRRF3    GQLHR-----KAR-------------KVEEVLEE--FE-TRGSVFEREYDKLICP-----   1294
CeEGO1    GKLFR-----EFK-------------AIDDVLKI--SE-ERDEQVEISIDETIKI-----   1149
CeRRF1    GKLFR-----EFQ-------------AIDNVIKI--SE-ERDEQYNIELDETIFV-----   1122
CeRRF2    GKLFR-----EFQ-------------GIDNAMKI--SE-EKSEQYKIEVDESIRV-----   1103
NcRRP3    GRLFR-----AVD-------------EKKIWTKNIK---LEVPSGGVPFWKEVESSLLKR   1046
AtRDR6    GRLYR-----RVK-------------EVYDEDAEASSE-ESTDPSAIPYDAVLEI-----   1028
AtRDR1    GKLFR-----EVK-------------ERAPP-LISIKS-FTLDVASKSYDKDMEV-----   959
AtRDR2    GKLYR-----AVK-------------SSLAQ-RK-P---EAESEDTVAYDVTLEE-----   989
ZmMOP1    GKLYR-----AAL-------------RHEED-AEAL---LPAGPPSCVYDPDLEV-----   984
SpRDP1    GKIFR-----YAA-------------RFQRESGR-P-A-TYNPIMNTVYDPCMKL-----   1066
NcSAD1    GKLYD-----MVK-------------RETFDMK----E-NYQLPFDNRILKHTKC-----   1128
          *
```

QDE1ΔN homodimerization interface residues — *rdr6-14*, W1039stop — Head

```
NcQDE1    ------------YYTFFKEISDKSRSSALLFTTLKNRIGEVEKEYGR-------------   1240
TtQDE1    ------------YFENFKALTAESRSLRAVLEALQNALGAVEHEWKV-------------   1205
AtRDR5    -E-----PVSEFHMQKCR----------LWYDNYRTEMTQAMKTDK--------------   886
AtRDR3    -E-----PVPKSHMDKFT----------SWYENYRSEMSQAMMETDK-------------   899
AtRDR4    -E-----PVSEFDMEKCK----------LWHKDYRKEMCQAMNSDD--------------   885
CeRRF3    -E-----DVD-VFFGNEIKLVQT----LTLRDEYVDRMQQLLDEYGI-----EDEASVVS   1338
CeEGO1    -D-----GYT-EYMASAK----------NDLARYNAQLRSMMENYGI-----KTEGEVFS   1187
CeRRF1    -T-----GFE-RYMESAQ----------KQLSSYNGQLRSIMENYGI-----RSEGEIMS   1160
CeRRF2    -D-----GFE-EYMEDAK----------KQLASYNGQLKSTMDTYGI-----QSEGEIMS   1141
NcRRP3    VRGIGQVQWQ-HRLDEAR----------RICESYEDGIKDAMVEFADSPTQPLKELEVVM   1095
AtRDR6    -P-----GFE-DLIPEAW----------GHKCLYDGQLIGLLGQYKV-----QKEEEIVT   1066
AtRDR1    -D-----GFE-EYVDEAF----------YQKANYDFKLGNLMDYYGI-----KTEAEILS   997
AtRDR2    -A-----GFE-SFIETAK----------AHRDMYGEKLTSLMIYYGA-----ANEEEILT   1027
ZmMOP1    -A-----GFD-EFLDAAE----------ERYEAYAERLGALMTYYSA-----EREDEILT   1022
SpRDP1    -P-----RFKTEYLNVAE----------EVKKHYDNDLRSIMARFDI-----STEYEVYT   1105
NcSAD1    -R-----ALRDGTLAKAR----------RIKSQYDTAMRRVMCQLEI-----ATEFEVWT   1167
                                   :
```

### Figure S10. Amino acid alignment of 11 RDRα and 5 RDRγ proteins (page 11 of 14 pages)

Amino acid sequence alignment of 11 RDRα and 5 RDRγ, and conservation of key residues in the domains described in Figure 1 and 2. Domain annotation (colored lines above sequence) is based on the RDR2 structure in this study. *Neurospora crassa*, Nc; *Thielavia terrestris*, Tt; *Schizosaccharomyces pombe*, Sp; *Arabidopsis thaliana*, At; *Zea mays*, Zm; *Caenorhabditis elegans*, Ce.

```
                                                                            Head
NcQDE1     --LVKNKEMR---DS---------KDPYPVRVNQVYEKW---------------CAITPEA    1272
TtQDE1     --LMSKGS-----SS---------SLTYPEKVRQLHAKW---------------CAIEPRA    1235
AtRDR5     --------------DE---------------SANEVIQRYKQEFYGAAG-FE----------    908
AtRDR3     ---------VK-RNQ---------------LTNEVIQRYKQDFYGAAG-FE----------    924
AtRDR4     --------------DD---------------SCNEVIQKYKQEFYSAAG-FK----------    907
CeRRF3     GHAASI-KRLAGMERDDYSFYHTDKVVELRYEKLYAVFRAKFFEEFGGEEINI-------   1390
CeEGO1     GCIVDMRNRISDKDQDDMSFFNTNQMIETKLTNLFKKYREIFFEEFEGGWEGNTEAFSRY   1247
CeRRF1     GCIVEMRNRISDKDQDDMSFYNTNQMIETKMTSLVCKFRETFFEEFGGFTV-KCTLLPNA   1219
CeRRF2     GCIIEMRNRISDSDQDDMSFYNTNRMIETKMTALVSKFRTIFFQQFGGFQE-VCTLLPDA   1200
NcRRP3     GFILNKKGIQSRRQRD--KSSKLSD----AFARITKMVTNVLRPSTP-------------   1136
AtRDR6     GHIWSMPK-YT---SK--KQGELKERLKHSYNSLKKEFRKVFEETIP-------------   1107
AtRDR1     GGIMRMSKSFTKRR--------DAESIGRAVRALRKETLSLFNASEE-------------   1036
AtRDR2     GILKTKEM-YLARDNR--RYGDMKDRITLSVKDLHKEAMGWFEKSCE-------------   1071
ZmMOP1     GNIRNKLV-YLRRDNK--RYFEMKDRIIAAVDALHAEVRGWLRACKE-------------   1066
SpRDP1     AFILFKDDLAK---TV--NEYGLREEVSFQFDLLKKKYTQEYLEKCA-------------   1147
NcSAD1     AFVMSKPRVGS--------DYKLQDNVGRESSALKQH----FKDQCK-------------   1202
                                               :
```

QDE1ΔN homodimerization interface residues — *rdr2-4*, W1083stop

```
                                                                            Head
NcQDE1     MDKSGAN-YD---------SKVIRLLELSFLADRE---------MNTWALLRASTAFKLY   1313
TtQDE1     VGRAGSNRLD---------AKTAALLEQPFLADRGG------AGTSYWALLRASTAFKAY   1280
AtRDR5     --DSKK-------SLEELYPQALALYKIVYDYAIHA------------------------    935
AtRDR3     --DSNK-------SLEELYPQALALYNVVYDYAIQE------------------------    951
AtRDR4     --ESKK-------ILEELYPKALALYNVT-------------------------------    927
CeRRF3     ENDGKNTRLKCT---KAMHEKIRQWYFVAYVQPKIN---------------------KAG   1426
CeEGO1     GRDSNILQRQCRAPTVQMMKKAVAWYRACYEEARIT---------------------REN   1286
CeRRF1     YDNGNCLNYRCEDPDQEVRKKAVAWYRACYECAQST---------------------REV   1258
CeRRF2     YNESNFFNFRCENPNEEIRKKAVAWYRACYECAKST---------------------REP   1239
NcRRP3     --PEEATS------ELHALELCLACFYVAGEKKS-------QPQ-ESW--------KRQI   1172
AtRDR6     --DHENLS--EEEKNILYEKKASAWYHVTYHPEWVKKSLELQDP------------DESS   1151
AtRDR1     --EE------------NESAKASAWYHVTYHSSYWGLYN------------------EGLN   1065
AtRDR2     --DEQ-----------QKKKLASAWYYVTYNPNHRD-----------------------E   1095
ZmMOP1     --D-------------DASRVASAWYHVTYHPDRRG-----------------------E   1088
SpRDP1     --LSNQSAFDSSEYEERINSAVAATYDVTYDQRV-----------KSV--------GNGT   1186
NcSAD1     --KEAGG---------DLLSFVSAMYRVTYEEVRIALFEAKQPHVRPDGRLGTR-KITPK   1250
```

QDE1ΔN homodimerization interface residues

```
                                                                            Head
NcQDE1     YHKSPKFVWQMAGRQLAYIKAQMTSRPG------E-GAPALMTAFMYA------------   1354
TtQDE1     YKTNPKFVWQMAGAQLAFIKAQMSSPGGGGGGGSD-GMPLLVTPLMYA------------   1327
AtRDR5     GVSKCRFVWKVAGPVLCRFYLNKKMQEKCLVCAPSVLKELWG------------------    977
AtRDR3     GVAKCTFAWNVAGPVLCKFYLKKTK-DKSVVASTSVLKKLLG------------------    992
AtRDR4     ------------------------------------------------------------    927
CeRRF3     RCIGQSLPWVAWDAL-CDLRRQLMLDKNDA--------VLRGKYPIAARLEEEIENSIER   1477
CeEGO1     K--KLSFAWLAYDVI-AKVKQDKSLTSDEV--------KMGGANPLYTMLDDHRSQYLVD   1335
CeRRF1     R--KLSFAWIAYDVI-AKVKETNVLNNERM--------QIGGANPMYTFLEEHRKQYLID   1307
CeRRF2     R--KLSFAWIAYDVI-AKIKETKVLENVEM--------NIGGANPMYTYLEHHRARYIND   1288
NcRRP3     ATGLESFRLVAGSALLLEIKAQEQKIR--L----------------------RHA--A-   1204
AtRDR6     HAAMLSFAWIAADYL-ARIKIRSREMG-SID-SAK-PVDSLAKF--LAQRL----------   1196
AtRDR1     RDHFLSFAWCVYDKL-VRIKKTNLGRR-QRQETLE-RLDHVLRF--G--------------   1107
AtRDR2     KLTFLSFPWIVGDVL-LDIKAENAQRQ-SVEEKTS-GLVSI-------------------   1133
ZmMOP1     K-RFWSFPWIICDTL-LAIKAARRCRK-RVEDAAV-PMDCDGS-----------------   1127
SpRDP1     TEVLISFPYLFSSRL-CQLSRKAMLTANNF------------------------------   1215
NcSAD1     TMPLVSFPWLFWDKL-GELARAGAVLQRRLDDGSE-DMDLLSDVPLVSQRRRGKHN--AS   1306
```

**Figure S10. Amino acid alignment of 11 RDRα and 5 RDRγ proteins (page 12 of 14 pages)**

Amino acid sequence alignment of 11 RDRα and 5 RDRγ, and conservation of key residues in the domains described in Figure 1 and 2. Domain annotation (colored lines above sequence) is based on the RDR2 structure in this study. *Neurospora crassa*, Nc; *Thielavia terrestris*, Tt; *Schizosaccharomyces pombe*, Sp; *Arabidopsis thaliana*, At; *Zea mays*, Zm; *Caenorhabditis elegans*, Ce.

```
                                                   QDE1ΔN homodimerization interface residues
NcQDE1     --------------------------------GLMPDKKFTKQYVARL---------EGD   1373
TtQDE1     --------------------------------GLAPDGRFVKQYLARL---------ECD   1346
AtRDR5     ------------------------------------------------------------   977
AtRDR3     ------------------------------------------------------------   992
AtRDR4     ------------------------------------------------------------   927
CeRRF3     Q---FDKFLKLKDLI----ESHKDALFLRRYVYFYGDQIIKMLFILKVWLERENVLPSSV   1530
CeEGO1     NSRKFEAFRQFSTPKTSGEQVKRAHRIIKMYTETYP-GLDAVLFMLDEWARISNLFENQS   1394
CeRRF1     HDADFKNFC-ELDHLITGEKSKEAISILKIYLEMIP-GLDSVFFMLMRWGESLRLFDGKP   1365
CeRRF2     HVDDFEKFRRNLDNLITGKESKKAVFILQRYIYMIP-GLDSVMFVLMRWGEALELFEGRS   1347
NcRRP3     RSGGFVG----------------------------------------VRGGSRVAGRGGR   1224
AtRDR6     ------------------------------------------------------------   1196
AtRDR1     ------------------------------------------------------------   1107
AtRDR2     ------------------------------------------------------------   1133
ZmMOP1     ------------------------------------------------------------   1127
SpRDP1     ------------------------------------------------------------   1215
NcSAD1     GSSDFMD----------------------------------------EHGDPLSYTRTSD   1326

NcQDE1     GSEYPDPEVYEVLGDDDFDGIGFTGNGDY-------------------------------   1402
TtQDE1     GSQYPDDDLEG----DGEDGGGGGGRGDDVD-----------------------------   1373
AtRDR5     ------------------------------------------------------------   977
AtRDR3     ------------------------------------------------------------   992
AtRDR4     ------------------------------------------------------------   927
CeRRF3     LSIWQLGRLLIRLGLGDLLGNPTIDYE-------KSLLMPTTMF-QQWISKKEDADEAP-   1581
CeEGO1     LREYHLSLLFILFATRQFSSVDGN----------AAKFFNKVDE-KSYKQSKTIGDFEPS   1443
CeRRF1     IKIYHFFLMFILFATRQLASADGN----------AEPFFKIIEK-EEYEKQKRDSSRGNI   1414
CeRRF2     MSIYHFLLMFIMFSTGQLASADGN----------AEQFFGKLDV-EEYRRRKKNGTLEPD   1396
NcRRP3     -------------------GGHRQPARVDVNTAAEAATLQTADTGAAFGLGSPSSTSAPT   1265
AtRDR6     ------------------------------------------------------------   1196
AtRDR1     ------------------------------------------------------------   1107
AtRDR2     ------------------------------------------------------------   1133
ZmMOP1     ------------------------------------------------------------   1127
SpRDP1     ------------------------------------------------------------   1215
NcSAD1     GKIIHYGQILNLFSHDDEDGDERNDAR-DRNSSSE------------------DSTHVSN   1367

NcQDE1     ------------------------------------------------------------   1402
TtQDE1     ------------------------------------------------------------   1373
AtRDR5     ------------------------------------------------------------   977
AtRDR3     ------------------------------------------------------------   992
AtRDR4     ------------------------------------------------------------   927
CeRRF3     -ILRNFDMGTMMLEFLRYLASQSFASAESISLRVFYEKDIVE---------PILTKSAQW   1631
CeEGO1     LYIEEKGKSQMMVKFLEFLASRKFRKMANLSFCALDFS--------------SIFMRGEW   1489
CeRRF1     DPLTEKKRSDMMVKFFQFMGCRKFRKMSTLSFCPLNFS--------------SIFMRGEW   1460
CeRRF2     IPLTRHQRSDMMFKFFQLMGSRKFRTMRSISFRPLGFS--------------SVFMSGEW   1442
NcRRP3     ------------------------TPSSGSNYTPSSGSDMRDGIFNSPTG--------PY   1293
AtRDR6     ------------------------------------------------------------   1196
AtRDR1     ------------------------------------------------------------   1107
AtRDR2     ------------------------------------------------------------   1133
ZmMOP1     ------------------------------------------------------------   1127
SpRDP1     ------------------------------------------------------------   1215
NcSAD1     ------------------------SSKSSSNLSPVAEEDLL--TFESPPASPGVTPASPQ   1401
```

#### Figure S10. Amino acid alignment of 11 RDRα and 5 RDRγ proteins (page 13 of 14 pages)

Amino acid sequence alignment of 11 RDRα and 5 RDRγ, and conservation of key residues in the domains described in Figure 1 and 2. Domain annotation (colored lines above sequence) is based on the RDR2 structure in this study. *Neurospora crassa*, Nc; *Thielavia terrestris*, Tt; *Schizosaccharomyces pombe*, Sp; *Arabidopsis thaliana*, At; *Zea mays*, Zm; *Caenorhabditis elegans*, Ce.

```
NcQDE1      ------------------------------------------------------------   1402
TtQDE1      ------------------------------------------------------------   1373
AtRDR5      ------------------------------------------------------------   977
AtRDR3      ------------------------------------------------------------   992
AtRDR4      ------------------------------------------------------------   927
CeRRF3      M-------PLH--LIAYRTFHSIAVSGRFDALHLDDEDAVD-------------------   1663
CeEGO1      Q-------IFH--LAALKTYYNVLFNLRFEELPVSTDPTTT-------------------   1521
CeRRF1      R-------IFH--ESALKTYYNILFNLRFEELPVSSDPTIT-------------------   1492
CeRRF2      M-------VFH--EPALKTYYNIMFNLRFEELPISTDPTTT-------------------   1474
NcRRP3      ---AGYQPELVHAQAGARTAG---------------------------------------   1311
AtRDR6      ------------------------------------------------------------   1196
AtRDR1      ------------------------------------------------------------   1107
AtRDR2      ------------------------------------------------------------   1133
ZmMOP1      ------------------------------------------------------------   1127
SpRDP1      ------------------------------------------------------------   1215
NcSAD1      VDLLGPTLTLTKAELAIGLAGP--VATTYTPPPTDEETVICHGHSNRASHLSASSSSLDP   1459
```

```
NcQDE1      ------------------------------------------------------------   1402
TtQDE1      ------------------------------------------------------------   1373
AtRDR5      ------------------------------------------------------------   977
AtRDR3      ------------------------------------------------------------   992
AtRDR4      ------------------------------------------------------------   927
CeRRF3      ---Q--ITESKDPILVNESLFS-----SRNYNDDYPISRSRILQSLKDWSG------VKE   1707
CeEGO1      ---VRSIIRENEPFVIELP---------------ANCDRSLVHRKLVEHTG------VKE   1557
CeRRF1      ---AETMDRECEPFVIELP---------------ENINVNDLINNMKKHTN------VST   1528
CeRRF2      ---HETRLRECEPFVIELP---------------EKANMDMLIKKLTEKSK------VSQ   1510
NcRRP3      -------------------------TSGGQVPSAAPANMPVRLAPADKVAHLAALYAQMQ   1346
AtRDR6      ------------------------------------------------------------   1196
AtRDR1      ------------------------------------------------------------   1107
AtRDR2      ------------------------------------------------------------   1133
ZmMOP1      ------------------------------------------------------------   1127
SpRDP1      ------------------------------------------------------------   1215
NcSAD1      QPLIEETSGENEDLLLDIYSASPPRADGARLASAVGSDVPVKGGPPP-------VWMEQV   1512
```

```
NcQDE1      ------------------------------------------------------------   1402
TtQDE1      ------------------------------------------------------------   1373
AtRDR5      ------------------------------------------------------------   977
AtRDR3      ------------------------------------------------------------   992
AtRDR4      ------------------------------------------------------------   927
CeRRF3      IIPREIT-------GTRKSDMIYVTS----------V-GTVLARQRLARLL----LL---   1742
CeEGO1      IFMRNMEKSVRSSDDVQKINMRLLVS----------TRGTLESMYKLRQLV----AVKVP   1603
CeRRF1      VKMRRQEKNPINDKAKPKTTVRYIVS----------VSGTLESIQMLKKLS----AVTIP   1574
CeRRF2      LNLRRQDKNK----KTSNSTVRYIVS----------ASGTFESTQQLRKLV----TVTSP   1552
NcRRP3      IYQRH-------------------------------------------------------   1351
AtRDR6      ------------------------------------------------------------   1196
AtRDR1      ------------------------------------------------------------   1107
AtRDR2      ------------------------------------------------------------   1133
ZmMOP1      ------------------------------------------------------------   1127
SpRDP1      ------------------------------------------------------------   1215
NcSAD1      IRMGHRI-------PTPPLDSIAIPNVIGVPQFESPLTGTVVASAIQDPFVSPSPAVATP   1565
```

**Figure S10. Amino acid alignment of 11 RDRα and 5 RDRγ proteins (page 14 of 14 pages)**

Amino acid sequence alignment of 11 RDRα and 5 RDRγ, and conservation of key residues in the domains described in Figure 1 and 2. Domain annotation (colored lines above sequence) is based on the RDR2 structure in this study. *Neurospora crassa*, Nc; *Thielavia terrestris*, Tt; *Schizosaccharomyces pombe*, Sp; *Arabidopsis thaliana*, At; *Zea mays*, Zm; *Caenorhabditis elegans*, Ce.

```
NcQDE1      ------------------------------------------------------------   1402
TtQDE1      ------------------------------------------------------------   1373
AtRDR5      ------------------------------------------------------------   977
AtRDR3      ------------------------------------------------------------   992
AtRDR4      ------------------------------------------------------------   927
CeRRF3      ---------SGETIRDAIANNVVPNEVRDEFL----------------------------   1765
CeEGO1      ----IKTYVTGQDVSTQMARLCYEKIVRGHINI---------------------------   1632
CeRRF1      ----IKSHWEGEEVSQQMASLCYQKVMNGEF-----------------------------   1601
CeRRF2      ----IKNHWEGQDLAQQMANFCYLKIMKDDL-----------------------------   1579
NcRRP3      ------------------------------------------------------------   1351
AtRDR6      ------------------------------------------------------------   1196
AtRDR1      ------------------------------------------------------------   1107
AtRDR2      ------------------------------------------------------------   1133
ZmMOP1      ------------------------------------------------------------   1127
SpRDP1      ------------------------------------------------------------   1215
NcSAD1      ATSGSGGGWGGGGGGGGGGYIGIGGPINKELVMGLDGGIKEEDREETDEEAEEVELEIDE   1625

NcQDE1      -------------1402
TtQDE1      -------------1373
AtRDR5      -------------977
AtRDR3      -------------992
AtRDR4      -------------927
CeRRF3      -------------1765
CeEGO1      -------------1632
CeRRF1      -------------1601
CeRRF2      -------------1579
NcRRP3      -------------1351
AtRDR6      -------------1196
AtRDR1      -------------1107
AtRDR2      -------------1133
ZmMOP1      -------------1127
SpRDP1      -------------1215
NcSAD1      DPIAVRYAQMAAL1638
```

```
              Metal A site            Metal B site
ScRpb1     477-NADFDGD-487          834-NQEDSM-839
Tt_ β′     735-NADFDGD-745          683-NFEDAI-688
NcQDE1    1003-GGDYDGD-1013         707-MNDGVG-712
RDR2       828-GGDLDGD-834          553-FSDGIG-558
```

Bridge helix AA alignment

```
                                                BH-HN                BH-HC
ScRpb1_809-846       ------------TPQEFFFHAMGGREGLIDTAVKTAETGYIQRRLVKALE  846
Tt_β′_1067-1103      -------------VLEYFISSHGARKGGADTALRTADSGYLTRKLVDVTH  1103
RDR2_868-900         ---------TLEEIHKFFVDYMISDTLGVISTAH---LVHADRDP-----  900
NcQDE1_1058-1098     TGSAAKEQTTYDMIQKSFHFALQPNFLGMCTNYKE-RLCYINNS------  1098
                                  : *            :  :      :  .
```

Trigger loop AA alignment

```
Tt_β′_1219-1268      ------------EAVGIVAAQS-IGEPGTQL----TMRTFHTGGVAGAADITQGLPRVIELFEARRP 1268
ScRpb1_1063-1108     -------------MVGVLAAQS-IGEPATQM----TLNTFHFAGVASK-KVTSGVPRLKEILNVA-- 1108
RDR2_901-939         EKARSQKCLELANLHSRAVDFAKTGAP-AEMPYALKPREF--------------------------- 939
NcQDE1_1099-1140     --VSNKPAIILSSLVGNLVDQSKQGIVFNEASWAQLRRELLG-GA---------------------- 1140
                                  .  .  :  *     :        . :
```

**Figure S11. Amino acid alignments of catalytic domains of RDR2, NcQDE-1 and multi-subunit RNAPs (related to Figure 2)**

Amino acid sequence alignment of the active site (metal A and B), bridge helix and trigger loop domains among *Arabidopsis thaliana* RDR2, *Saccharomyces cerevisiae* (Sc) Pol II, bacterial (*Thermus thermophilus*, Tt) RNAP, and *Neurospora crassa* (Nc) QDE-1. Amino acid residues discussed in the text are highlighted in yellow.

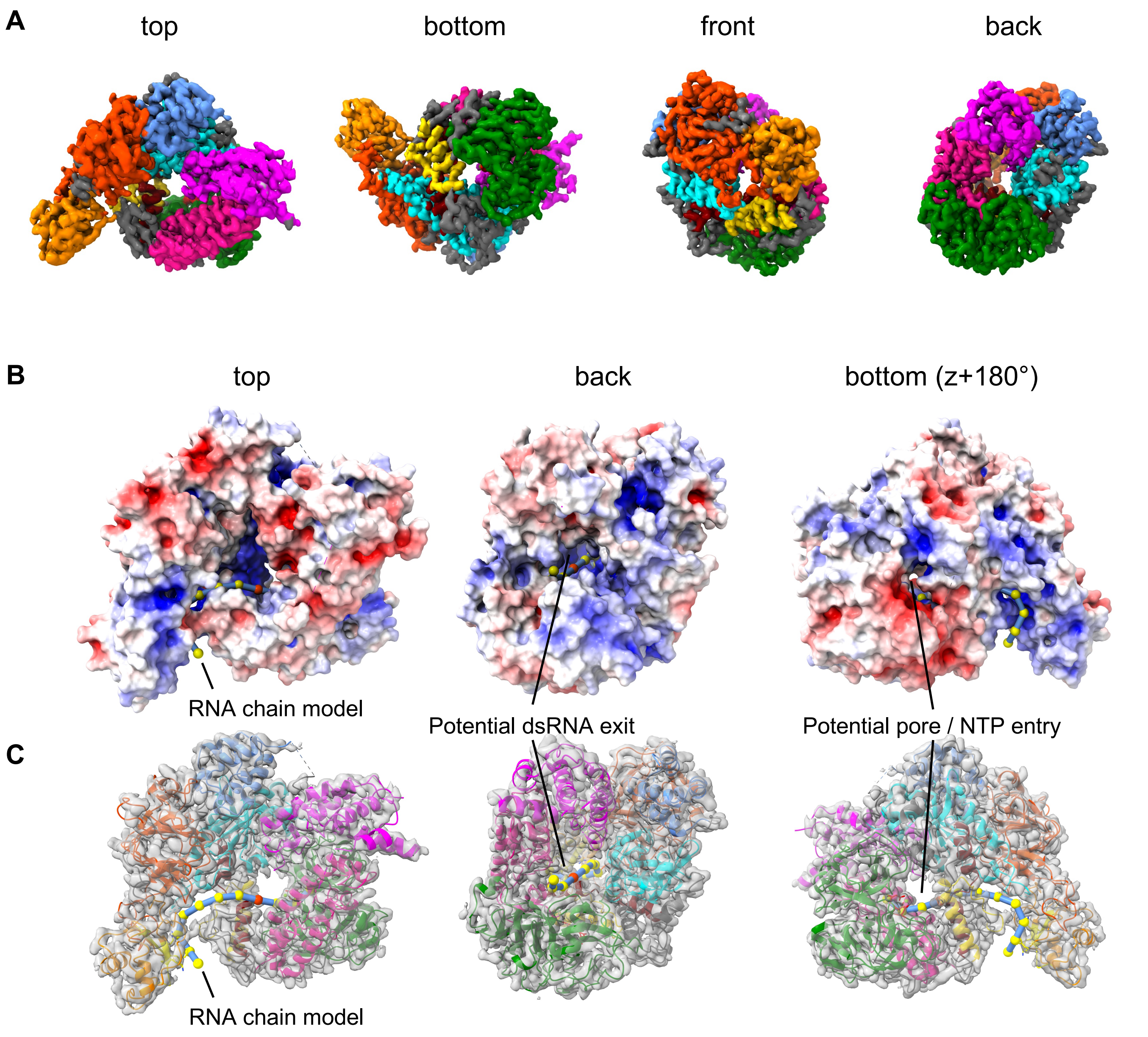

**Figure S12. Alternative views of RDR2 and a proposed RNA path (related to Figure 3)**
(A) Top, bottom, front, and back views of RDR2. Domains are colored as in Figure 1. (B) Top, back and bottom views of RDR2 colored according to the electrostatic surface potential with negative, neutral, and positive charges shown in red, white, and blue, respectively. RNA is depicted as beads on a cyan-colored string, with beads depicting individual bases. The red bead depicts the position that basepairs with the initiating nucleotide of the complementary strand (position +1). (C) Top, back and bottom views of RDR2 with the predicted dsRNA exit channel and NTP entry pores highlighted.

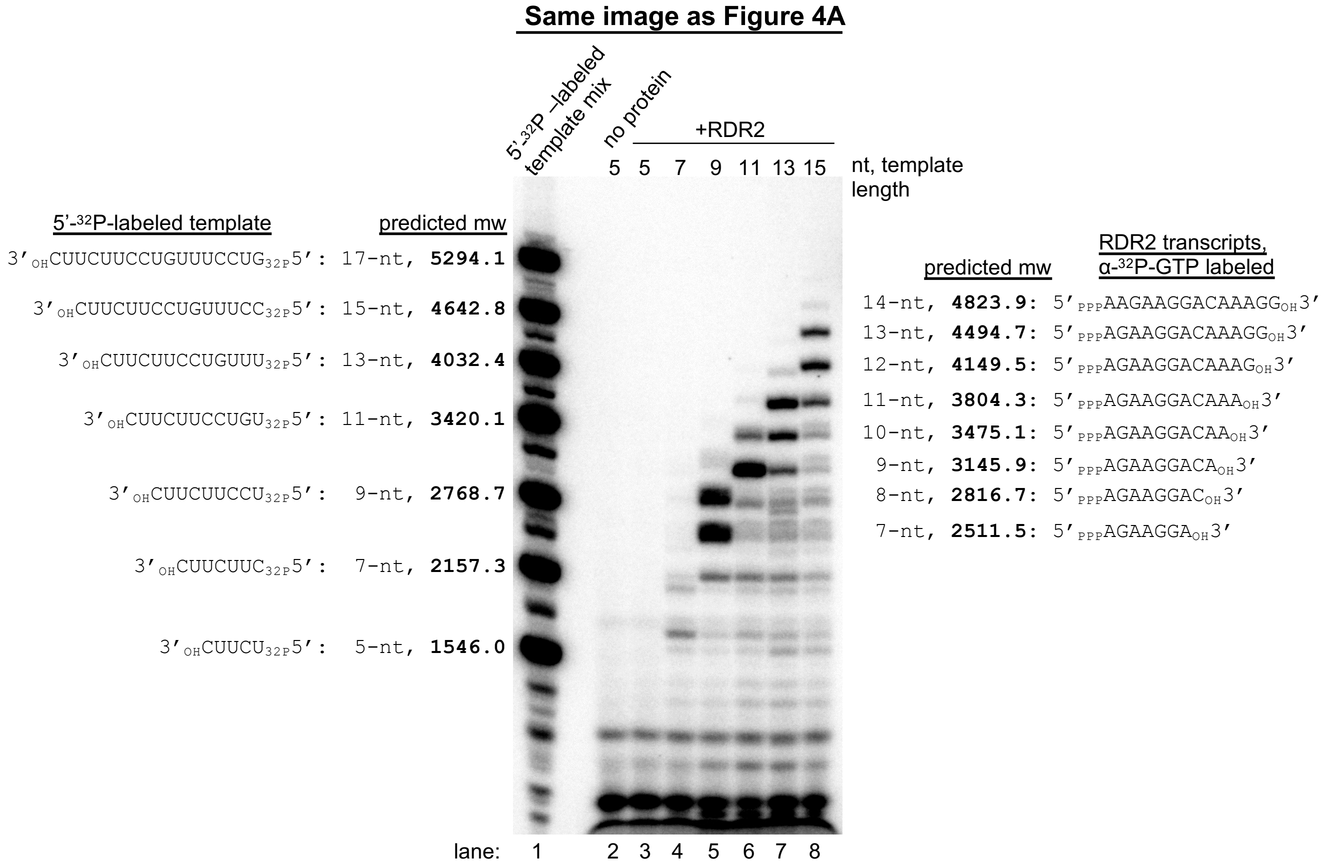

**Figure S13. Molecular masses of template RNAs (pyrimidine-rich) and expected RDR2 product RNAs (purine-rich) to explain the different mobilities of template and product RNAs in Figure 4A**

For template and product bands observed in Figure 4A, the known or predicted RNA sequences and their masses (mw) are shown. Purine-rich transcripts have a higher average mass per nucleotide than the pyrimidine-rich temp[late RNAs.

#### A. Known missense mutations in RDR2 or its paralog, RDR6 mapped onto the RDR2 structure

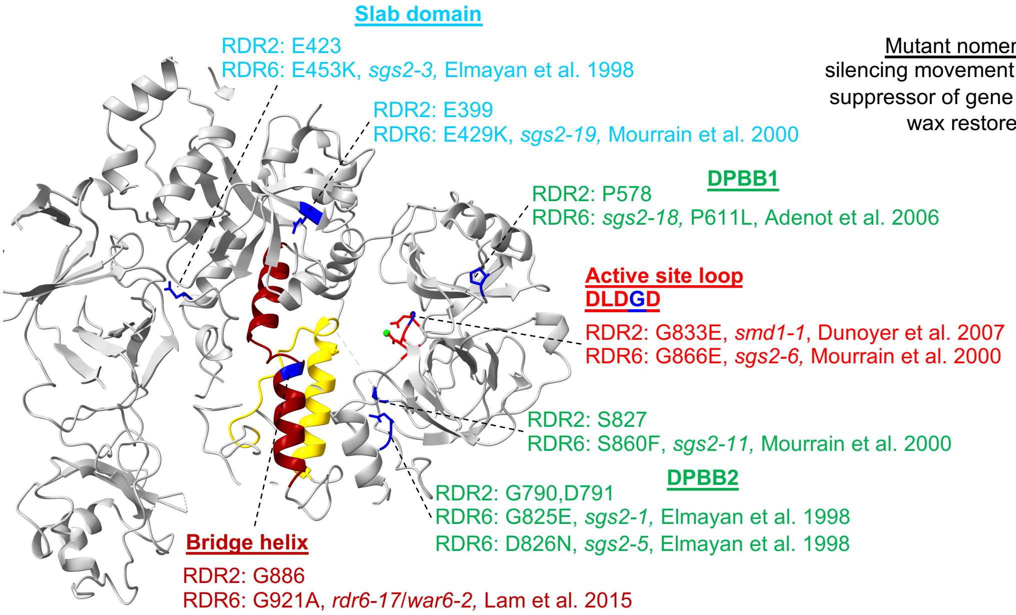

#### B. Known nonsense mutations in the head domain of RDR2, RDR6 or Mop1

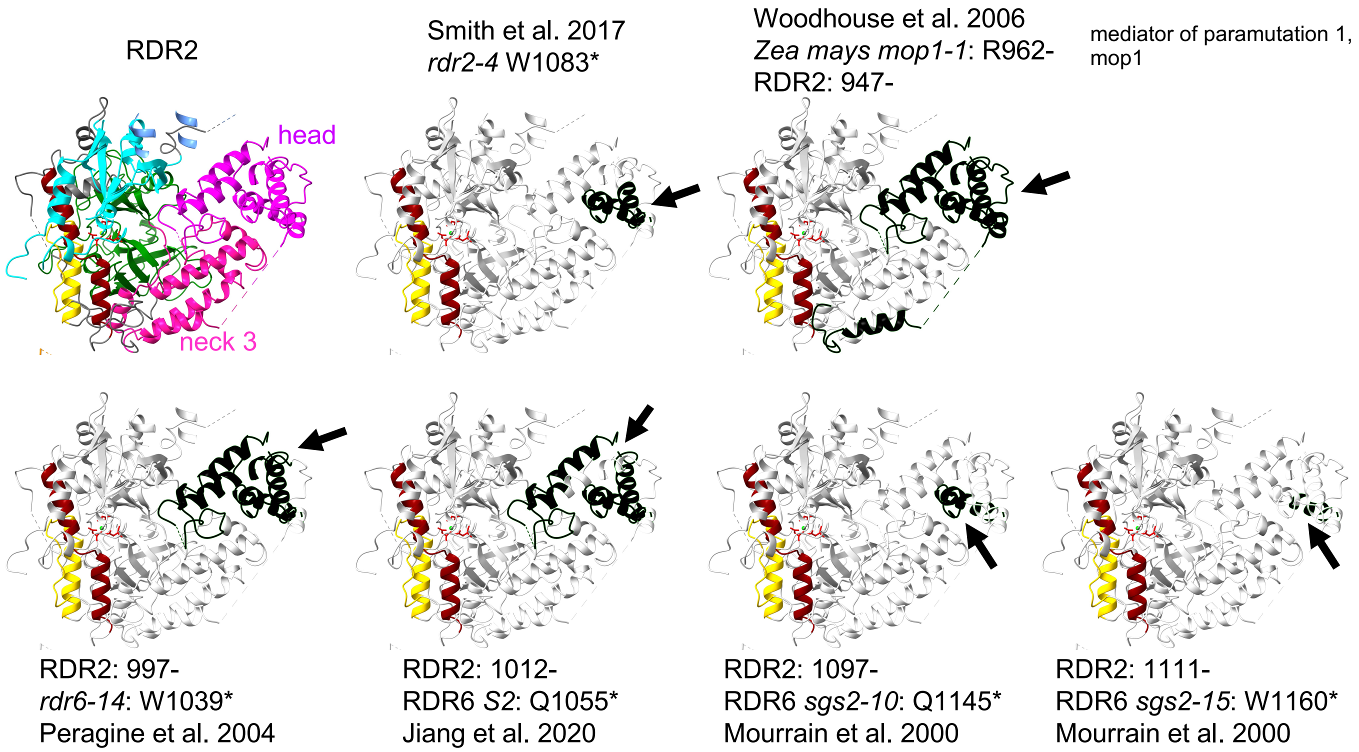

**Figure S14. Known mutations of RDRs superimposed onto RDR2**
(A) Missense mutations found in RDR2 or its paralog, RDR6 are mapped onto the corresponding residues of the RDR2 structure. A missense mutation found in RDR2, G833E maps to the active site loop. Mutations mapped onto slab domain are colored in blue; those in DPBB are in green; those in the active site are in light red, and those in the bridge helix in orange red. (B) Nonsense/insertion mutations found in RDR2, RDR6 or Mop1 (the RDR2 homolog in *Zea mays*) were mapped onto the corresponding residues of RDR2. These mutations result in truncation of the head domain. Truncated portions of the head domain are colored in black, with truncation positions indicated by black arrows. Other domains are colored as in Figure 1C.

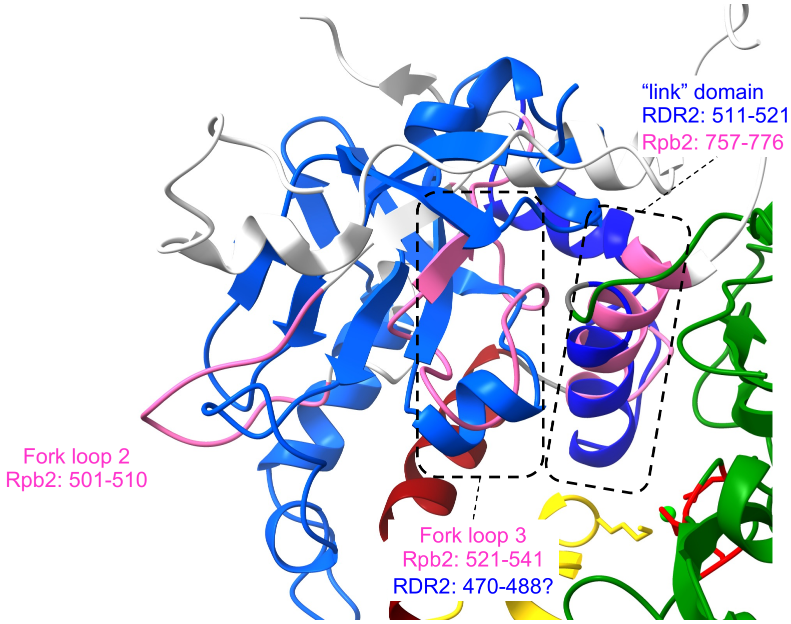

**Figure S15. Comparison between the RDR2-slab and Pol II-fork loops**
The RDR2 slab domain (blue) was superimposed onto the yeast Pol II (PDB: 2E2I) fork loops (white). Fork loop 2, fork loop 3 and the link domain helix are colored in magenta. The superimposed link and fork loop 3 domains are indicated by broken rectangles.

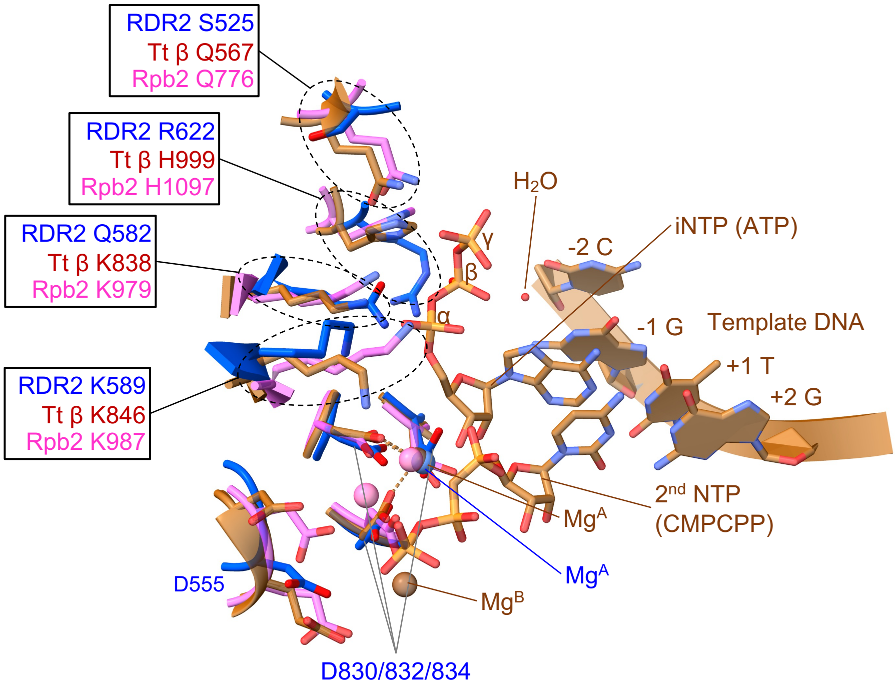

**Figure S16. Structural comparisons suggesting a structural basis for RDR2 internal initiation**
Potential initiating NTP (iNTP) interacting residues in RDR2. The three aspartates of the RDR2 active site and four residues that correspond to iNTP-interacting residues that are invariant in multi-subunit RNAPs (yeast Pol II, 2E2H and bacterial *Thermus thermophilus* RNAP, 4Q4Z) are superimposed, as described in Figure 2A. RDR2 is colored in blue; yeast Pol II in magenta; and bacterial RNAP in brown.
